# A molecular glue degrader-antibody conjugate for HER2-targeted cyclin K degradation

**DOI:** 10.64898/2026.11.12.760175

**Authors:** Sofia A. Srdanovic, Alina Chrzastek, Laura C. Helbling, Pauline Franz, Aidan B. Pidd, Damjan Lalović, Lea N.C. Rochet, Amy McIntosh, Janine L. Gray, Jack W. Houghton, Jianan Lu, Josephine Walton, Thomas Mills, Vanessa Stöcker, Nicolas Veillard, Paul W. Hogg, Zuzanna Kozicka, Edward W. Tate

## Abstract

Cyclin-dependent kinase 12 (CDK12) is an attractive therapeutic target due to its critical role in transcriptional regulation and DNA damage response and is frequently co-amplified with *ERBB2* (HER2) in breast cancer. CDK12 forms an active complex with its obligate partner cyclin K (CCNK), and pharmacological degradation of CCNK has emerged as a promising strategy for disrupting CDK12-dependent transcriptional programs. However, the broad expression and essential functions of the CDK12/CCNK complex may limit the therapeutic window of highly potent molecular glue degraders.

Here, we investigate whether antibody-mediated delivery can impart cellular selectivity to a molecular glue degrader through targeting the HER2 receptor. Molecular glues lack the modular architecture of PROTACs, thus linker placement can be the central design problem. We designed SAS1001, a CCNK molecular glue degrader with a solvent-exposed piperazine primed for linker attachment. SAS1001 induced potent and ubiquitin-proteasome system-dependent degradation of CCNK and inhibited proliferation in a range of breast cancer cells. Conjugation to trastuzumab (Tz) generated the HER2-targeted degrader-antibody conjugate Tz-SS2009, which internalized preferentially in HER2-positive cells and produced enhanced CCNK degradation and antiproliferative effects, relative both to HER2-negative cells and to an isotype-matched non-targeting conjugate. Whole proteome analysis further demonstrated CCNK depletion and revealed downstream changes consistent with established roles of the CDK12/CCNK complex in transcriptional regulation.

These findings demonstrate that antibody-mediated delivery can redistribute the cellular activity of a potent molecular glue degrader towards antigen-positive cells. In the CR8/DS18 purine series, substitution at a solvent-exposed position preserved CCNK degradation when the degrader was delivered on a full-length IgG. More broadly, this work extends antibody-directed degradation beyond the CRBN-based glues that have dominated the field and illustrates how structural characterization of a glue scaffold can identify a linker attachment point that does not compromise activity.

## Introduction

Cyclin-dependent kinase 12 (CDK12) is a transcriptional serine/threonine kinase in the cyclin-dependent kinase (CDK) family that forms an active complex with its regulatory partner, cyclin K (CCNK).^1^ Through regulation of RNA polymerase II (RNA Pol II) transcription and RNA processing, the CDK12/CCNK complex contributes to the maintenance of gene expression programs and genomic stability.^2,3^ This includes regulation of essential genes involved in the DNA damage response (DDR) pathway like *BRCA1/2* and Fanconi anemia family members.^4,5^

While CDK12 is broadly expressed in mammalian tissue,^6^ dysregulation is common in cancer and has been associated with increased invasiveness and poorer survival outcomes.^7–9^ CDK12 inhibition has been shown to reduce the expression of a subset of DDR genes, including *BRCA1* and *ATR*, compromising maintenance of the genome and sensitizing cells to DNA-damaging agents.^10,11^ These observations have motivated efforts to target the CDK12/CCNK complex pharmacologically.^12–22^ Selective kinase inhibition has proven difficult, as the serine/threonine kinase domain is highly conserved across the CDK family.^3^ An alternative strategy to overcome the limitations of inhibition is targeted protein degradation. In this approach, a small molecule induces or remodels protein–protein interactions, directing a target protein to the ubiquitin–proteasome system (UPS) for degradation. Selectivity can therefore arise from these interfaces rather than from discrimination between conserved active sites. Within the CDK12/CCNK complex, the cyclin is an attractive target given that CDK12 and its paralog CDK13 (which shares over 93% sequence identity in the kinase domain) both depend on CCNK, so removal prevents formation of both active complexes which have been implicated in disease.^3,23^

The discovery that the pan-CDK inhibitor CR8 (Figure 1a) induces selective proteasomal degradation of CCNK by Słabicki *et al.* provided the first example of this approach for CDK12/CCNK.^16^ CR8 acts as a molecular glue between the CDK12/CCNK complex and DDB1, the adaptor of the CUL4 E3 ligase. This recruitment of the DDB1-CUL4 complex to CDK12 positions CCNK for polyubiquitylation and subsequent proteasomal degradation. Without the obligate cyclin partner, CDK12 can no longer form an active complex, preventing phosphorylation of RNA Pol II. This serendipitous discovery enabled medicinal chemistry to probe structural features required for complex formation, reduce inhibition of CDK family members, and optimize degradation potency.^12,14,19,24–26^

**Figure 1.**
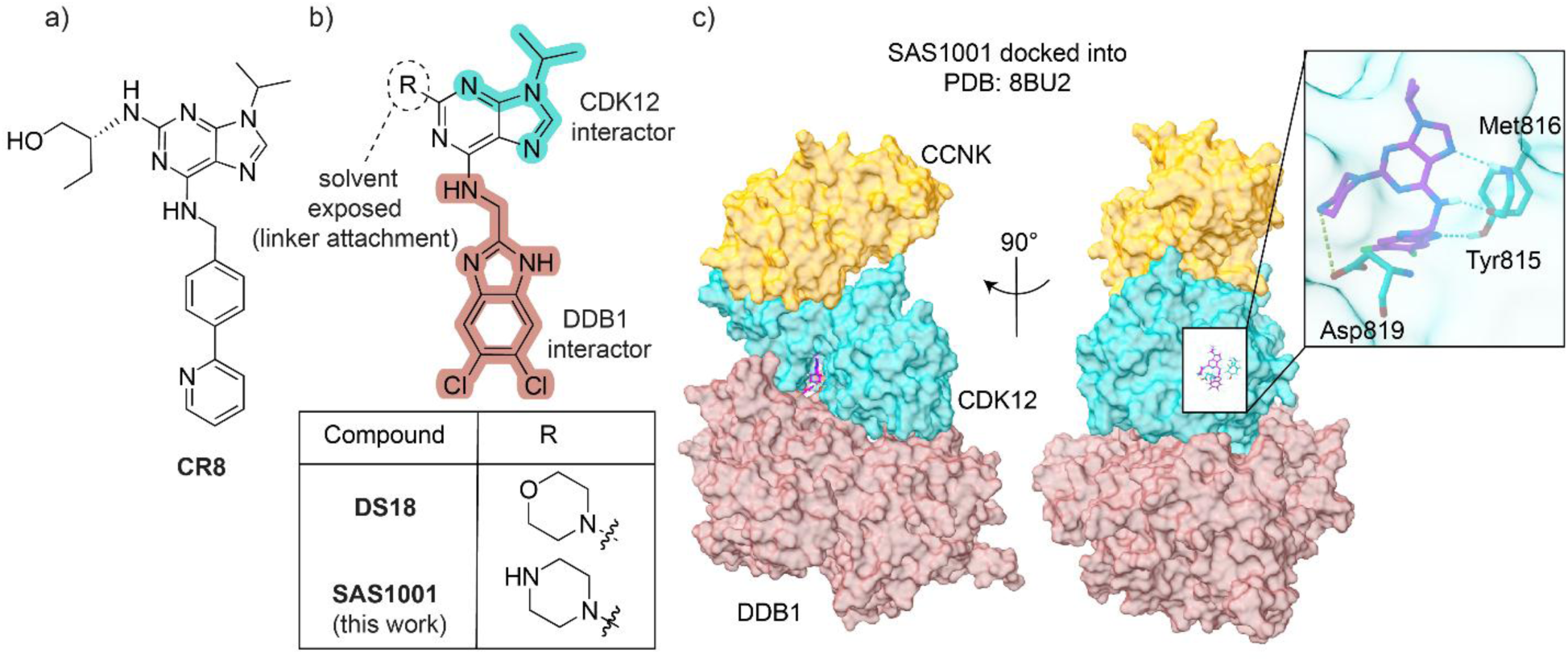
Design of the linker-compatible CCNK molecular glue degrader SAS1001 and proposed mechanism of antibody-directed CCNK degradation. **(a)** Structure of the pan-CDK inhibitor and selective CCNK degrader CR8.^16^ (b) DS18/SAS1001 common scaffold. Previous SAR studies identified the R substituent (circled) as largely solvent exposed,^12^ suggesting that replacement of the morpholine of DS18 with a piperazine would be tolerated, providing a linker attachment site while retaining the regions responsible for CDK12 (cyan) and DDB1 (salmon) engagement. Inset table: the R substituent of DS18 and SAS1001. (c) SAS1001 docked into the DDB1–CDK12/CCNK ternary complex (PDB 8BU2, 3.13 Å), shown in two orientations related by a 90° rotation, with CDK12 in cyan, DDB1 in salmon and CCNK in yellow. Inset: close-up of the docked pose of SAS1001 (purple) in the CDK12 ATP site, showing predicted hydrogen bonds (cyan dashed lines) to the hinge residues Tyr815 and Met816 and a predicted salt bridge (green dashed line) between the protonated piperazine and Asp819.

However, there are a number of potential challenges in the use of these highly potent and selective CCNK degraders as chemotherapeutic agents, since systemic degradation of this ubiquitously expressed protein can lead to on-target toxicity in normal tissues.^27–29^ Although CCNK is understood principally as the obligate cyclin of CDK12 and its paralogue CDK13, kinase-independent roles have been proposed, including direct interaction with β-catenin and regulation of Aurora B,^30,31^ which would broaden the consequences of systemic depletion. Thus, the ability to selectively direct the degrader to diseased tissue is desirable.

Increased expression of CDK12 in breast cancer is frequently accompanied by co-amplification of *ERBB2*, which encodes the human epidermal growth factor receptor 2 (HER2) transmembrane receptor tyrosine kinase.^8^ Approximately 15-20% of all breast cancers exhibit overexpression of HER2, and are termed HER2-positive.^32^ Historically, this was associated with a poor prognosis but with the advent of targeted therapeutics, such as trastuzumab (Tz) monoclonal antibodies and antibody-drug conjugates, survival has increased dramatically.^33^ The established clinical utility of HER2 as a tumor-associated antigen makes it an attractive target for selective delivery of a CCNK degrader, while the underlying biology of HER2-positive cancers provides a compelling rationale for this therapeutic strategy. In particular, the co-amplification of *ERBB2* and *CDK12* on the 17q12 amplicon has been associated with more aggressive tumor phenotypes and metastatic progression.^8,34,35^ Furthermore, HER2-driven tumors exhibit increased dependence on transcriptional programs downstream of oncogenic signaling pathways, including MYC, PI3K/AKT and MAPK,^36^ suggesting that disruption of transcriptional regulators such as the CDK12/CCNK complex may offer therapeutic benefit. Consistent with this, CDK12 has been implicated in Tz resistance, and CDK12 inhibition has been shown to restore Tz sensitivity in preclinical models.^8^ Together, these observations support the development of HER2-targeted CCNK degraders as a potential therapeutic approach for HER2-positive cancer.

Antibody-drug conjugates provide a clinically validated strategy for improving the selectivity of cytotoxic payloads by exploiting tumor-associated antigen-mediated internalization and lysosomal trafficking.^37–39^ Extending this concept to targeted protein degradation, degrader-antibody conjugates (DACs) can bias degrader delivery and activity to antigen-positive cells while retaining the catalytic pharmacology of the degrader.^40–43^ Indeed, we previously demonstrated HER2-dependent delivery of a BRD4 PROTAC using a Tz-based antibody-PROTAC conjugate.^40^ While antibody-PROTAC conjugates have demonstrated the feasibility of antibody-directed protein degradation, extension to molecular glue degraders presents distinct design challenges and remains comparatively less explored. Molecular glue degraders as payloads offer several potential advantages over bifunctional degraders, including lower molecular weight, simplified structures and highly efficient target degradation.^44,45^ In contrast to PROTACs, molecular glues do not possess two separable target-and E3-binding components, instead relying on induced or stabilized protein-protein interactions to achieve degradation.^46^ These induced protein-protein interactions can be highly sensitive to relatively minor structural modifications of the degrader scaffold.^47^ The development of linker-payload constructs typically relies upon molecular glue degraders that possess suitable functional handles, such as primary or secondary amines, to avoid compromising degradation activity. Consequently, molecular glue payloads explored for DAC generation have largely focused on immunomodulatory imide drug (IMiD)-based degraders.^48,49^ These have well-characterized SAR and established linker attachment sites that are extensively exploited in cereblon (CRBN)-recruiting PROTACs and readily adaptable for antibody conjugation.^50,51^ Antibody-delivered CRBN molecular glues have entered clinical evaluation, with two conjugates from Orum Therapeutics: the anti-CD33 GSPT1 conjugate ORM-6151 and the HER2-directed GSPT1 conjugate ORM-5029.^48,52–55^ There have also been further preclinical advances using the next-generation CRBN E3 ligase modulators (CELMoDs) in the context of degrader-antibody conjugates, with Guo *et al.* screening a modular CRBN-binder library to identify the conjugatable CELMoD payload I034, and demonstrating complete tumor eradication with the resulting CD38-targeted conjugate.^56^

Dependence on CRBN presents a potential route for resistance,^57^ as loss of CRBN is tolerated in cultured cells and in mice, and is even compatible with life in humans (albeit with neurodevelopmental effects).^58,59^ Molecular glue degraders acting through other E3 ligases may be less vulnerable to resistance, for example those involving DDB1, which is broadly essential.^60–63^ Delivery of a non-CRBN glue on an antibody-like scaffold has nonetheless been reported only once in the peer-reviewed literature, with the CCNK glue CR8 appended to a HER2-binding affibody that engages its antigen covalently and releases payload by disulfide exchange.^64,65^ Patent disclosures have additionally described trastuzumab conjugates of CCNK glues from a distinct chemotype.^66^ Whether a purine-series CCNK glue can be modified for conventional antibody conjugation while retaining degradation potency remains open.

We sought to extend antibody-directed protein degradation to encompass non-CRBN based molecular glue degraders. We report a Tz-based DAC for antibody-directed delivery of the CCNK molecular glue degrader SAS1001 (Figure 1b) to HER2-positive breast cancer cells (Figure S1). We demonstrate that conjugation to Tz confers preferential CCNK degradation and antiproliferative effects in HER2-positive cells, relative to a non-targeting control DAC. Whole-proteome analysis further supports engagement of the intended CCNK degradation pathway and reveals downstream molecular consequences of DAC treatment. Together, these findings establish antibody-directed delivery as a strategy for biasing the activity of CCNK molecular glue degraders towards antigen-positive cancer cells.

## Results

### Design and validation of a linker-compatible CCNK molecular glue degrader

A major obstacle to the development of molecular glue DACs is the sensitivity of glue-induced protein-protein interactions to structural modification, which can be disrupted by even minor chemical changes.^14^ Accordingly, we sought to determine whether a CCNK molecular glue degrader could be converted into a DAC-compatible payload while retaining degradation activity.

To develop a Tz-based DAC for delivery of a CCNK degrader to HER2-positive cells, we required a molecular glue scaffold that could accommodate installation of a self-immolative spacer. *Para*-aminobenzyl groups are often incorporated into cleavable linkers in order to release an unmodified payload^67^ and this requires a suitable functional group on the payload; typically, a hydroxyl (*p*-aminobenzylcarbonate) or amine (*p*-aminobenzylcarbamate, PABC).^68^ We opted for a carbamate approach given the precedent for increased stability versus carbonates.^69,70^

Kozicka *et al.* performed extensive SAR studies on the CR8 scaffold (Figure 1a), which indicated that the 2-position (R, Figure 1b) was primarily solvent exposed, with limited interactions with CDK12.^12^ Thus, by taking one of the most promising hits, DS18 (Figure 1b), and substituting the morpholine in the R position to a piperazine, we developed SAS1001 bearing a linker-compatible amine at R (Figure 1b). Molecular docking into the DDB1-CDK12-CCNK complex suggested that SAS1001 retains the interaction mode of DS18, occupying the interface between DDB1 and CDK12 (Figure 1c). An additional salt bridge was predicted to form between the charged piperazine and Asp819 on CDK12 (Figure 1c), providing a potentially stabilizing interaction (Figure S2a, S2b), an interaction previously postulated by Kuchukulla *et al.* in their SAR study into the CR8 scaffold.^24^ The binding poses of DS18 and SAS1001 were closely aligned (Figure S2), suggesting that modification to a piperazine may be tolerated. At the time this work was initiated, only pan-CDK inhibitors possessed amines suitable for direct DAC development.^24^ While this work was underway, Zhang, Ghosh and Pandurangan *et al.* reported piperazine-containing derivatives of the scaffold.^19,25,26^ However, these compounds were not developed as linker attachment handles for DAC generation, with traceless release in mind. Synthesis of SAS1001 proceeded through a 5-step synthetic route (Scheme S1).^12^

To assess the ability of SAS1001 to induce ternary complex formation, we employed a time-resolved fluorescence resonance energy transfer (TR-FRET) assay previously developed and optimized by Kozicka *et al*.^12^ SAS1001 promoted ternary complex formation between CDK12/CCNK and DDB1 with an EC₅₀ of 3.3 nM, compared with 8.4 nM for CR8 assayed in parallel (Figure 2a). These values were broadly consistent with those previously reported for degraders in this series, including DS18, supporting retention of potent ternary complex formation following the morpholine-to-piperazine substitution.^12^

**Figure 2.**
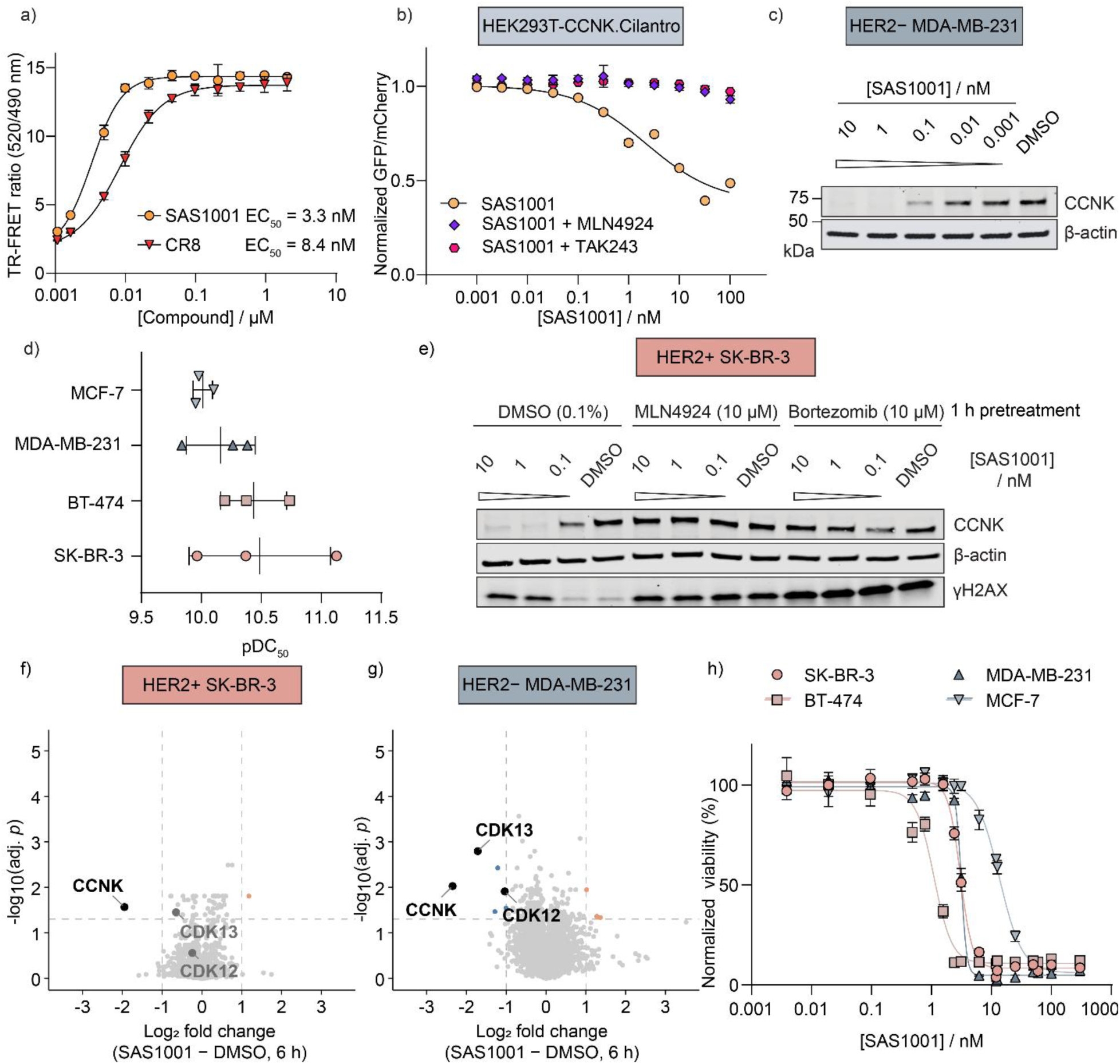
**SAS1001 induces potent and selective CCNK degradation through a Cullin-and proteasome-dependent mechanism. (a)** TR-FRET assay measuring glue-induced ternary complex formation between CDK12–cyclin K and DDB1. Biotinylated CDK12–cyclin K (50 nM), Alexa Fluor 488-labelled DDB1 (100 nM) and terbium-labelled streptavidin (4 nM) were incubated SAS1001 or CR8 and the TR-FRET ratio (520/490 nm) was recorded. Curves are four-parameter nonlinear regression fits with the bottom constrained to the no-compound control. Mean ± SD of three technical replicates from a single experiment. **(b**) CCNK degradation in HEK293T-CCNK.Cilantro reporter cells, assessed as the decrease in GFP fluorescence normalized to mCherry, following 16 h treatment with SAS1001 alone or after 1 h pretreatment with MLN4924 (1 µM) or TAK243 (1 µM). Mean ± SD of technical triplicates from a single biological replicate. The SAS1001 dose–response was performed in three independent biological replicates (DC₅₀ = 0.92 nM, pDC₅₀ = 9.0 ± 0.4); pretreatment with MLN4924 and TAK243 was performed in biological duplicate. **(c)** Representative western blot showing concentration-dependent CCNK degradation following 24 h treatment with SAS1001 in HER2-negative MDA-MB-231 cells. Western blots are representative of three independent biological replicates, performed in each line. **(d)** Apparent pDC₅₀ values for SAS1001-induced CCNK degradation in HER2-positive (SK-BR-3, BT-474) and HER2-negative (MDA-MB-231, MCF-7) cells, derived from four-parameter fits of three independent biological replicates. Points are individual replicates, line and whiskers show mean ± SD **(e)** Pretreatment of HER2-positive SK-BR-3 cells with MLN4924 or bortezomib (10 µM) for 1 h prior to SAS1001 treatment rescues CCNK levels, consistent with a Cullin-and proteasome-dependent mechanism. Accumulation of γH2AX was seen at the top two concentrations of SAS1001, but also upon pretreatment by the UPS inhibitors. Representative blot, performed in biological triplicate. DIA-based whole-proteome analysis in SK-BR-3 **(f)** and MDA-MB-231 **(g)** following 6 h treatment with SAS1001. Differential protein abundance was assessed by two-sided Student’s t-test; p-values were corrected by the Benjamini–Hochberg procedure. Proteins passing adj. p < 0.05 and |log₂FC| > 1 are colored by direction (orange, increased; blue, decreased), with dashed lines marking the thresholds. CCNK, CDK12 and CDK13 are labelled throughout, in black where they pass both thresholds and grey where they do not. Four replicates per condition (three for SAS1001 in **(f)**, following exclusion of one outlier) **(h)** Dose-response proliferation assays following 5 days treatment with SAS1001 in HER2-positive (SK-BR-3, BT-474) and HER2-negative (MDA-MB-231, MCF-7) breast cancer cell lines, demonstrating potent antiproliferative activity across multiple models. Curves are from one representative biological replicate, with mean ± SD of technical triplicates; summary IC₅₀ values from three independent biological replicates are given in Figure S8.

We next assessed SAS1001 in the HEK293T-CCNK.Cilantro reporter line.^12^ SAS1001 induced CCNK_eGFP_ degradation with a sub-nanomolar DC_50_ of 0.92 nM (Figure 2b; Figure S3). CR8 and SR-4835, assayed in parallel, gave DC₅₀ values of 13 nM and 6.0 nM respectively (pDC_50_ = 7.9 ± 0.3 and pDC_50_ = 8.22 ± 0.19, Figure S3b), confirming the high cellular potency of SAS1001 relative to these reference compounds (previously reported as 24 and 14 nM, respectively)^12^. Both findings suggest that the morpholine-to-piperazine substitution is well tolerated and preserves the potent CCNK degradation activity characteristic of the DS18 scaffold. We explored the ubiquitin-proteasome system (UPS) dependence in the reporter assay, with 1 h pretreatment with either MLN4924 or the ubiquitin-activating enzyme (E1) inhibitor TAK243. Both abolished SAS1001-induced CCNK_eGFP_ loss (Figure 2b), consistent with degradation requiring Cullin neddylation and ubiquitin conjugation.

Based on DepMap RNA expression data (Figure S4a) and their established use as models for HER2-targeting ADCs, BT-474 and SK-BR-3 were selected as representative HER2-positive cell lines, and MDA-MB-231 and MCF-7 as HER2-negative lines.^62,63^ Western blot analyses confirmed protein expression of cathepsin B (CatB) in all selected cell lines, and appropriate selective expression of HER2 (Figure S4b). CatB is a lysosomal cysteine protease frequently upregulated in invasive and metastatic cancers,^71^ and commonly exploited to trigger intracellular payload release in ADCs. CDK12 and CCNK expression across the panel are shown in Figure S5.

Western blot analyses demonstrated that SAS1001 induced potent, dose-dependent CCNK degradation across all four cell lines (Figure 2c, 2d; Figure S5). The maximal level of degradation (D_max_) exceeded 99% in SK-BR-3, BT-474 and MDA-MB-231 cells, consistent with complete degradation, whereas residual CCNK remained quantifiable in MCF-7 cells at 83% degradation (Figure S5n). One possibility is that this reflects cell line dependent differences in synthesis rate of CCNK.^72^ Apparent DC₅₀ values, estimated by densitometry, spanned a 3-fold range across the panel, from 32 pM in SK-BR-3 to 97 pM in MCF-7 cells (pDC₅₀ 10.5 ± 0.6 and 10.01 ± 0.08 respectively; Figure 2d, Figure S5m, S5n).

At the 24 h timepoint, CDK12 levels were variable, while CCNK was consistently depleted across the cell lines examined (Figure S5a-l). This is consistent with previous structural and mechanistic studies in which CDK12 functions as a “bridge” protein within the glue-induced complex with DDB1, positioning CCNK for ubiquitylation and preferential degradation.^73,74^ We therefore focused subsequent western blot studies solely on CCNK degradation. The much lower apparent DC₅₀ obtained by western blot in the breast cancer lines (32–97 pM) likely reflects differences in cell background and in the turnover of overexpressed CCNK_eGFP_ relative to endogenous CCNK, and the two assay formats are therefore not directly comparable. Taken with the TR-FRET data, we interpret SAS1001 as comparable to or modestly more potent than DS18.

The UPS dependence was further investigated in the endogenous line SK-BR-3, where pretreatment for 1 h with either bortezomib (proteasome inhibitor) or MLN4924 followed by 24 h treatment by SAS1001 led to an almost complete rescue of degradation of CCNK (Figure 2e; Figure S6), consistent with retention of a proteasome and Cullin-dependent degradation mechanism via the UPS. The presence of DNA damage breaks as indicated by the accumulation of γH2AX at the two highest concentrations suggested that treatment with SAS1001 induced a DNA damage response (Figure 2e; Figure S6),^75^ in line with the known downstream effects of CCNK depletion.^5^ γH2AX was also induced by bortezomib and MLN4924 alone.

To define the timescale of degradation and the onset of downstream effects, whole proteome analysis by data-independent acquisition (DIA) was performed in SK-BR-3 and MDA-MB-231 as separate experiments after 2, 6 and 24 h of SAS1001 treatment, each against their corresponding DMSO controls (Figure 2f, 2g; Figure S7). CCNK depletion was progressive, reaching log₂ fold changes of −1.95 (SK-BR-3) and −2.35 (MDA-MB-231) at 6 h, and −2.76 in SK-BR-3 at 24 h. Over the same period the number of other proteins passing both thresholds rose from none at 2 h to one other than CCNK at 6 h and 150 at 24 h in SK-BR-3. We were unable to quantify the proteomic changes for MDA-MB-231 after 24 h due to low protein quantity, as a result of cell death. We therefore selected 6 h to assess degradation selectivity, capturing primary degradation before the accumulation of secondary transcriptional and proteomic consequences.

At 6 h, CCNK was among the most significantly depleted proteins in both lines, where it was one of two in 7,270 quantified proteins altered in SK-BR-3 and 11 of 7,599 in MDA-MB-231 (Figure 2f, 2g), supporting the conclusion that SAS1001 is a selective CCNK degrader. In MDA-MB-231 cells the partner kinases CDK12 and CDK13 were also significantly depleted (Figure 2g); in SK-BR-3 both were reduced at 6 h but did not meet the significance thresholds. Degradation proceeded at least as rapidly in HER2-negative MDA-MB-231 as in HER2-positive SK-BR-3 at every timepoint measured, indicating that the unconjugated degrader has no intrinsic kinetic preference for HER2-positive cells.

SAS1001 also showed strong antiproliferative effects across a range of breast cancer cell lines (Figure 2h, Figure S8), with IC_50_ values in the low nanomolar spanning a 14-fold range from 1.0 nM (pIC_5_= 8.99 ± 0.10) for the most sensitive line, BT-474, to 14 nM (pIC_50_ = 7.86 ± 0.03) for MCF-7. In comparison, degradation potency varied only 3-fold (DC_50_ 32–97 pM; Figure S5n), indicating that the differences in proliferation reflect cell-line-intrinsic sensitivity to CCNK loss rather than differential degradation. Against MDA-MB-231, the HER2-negative line used as the comparator throughout this study and the more sensitive of the two HER2-negative models, SAS1001 was only 1.7-fold (SK-BR-3) and 3.8-fold (BT-474) more potent, establishing that the degrader itself has little intrinsic preference for the HER2-positive lines.

Thus, we demonstrated that the morpholine to piperazine switch is well-tolerated, and preserves degradation potency and mechanism, with no evidence of altered degradation by whole-proteome analysis. Importantly, the phenotypic assays showed that the degrader alone did not display any significant inherent preference for HER2-positive cells. With SAS1001 biologically characterized, we next sought to prepare the linker-payload construct.

### Design and synthesis of linker-payload constructs

To enable intracellular release of the unmodified degrader, we selected a valine-alanine (VA) dipeptide linker in conjunction with a PABC group and a polyethylene glycol (PEG) spaced terminal maleimide (mal) (Figure 3a). This clinically validated ADC linker has been shown to have favorable stability *in vivo* and compatibility with hydrophobic payloads.^76,77^ The VA dipeptide provides a CatB cleavable trigger,^78–80^ while the PABC moiety undergoes spontaneous 1,6-elimination following proteolysis to release the unmodified payload (Figure S9a).^79,81^

**Figure 3.**
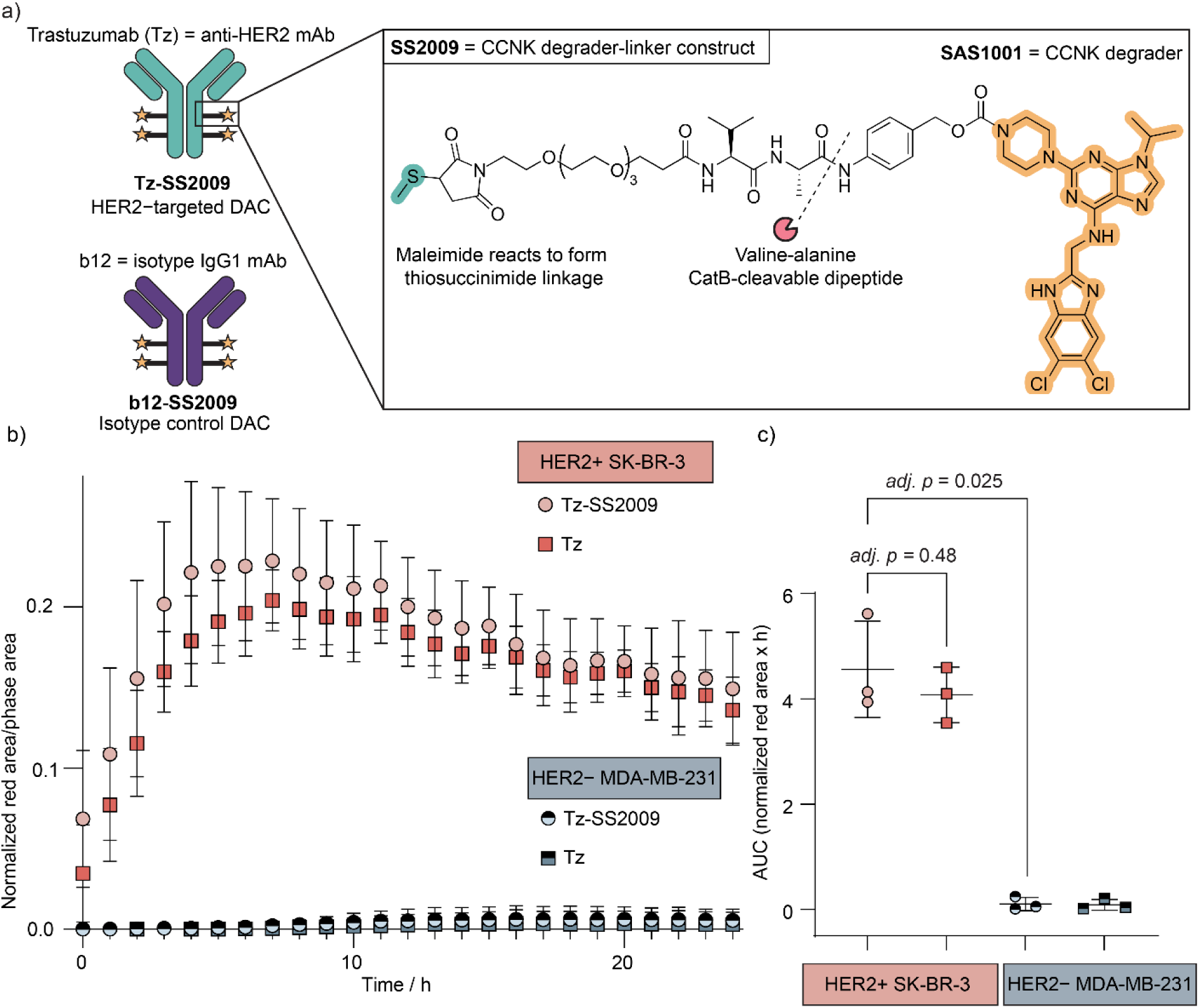
Tz-SS2009 internalizes preferentially into HER2-positive cells. **(a)** Structure of linker-payload SS2009, incorporating the CCNK degrader SAS1001 and schematic representation of the HER2-targeted degrader-antibody conjugate Tz-SS2009 and isotype control DAC b12-SS2009. **(b)** Quantification of internalization using the FabFluor system demonstrates enhanced uptake of Tz-SS2009 and Tz in SK-BR-3 cells, while minimal uptake was observed in HER2-negative MDA-MB-231 cells. Internalization was defined as red area per image with background (buffer only) subtracted and normalized to the phase (confluence) area per image. Mean ± SD for a representative assay performed in biological triplicate. **(c)** Area under the curve for the time courses in **(b)**, calculated per biological replicate after subtraction of the buffer-only control. Two pre-specified comparisons were analyzed by Welch’s unequal-variance t-test with Holm correction: Tz-SS2009 in SK-BR-3 versus MDA-MB-231 (adj. p = 0.025), and Tz-SS2009 versus Tz within SK-BR-3 (adj. p = 0.48). Points are individual biological replicates; bars show mean ± SD (n = 3).

The synthetic scheme to generate linker-degrader SS2009 (Scheme S2) was initiated by transformation of the terminal hydroxyl of alloc protected VA-PAB-OH into an electrophilic *p*-nitrophenyl carbonate (6) using bis(4-nitrophenyl)carbonate prior to carbamate formation with SAS1001. Alloc deprotection under typical conditions yielded the VA-PABC-degrader construct (8). Finally, the maleimide (mal) and PEG linker (PEG4) were installed by amide coupling under mild conditions using an activated *N*-hydroxysuccinimide ester to give the linker-payload mal-PEG4-VA-PABC-SAS1001 (SS2009).

Incubation of SS2009 with CatB (from human liver) resulted in progressive consumption of SS2009 and the generation of free SAS1001, as monitored over the course of 16 h by reversed-phase high-performance chromatography (RP-HPLC), confirming both enzymatic cleavage and traceless payload release (Figure S9b, S9c).

### Antibody conjugation and characterization

Targeted DAC Tz-SS2009 (anti-HER2) and isotype control b12-SS2009 (anti-HIV gp120), Figure 3a, were synthesized by tris(2-carboxyethyl)phosphine (TCEP)-mediated partial reduction of the mAb disulfide bonds, followed by Michael addition to the maleimide-containing SS2009. b12 does not recognize any known human cell-surface antigens and it is widely used as a negative control for evaluating targeting-independent antibody-associated effects in preclinical studies.^82,83^ Likewise, Tz was chosen due to its extensive characterization and use as an antibody for HER2-targeting, including in the clinic.^84^ Both conjugates were assessed to have an average drug-antibody ratio (DAR) of approximately 4 by polymeric reversed-phase chromatography (PLRP) and hydrophobic interaction chromatography (HIC) (Figure S10, S11). DAR ∼4 is comparable to many clinically approved ADCs and represents a balance between payload delivery, tolerable immunogenicity and conjugate aggregation.^85^ Importantly, the conjugation strategy yielded DACs with similar DARs, enabling subsequent comparison of the effects of HER2-directed versus non-targeted delivery.

### HER2-targeted conjugates internalize preferentially in HER2-positive cells

In order to assess whether the addition of a hydrophobic payload affected the otherwise rapid internalization by Tz,^86^ internalization was monitored in real time using the pH-responsive Sartorius FabFluor labelling system (Figure 3b, 3c). DACs or monoclonal antibodies were incubated with FabFluor reactive fragments, which fluoresce red upon exposure to the acidic environment of endosomal and lysosomal compartments (Figure S12). Tz-SS2009 and unconjugated Tz produced red fluorescence signals in HER2-positive SK-BR-3 cells, whereas signal in HER2-negative MDA-MB-231 cells remained at or near baseline throughout the experiment. Quantification of FabFluor signal by area-under-the-curve (AUC) analysis confirmed significantly greater internalization of Tz-SS2009 in HER2-positive SK-BR-3 than HER2-negative MDA-MB-231 cells (Welch’s t-test with Holm correction, adj. *p* = 0.025; Figure 3c), consistent with HER2-mediated internalization and trafficking of the DAC to acidic compartments. AUC did not differ significantly between Tz-SS2009 and unconjugated Tz in SK-BR-3 cells (adj*. p* = 0.48) indicating that payload conjugation does not impair receptor-mediated internalization.

### Tz conjugation imparts preferential HER2-directed activity to the CCNK molecular glue degrader

Next, we sought to investigate both whether DACs retained the CCNK-degrading capabilities of the free degrader, and whether conjugation to Tz leads to measurable HER2-targeted selectivity. Given the effects on cell proliferation that we had seen with SAS1001 and to avoid selection for cell populations capable of tolerating sustained CCNK loss, a 72 h timepoint was chosen for DAC degradation assays. We hypothesized that this would allow for DAC internalization, payload release and CCNK degradation prior to extensive secondary proteomic and transcriptional changes.

Tz-SS2009 induced potent CCNK degradation in HER2-positive cells, with substantial CCNK depletion observed at low nanomolar concentrations (Figure 4a; Figure S13). Densitometry indicated a DC_50_ of 270 pM (pDC_50_ = 9.57 ± 0.10) in BT-474 and 210 pM (pDC_50_ = 9.7 ± 0.6) antibody equivalents in SK-BR-3 (Figure S13n). Accounting for DAR, this is approximately 1 nM in payload equivalents, making the DAC around 25-to 30-fold less potent than the free degrader. Although some degradation was observed following treatment with the isotype control DAC b12-SS2009 at the top concentration of 10 nM, significantly more degradation was seen following treatment with Tz-SS2009 (Figure 4b). Densitometric analysis of the western blots revealed a 45-fold reduction in CCNK abundance in Tz-SS2009-treated BT-474 cells relative to b12-SS2009-treated cells at 10 nM DAC (Figure 4b, adj. *p* < 0.0001).

**Figure 4.**
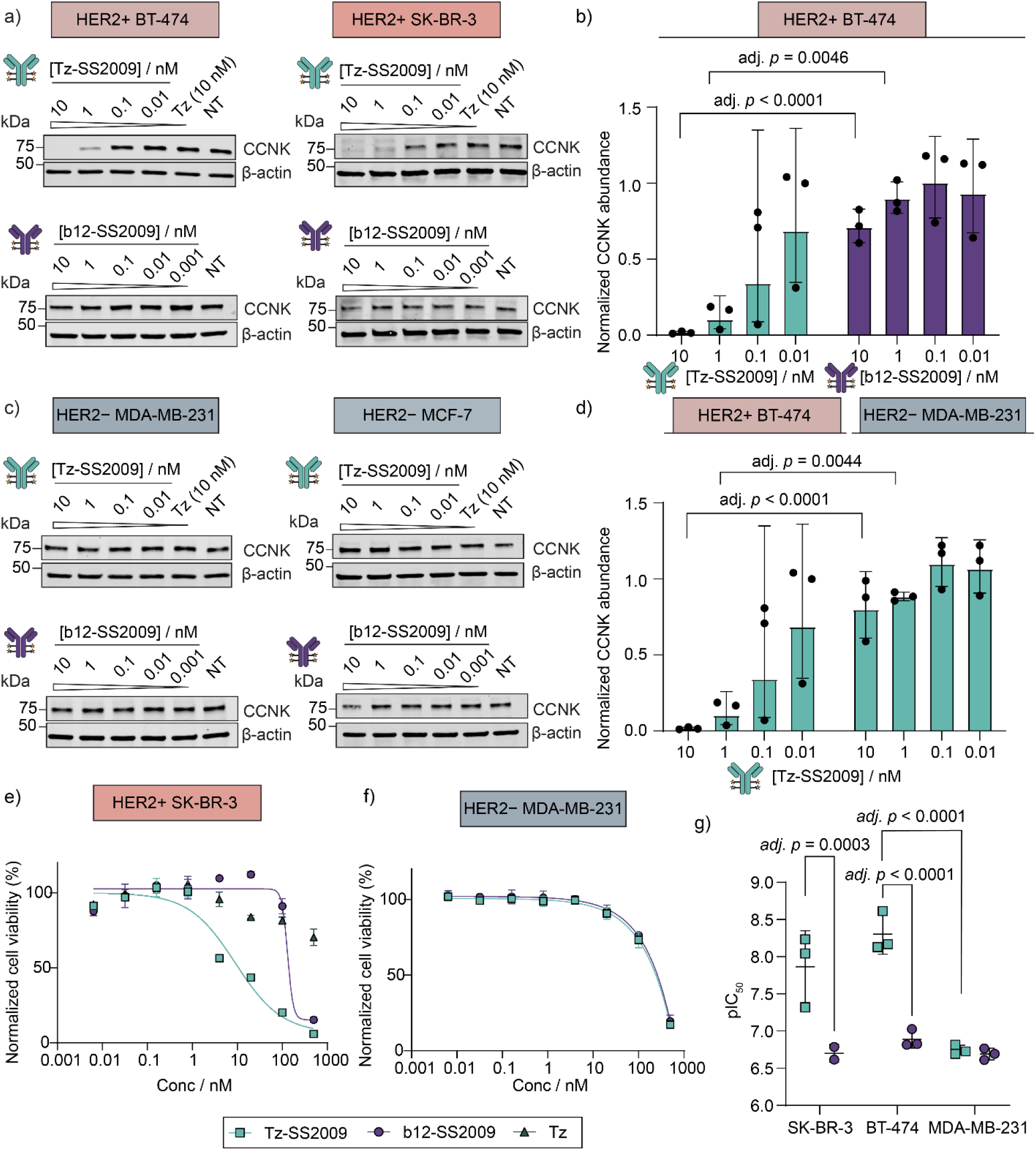
Tz-mediated delivery shifts CCNK degradation and antiproliferative activity towards HER2-positive cells. **(a)** Representative western blots showing concentration-dependent CCNK degradation after 72 h treatment with Tz-SS2009 or the non-targeting isotype control DAC b12-SS2009 in HER2-positive BT-474 and SK-BR-3 cells. Unconjugated Tz (10 nM) and untreated (NT) controls are included on the Tz-SS2009 blots; NT alone on the b12-SS2009 blots. Western blots are representative of three independent biological replicates. **(b)** Quantification of CCNK abundance in BT-474 cells following treatment with Tz-SS2009 or b12-SS2009. At 10 nM (45-fold lower CCNK abundance, adj. p < 0.0001) and 1 nM (8.6-fold lower, adj. p = 0.0046), Tz-SS2009 induced significantly greater degradation than b12-SS2009 as assessed by ordinary two-way ANOVA with Šídák’s multiple-comparisons. All comparisons, effect sizes and confidence intervals are given in Figure S15. Geometric mean ± geometric SD factor of three independent biological replicates (shown as points). **(c)** Representative western blots showing limited CCNK degradation after 72 h treatment in HER2-negative MDA-MB-231 and MCF-7 cells, with treatments as in **(a). (d)** Quantification of CCNK abundance following Tz-SS2009 treatment in HER2-positive BT-474 and HER2-negative MDA-MB-231 cells. Geometric mean ± geometric SD factor of three independent biological replicates (shown as points). **(e)** Dose–response proliferation assay in HER2-positive SK-BR-3 cells treated with Tz-SS2009, b12-SS2009 or unconjugated Tz for 9 days. Curves are fitted for Tz-SS2009 and b12-SS2009; Tz did not reach 50% inhibition at the highest concentration tested (500 nM) and was not fitted. **(f)** Dose–response proliferation assay in HER2-negative MDA-MB-231 cells treated with Tz-SS2009 or b12-SS2009 for 9 days. Unconjugated Tz was not tested in this cell line. Data in **(e)** and **(f)** are from one representative biological replicate (mean ± SD of technical triplicates) with four-parameter nonlinear regression fits; summary IC₅₀ values from three biological replicates are given in Figure S16e. **(g)** pIC₅₀ values for Tz-SS2009 and b12-SS2009, derived from four-parameter fit and analyzed by ordinary two-way ANOVA, where the interaction was significant (F(2, 11) = 12.58, p = 0.0014). Displayed p values are from Šídák’s multiple-comparisons test; all comparisons, effect sizes and confidence intervals are given in Figure S17. Points are individual biological replicates; bars show mean ± SD. All concentrations refer to antibody (conjugate) equivalents.

Little to no CCNK degradation was observed in HER2-negative cells following treatment with either DAC (Figure 4c; Figure S14). Comparison of Tz-SS2009 treatment in HER2-positive BT-474 and HER2-negative MDA-MB-231 cells revealed a 51-fold reduction in CCNK abundance in BT-474 cells at 10 nM (Figure 4d, adj. *p* < 0.0001). These comparisons between both delivery of Tz-SS2009 into HER2-positive versus HER2-negative cells, as well as the decreased ability of the isotype control to enter either cell at the matched antibody concentration allowed us to identify a widening cellular selectivity window for HER2-overexpressing cells (Figure S15).

### Redistribution of CCNK degradation activity results in selective growth inhibition

Having established that our DACs could induce CCNK degradation in HER2-positive cells, we next investigated their antiproliferative activity. HER2-positive SK-BR-3 cells displayed an approximately 14-fold greater sensitivity to Tz-SS2009 after 9 days of treatment than to the isotype control b12-SS2009 (Figure 4e; Figure S16a, S16e), with an IC_50_ of 14 nM (pIC_50_ = 7.9 ± 0.5) compared with 200 nM (pIC_50_ = 6.70 ± 0.12). Unconjugated Tz did not reach 50% inhibition of cellular proliferation at concentrations up to 500 nM (61% viability at 500 nM; Figure S16e) in SK-BR-3 cells, indicating that the enhanced potency of Tz-SS2009 arises from antibody-directed delivery of the CCNK degrader rather than from HER2 pathway blockade by Tz itself.

Consistent with the degradation data, in HER2-negative MDA-MB-231 cells Tz-SS2009 and b12-SS2009 were indistinguishable (IC_50_ 180 nM, pIC_50_ = 6.75 ± 0.06 and 200 nM, pIC_50_ = 6.69 ± 0.08 respectively; Figure 4f; Figure S16b, S16e). At higher DAC concentrations, antiproliferative activity was seen in MDA-MB-231 cells with a viability at the top concentration of Tz-SS2009 and b12-SS2009 dropping to 17% and 23%, respectively. This may be indicative of low-level antigen-independent uptake and release of the degrader, or premature cleavage of the linker. MCF-7, the less sensitive of the two HER2-negative lines (SAS1001 IC₅₀ 14 nM versus 3.8 nM) and treated for 7 rather than 9 days, showed no corresponding loss of viability (Figure S16d). This reduced treatment length was necessary due to generalized toxicity seen after 9 days, attributed to nutrient depletion. We did not observe any toxicity at the concentrations assayed (IC_50_ > 500 nM, Figure S16d, S16e), although due to the difference in assay design, MCF-7 was excluded from subsequent matched analysis.

A shift in IC_50_ between Tz-SS2009 and b12-SS2009 was also observed in BT-474 cells (5.0 nM (pIC_50_ = 8.3 ± 0.3) versus 130 nM (pIC_50_ = 6.89 ± 0.12), Figure S16e). However, as expected, BT-474 cells were highly sensitive to unconjugated Tz,^87,88^ with Tz-SS2009 and Tz having indistinguishable IC_50_ values (5.0 vs 5.9 nM (pIC_50_ = 8.2 ± 0.2)). This was not seen for SK-BR-3 and confounded direct attribution of antiproliferative effects to CCNK degradation. Both DACs produced a lower plateau than Tz (Tz-SS2009 6%, b12-SS2009 2.9%, versus 19% for Tz). That both conjugates exceed Tz at this concentration, irrespective of targeting, is consistent with a payload-driven mechanism distinct from HER2 blockade but does not itself demonstrate antigen-dependent delivery at 500 nM.

Together, these findings demonstrate antibody-directed enhancement of antiproliferative activity in SK-BR-3 cells. Within each HER2-positive line, Tz-SS2009 was significantly more potent than the non-targeting b12-SS2009 (SK-BR-3, adj. *p* = 0.0003, BT-474, adj. *p* < 0.0001; Figure 4g). Across cell lines, conjugation to trastuzumab afforded a cellular selectivity of 13-fold (SK-BR-3) and 36-fold (BT-474) relative to MDA-MB-231 (adj. *p* = 0.0005 and adj. *p* < 0.0001 respectively; Figure S17), an approximately 8-to 10-fold improvement over the unconjugated degrader SAS1001 (Figure 2h; Figure S8). However, the inherent sensitivity of BT-474 to Tz provided challenges in assessing the extent to which antiproliferative effects could be attributed to the DAC rather than Tz monotherapy.^87,88^

### HER2-directed CCNK degradation induces downstream proteome changes

To investigate the early downstream effects of DAC treatment and deconvolute any effect from Tz itself, we performed DIA-based whole proteome analysis after 24 h of treatment in HER2-positive SK-BR-3 and BT-474 and in HER2-negative MDA-MB-231 cells. Tz-SS2009, Tz and b12-SS2009 (10 nM) were each compared to the formulation buffer in all three lines (Figure 5a; Figure S18). Each of these paired comparisons addresses a different question (Figure 5a; Figure S19). Tz-SS2009 versus the isotype control b12-SS2009 (non-binding IgG1 against an unrelated antigen) within a HER2-positive line matches payload and linker, isolating the contribution of antigen recognition. Because b12-SS2009 is not expected to undergo antigen-mediated internalization in either line, any effect of this conjugate indicates non-specific uptake or release. Tz-SS2009 versus unconjugated Tz in the same line isolates the consequences of the delivered payload, although conjugation may itself alter antibody behavior. Tz-SS2009 in HER2-positive versus HER2-negative lines reveals preferential activity associated with HER2 expression, although this comparison is additionally confounded by intrinsic differences between the lines.

**Figure 5.**
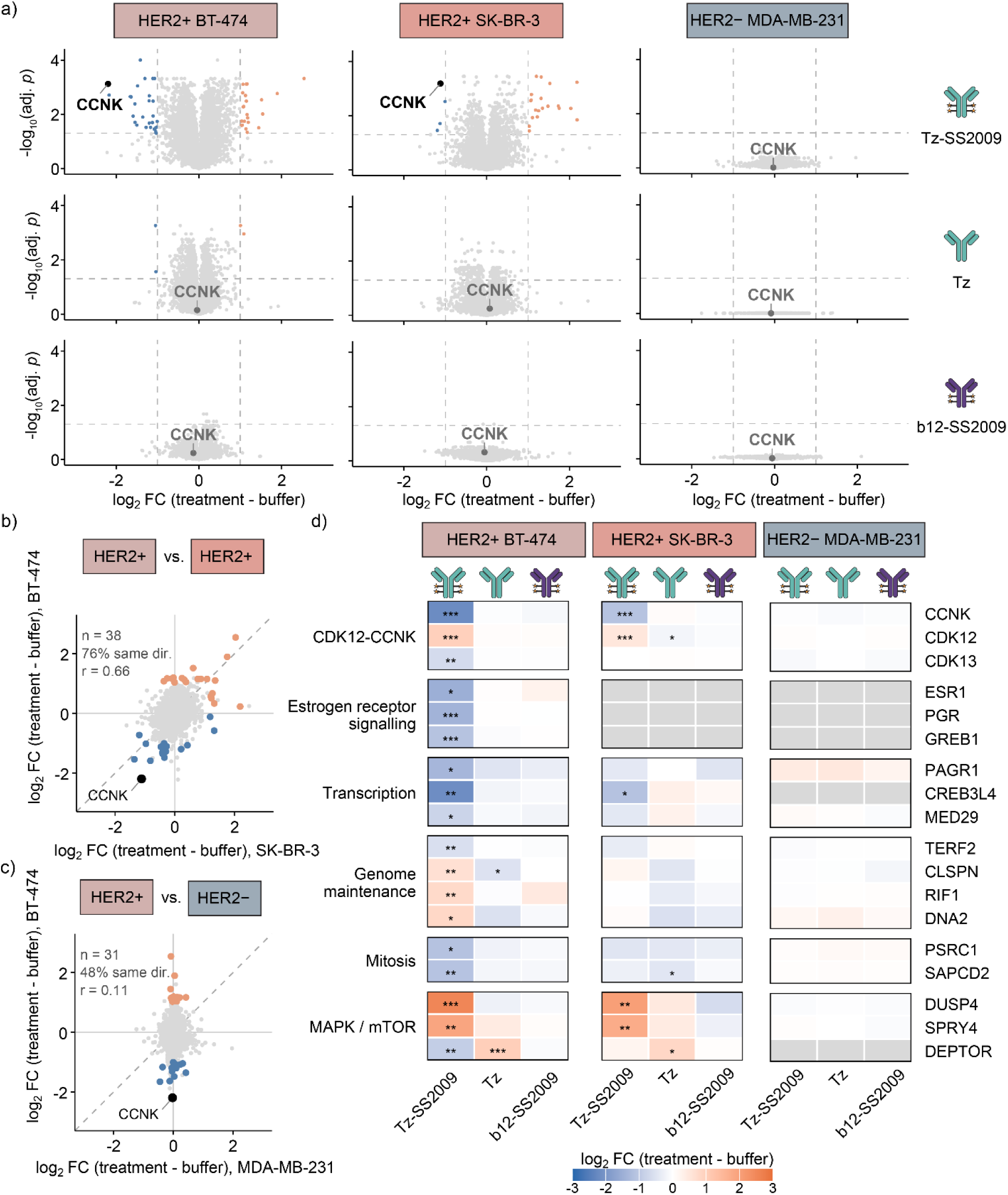
HER2-targeted delivery of Tz-SS2009 induces selective CCNK depletion and conserved downstream proteomic remodeling in HER2-positive breast cancer cells. BT-474, SK-BR-3 and MDA-MB-231 were treated for 24 h with 10 nM Tz-SS2009, Tz or b12-SS2009, or with formulation buffer, and analyzed by label-free DIA proteomics (n = 4, except Tz-SS2009 in BT-474, n = 3). All comparisons are against the buffer control of the same cell line by two-sided Student’s t-test; p-values were corrected by the Benjamini–Hochberg procedure within each cell line × treatment comparison, and significance is adj. p < 0.05 throughout. Immunoglobulin proteins were excluded before correction (Figure S18). (a) Volcano plots for each treatment (rows) and cell line (columns), on identical axes. Proteins passing adj. p < 0.05 and |log₂FC| > 1 are colored by direction (orange, increased; blue, decreased); dashed lines mark the thresholds. (b) and (c) Concordance of the Tz-SS2009 response between cell lines. All proteins quantified in both lines are plotted in grey; those passing threshold in either line are colored by their direction in BT-474. The dashed diagonal is y = x. Directional agreement is the proportion of colored proteins whose fold change has the same sign in both lines; where the second line carries no related response its values scatter about zero and the expected proportion is 50%. The two HER2-positive lines agree for 29 of 38 proteins (76%, binomial p = 0.0017, Pearson r = 0.66). (b) Against MDA-MB-231, agreement is 15 of 31 (48%, p = 1.0, r = 0.11), indistinguishable from chance, with the colored proteins collapsing onto x ≈ 0 (c). The statistical test performed in (b) and (c) was an exact two-sided binomial test against 50%. (d) Proteins changed by Tz-SS2009 in BT-474, the most responsive line, shown across all three cell lines and all three treatments. Rows were selected in BT-474 alone; SK-BR-3 and MDA-MB-231 are displayed without further filtering. Other than CDK12/CDK13/CCNK, rows were retained only where the protein changed by more than 1.4-fold against buffer (adj. p < 0.05) and in the same direction against trastuzumab at matched antibody exposure (change attributable to the delivered payload rather than to HER2 blockade). Tile color denotes magnitude and asterisks significance (* adj. p < 0.05, ** adj. p < 0.01, *** adj. p < 0.001). Grey tiles are proteins not quantified in that cell line. The complete set of proteins changed across the three lines is given in Figure S20.

Consistent with degradation of the intended target, whole proteome analysis showed significant depletion of CCNK in both HER2-positive lines treated with the HER2-targeting DAC Tz-SS2009 (log₂ fold change −2.20 in BT-474 and −1.12 in SK-BR-3, Figure 5a; Figure S18). BT-474 cells appeared more sensitive to Tz-SS2009, with 51 proteins changed compared to 24 in SK-BR-3, consistent with our earlier characterization of SAS1001 (Figure 2; Figure S7). In contrast, no proteomic perturbation was seen in HER2-negative MDA-MB-231 cells with the same treatment, supporting preferential delivery of the CCNK degrader to HER2-positive cells. In addition, the IgG1 proteins IGHG1 (BT-474 and SK-BR-3) and IGKC (SK-BR-3 only) were upregulated only in HER2-positive cells treated with either Tz-SS2009 or Tz, and neither was quantified in MDA-MB-231 under any condition (Figure S18). Neither of these proteins is expected in these cell lines, and their presence is known to be suggestive of cell-associated Tz.^89^ The isotype control b12-SS2009 is also derived from an IgG1 protein but shows no enrichment in any cell line, indicating absence of binding and internalization, giving further orthogonal support to selective HER2-targeted DAC uptake.

The response to Tz-SS2009 was shared between the two HER2-positive lines, where of the proteins changing in SK-BR-3 or BT-474, 76% moved in the same direction in both (29 of 38 proteins, binomial p = 0.0017, Pearson r = 0.66; Figure 5b). In contrast, against MDA-MB-231 agreement fell to chance (15 of 31 proteins, 48%, p = 1.0, r = 0.11), with the BT-474-responsive proteins clustering at zero (Figure 5c). The response in the HER2-negative line was therefore absent rather than merely weaker. The isotype control b12-SS2009 produced no significant proteomic changes in any cell line (Figure 5a), indicating that antigen-independent uptake is negligible at 10 nM over 24 h.

Within the target complex itself, CDK13 was also depleted by Tz-SS2009 in BT-474 (Figure 5d). Interestingly, CDK12 abundance increased rather than decreased in both HER2-positive lines (Figure 5d), in contrast to its depletion by free SAS1001 (Figure 2g). In our characterization of SAS1001 by Western blot, we observed a concentration-dependent pattern in which doses around the DC₅₀ appeared to increase CDK12 abundance after 24 h, whereas higher concentrations reduced it (Figure S5). As the conjugate delivers the degrader into cells more slowly than the free compound given the multi-step process, HER2-positive cells treated with Tz-SS2009 may experience lower intracellular degrader exposure. One possible explanation is that CDK12 is initially stabilized upon engagement of the complex, followed by its depletion at higher degrader concentrations as CCNK is degraded.

Beyond CCNK depletion, several protein-level changes consistent with biological processes previously associated with CDK12/CCNK function were seen (Figure 5d, Figure S20). These were found to include proteins involved in estrogen signaling (ESR1, PGR and GREB1),^90^ transcriptional regulation (PAGR1, CREB3L4 and MED29),^3^ genome maintenance (TERF2, CLSPN, RIF1 and DNA2)^91^, mitotic progression (PSRC1 and SAPCD2) and components of the HER2-regulated MAPK and mTOR pathways (DUSP4, SPRY4 and DEPTOR).^92,93^ The transcription-related proteins were typically downregulated, whereas the DNA repair proteins changed in both directions. The coordinated reduction of the DNA damage response proteins characteristic of CDK12 loss was not evident at 24 h. DEPTOR was an exception, in that it responded to Tz alone in the opposite direction to Tz-SS2009. Though efforts were taken to capture the proteomic changes that were related to degradation through the 24 h timepoint, these changes could also be consistent with broadly cytotoxic effects.

Whereas it had been challenging to distinguish phenotypic effects arising from Tz treatment and subsequent HER2 blockade, whole proteome analysis gave a clearer picture. Trastuzumab alone affected four proteins in BT-474 and none in SK-BR-3 relative to buffer, whereas direct comparison of Tz-SS2009 with Tz in BT-474 (isolating the contribution of the delivered degrader) changed 72 proteins (Figure S19). This substantially broader proteomic response to Tz-SS2009 than to Tz alone supports additional biological effects associated with effective delivery of the CCNK degrader, consistent with the differences in E_max_ observed between Tz-SS2009 and Tz in the proliferation assays (Figure S16).

In contrast, Tz-SS2009 produced neither CCNK depletion nor widespread proteomic remodeling in HER2-negative MDA-MB-231 cells (Figure 5a). To establish whether this reflects an absence of payload delivery rather than insensitivity of MDA-MB-231 to the degrader itself, we compared the conjugate with free SAS1001 in the same line. Unconjugated SAS1001 at 10 nM killed MDA-MB-231 cells within 24 h, to the extent that proteomic analysis was not possible (Figure S7). At DAR 4 (an average of four degraders per antibody), 10 nM Tz-SS2009 carries 40 nM of the same payload, yet these cells remained healthy and had the greatest proteome depth of any condition analyzed (7,824 protein groups; Figure S19), consistent with an unperturbed sample.

Finally, the non-targeting DAC b12-SS2009 did not affect any protein in any line (Figure 5a). Collectively, these data indicate that Tz-SS2009 induces selective depletion of CCNK in HER2-positive cells and produces a targeted set of early downstream proteomic changes consistent with known CDK12/CCNK biology.

## Discussion

Degraders acting through catalytic mechanisms can achieve substantial target depletion even at low exposure. Where the target is broadly expressed and essential, such as CCNK, indiscriminate degradation can occur in both healthy tissue and tumor, narrowing the therapeutic window. Selective delivery through an antibody conjugate offers a way to preserve potency while limiting exposure to antigen-negative tissue. Molecular glues are especially tractable payloads in this setting, as their smaller size and generally lower hydrophobicity relative to PROTACs may favor ease of conjugation and reduce DAC aggregation. Here we have developed a Tz-based DAC for antibody-directed delivery of a CCNK molecular glue degrader to HER2-positive cancer cells, with the aim of disrupting the CDK12/CCNK complex.

Whereas unconjugated SAS1001 degraded CCNK irrespective of HER2 status, conjugation to Tz yielded a DAR 4 DAC that internalized preferentially in HER2-positive cells and redistributed both CCNK degradation and antiproliferative activity towards them, with 51-fold greater CCNK depletion in BT-474 than in MDA-MB-231 at 10 nM (Figure 4d; Figure S15). Comparison with the isotype control b12-SS2009 showed this shift to be antigen-dependent. Residual activity at higher concentrations may reflect low-level antigen-independent exposure (such as non-specific uptake, premature extracellular payload release or linker instability), the consequences of which are amplified by the catalytic mechanism of molecular glue degraders. Proteomic analysis supported on-target engagement and showed that the selectivity extends beyond CCNK to downstream proteins, although further studies will be required to determine the mechanistic basis of individual pathway alterations. Cellular selectivity can therefore be imparted to a DDB1-CUL4-dependent CCNK molecular glue degrader without altering the underlying degradation mechanism, extending antibody-directed degradation beyond the more widely explored CRBN-based DACs.

Tz-SS2009 differs from the single reported non-CRBN molecular glue conjugate in two respects: it is delivered on a full-length IgG conjugated through a protease-cleavable linker rather than an affibody covalently crosslinked to the antigen,^64^ and its linker releases the unmodified degrader upon cleavage by CatB (Figure S9). Thus, Tz-SS2009 enables CCNK degradation preferentially in HER2-positive cells via a conventional antibody-drug conjugate structure. The payload is an optimized DS18-derived glue approximately 14-fold more potent than CR8 in the same reporter assay (Figure S3b), although the two conjugates have not been compared directly. The patent-disclosed CCNK molecular glue degrader conjugates (WO2025045758A1) carry pyrazolo[1,5-*a*][1,3,5]triazine payloads, a chemotype distinct from the purine series examined here.^66^

Our findings demonstrate that a structurally characterized, non-CRBN molecular glue scaffold can be adapted for conventional antibody conjugation through structure-guided attachment of a traceless, protease-cleavable linker, while retaining CCNK degradation activity and achieving preferential effects in antigen-positive cells. We anticipate that similar approaches could be extended to additional molecular glue scaffolds and tumor-associated antigens, although further studies will be required to determine their generalizability and potential to improve the therapeutic index in vivo.

## Limitations

Despite an average DAR of 4, the degradation potency of Tz-SS2009 was lower than that of the unconjugated degrader, on a nominal payload concentration basis. This observation suggests that processes such as receptor-mediated uptake, endolysosomal trafficking, linker cleavage and intracellular payload release limit the overall cellular effect.^38,94^ One potential factor may be sequestration of the released degrader in the acidic lysosome,^95^ through protonation of the basic piperazine moiety, which would reduce membrane permeability of SAS1001 and potentially limit the fraction able to reach the nucleus.^96^ Possible avenues to mitigate this include masking strategies, alternative attachment handles,^95,97^ or linkers that can release payloads extracellularly in the tumor microenvironment.^98^

Selection of an appropriate assay duration was important for evaluating DAC-mediated antiproliferative activity. As DACs require multiple processes to occur prior to target degradation and phenotypic response compared to small molecules, short-term viability assays may underestimate their activity. Although internalization occurred rapidly (Figure 3b, 3c), initial 3-and 5-day endpoint assays demonstrated a pronounced delay between treatment with Tz-SS2009 and growth inhibition, hampering reliable IC_50_ calculations. This observation is consistent with the delayed phenotypic effects expected following CCNK degradation and subsequent perturbation of cell cycle progression. All subsequent endpoint DAC cell proliferation assays, apart from MCF-7, were performed at 9 days. MCF-7 cells did not tolerate a 9-day incubation, including the non-treated control. As a result, 7-day assays were performed. While we performed treatments singly, subsequent in vitro studies should consider re-dosing and media replacement to avoid the impact of nutrient depletion.

Finally, while this work demonstrates the cellular efficacy of the DAC approach for delivering a highly potent CCNK molecular glue degrader to HER2-positive cells, the therapeutic index remains to be established *in vivo* and in the clinic. Clinical experience with DACs is limited, with the two GSPT1-targeting conjugates having both been terminated.^54,55^ Further evaluation in xenograft or patient-derived tumor models would allow exploration of whether the preferential cellular activity observed here translates into improved tumor selectivity, target engagement and tolerability.

## Resource availability

Further information and requests for resources and reagents should be directed to and will be fulfilled by the lead contact, Edward W. Tate. All unique/stable reagents generated in this study are available from the lead contact with a completed materials transfer agreement. Mass spectrometry proteomics data and R scripts are available from the lead contact on request.

## Acknowledgements

This work was supported by the Engineering and Physical Sciences Research Council (EPSRC studentship 2753945 to S.A.S.), co-funded by ADC Therapeutics. We thank ADC Therapeutics for their generous support, both for co-funding S.A.S.’s studentship and for hosting S.A.S. in their laboratories, where DAC conjugation and characterization were performed. We thank Matt White and Daisy Williams for proofreading and for valuable discussions on proteomic analysis. We also thank Edward Bartlett for laboratory management and ordering, Erika Bernardini for assistance with cell culture, James Zhang for assistance with biological assays, and Thomas Burden for synthetic chemistry support. We are grateful to Efthymios Gavriil, Thomas Jackson, Chloé Freyermuth and Anne Schuhmacher for helpful discussions. We thank Matthew Bird for discussions on ADCs.

## Contributions

S.A.S., A.B.P., and E.W.T. conceived the project. E.W.T. acquired funding. S.A.S. synthesized and characterized the degrader and linker-payload, performed the biological characterization and data analysis, prepared the figures, and wrote the manuscript. A.C. performed the scale-up and characterization of the DACs. L.C.H. performed the CCNK_eGFP_ reporter and TR-FRET assays. D.L. performed the molecular docking. J.L. ran the proteomic samples. A.M. assisted in small molecule characterization. L.N.C.R. (cathepsin B assay), J.W.H. (proteomics), J.W. (biological assays), V.S. (mass spectrometry), T.M. (preparative HPLC and data processing), and J.L.G. (statistical analysis) provided training and advice. N.V. and P.W.H. contributed synthesis suggestions. A.B.P., A.C., P.F., and E.W.T. supervised the work, and Z.K. supervised L.C.H. S.A.S., A.B.P., A.C., P.F., D.L., J.W., J.L.G., Z.K and E.W.T. reviewed and edited the manuscript. All authors approved the manuscript.

## Conflict of Interests

The authors do not have any conflicts of interest to disclose.

## Declaration of Funding

Engineering and Physical Sciences Research Council (2753945; 2753246; 2741582); Swiss National Science Foundation (P500-2_235357); European Commission (101282234); Cancer Research UK (DRCNPG-Nov21\100001; SEBPCTA-2024/100003); Medical Research Council (MR/X502959/1)

## Use of artificial intelligence

During the preparation of this work, the authors used Microsoft Copilot and Claude Opus 5 in order to suggest edits to text for concision and language clarity. Claude Opus 5 was also used to assist in generation of the data visualization scripts for proteomics. The final code was verified and executed by the authors. After using these tools, the authors thoroughly reviewed and edited the content as needed and take full responsibility for the content of the publication.

## Methods

### Cell lines

Cells were obtained from the American Type Culture Collection (ATCC). Cells were thawed and subcultured at least twice before their use in assays. Cells were regularly tested for mycoplasma contamination. Cells were maintained at 37 °C and 5% CO_2_ in a humidified atmosphere. BT-474 (HTB-20, ATCC) and MDA-MB-231 (HTB-26, ATCC) were cultured in high-glucose Dulbecco’s modified Eagle medium (DMEM) supplemented with GlutaMAX (61965059, Gibco) with 10% fetal bovine serum (FBS, A5256701, Gibco). MCF-7 (HTB-22, ATCC) were cultured in DMEM low glucose, pyruvate medium (11885084, Gibco) with 10% FBS. SK-BR-3 cells (HTB-30, ATCC) were cultured in McCoy’s 5a medium (16600082, Gibco) with 10% FBS. HEK293T Cas9 CCNK cells were cultured in DMEM high glucose (4.5 g/L with stable glutamine; BioConcept AG, Cat. No. 1-26F50-I), supplemented with 10% FBS (Cytiva, Cat. No. SH30088.03HI) and 1% penicillin–streptomycin (BioConcept, Cat. No. 4-01F00-H).

Cells were subcultured by washing with Dulbecco’s PBS (DPBS, D8537, Sigma-Aldrich) and detached by 0.25% trypsin-EDTA (T4049, Sigma-Aldrich).

### Molecular docking

The crystal structure of the CCNK/CDK12/DDB1 complex (PDB:8BU2) was visualized in ChimeraX (v1.12). The structure was prepared for docking using Schrodinger’s Maestro (v14.8) by filling the missing side chains, optimizing H-bond assignments and minimizing and deleting waters. Receptor docking grid was defined around the RVQ ligand within PDB:8BU2 by using its centroid as the grid center. SAS1001 ligand was prepared using Maestro’s LigPrep function, allowing for ionization, desalting and tautomer generation while retaining specified chiralities. The docking was performed via Glide within Maestro, with flexible ligand sampling and standard precision docking. The top ranked ligand was visualized with ChimeraX and 2D ligand interaction plots were obtained from Maestro.

### Cell proliferation assays

Cells were seeded in a 96-well flat-bottomed plate at 1000-5000 cells per well (9-day DAC assay) or 5000-10000 cells per well (5-day degrader assay). The seeding density was chosen dependent on the doubling time of the cell lines. Cells were treated 24 h post-seeding with compounds and controls (positive: puromycin (2 µg/mL, A1113803, Gibco); negative: DMSO 0.1% (degrader assay) or not treated (DAC assay)) in technical triplicates (technical duplicate for BT-474 DAC assays). The plates were incubated for 5 days (degrader assay) or 7/9 days (DAC assay). MTS assay solution (Promega Cell Titer 96 Aqueous G3581) was added (1:4), and the plates incubated for 4 h before absorbance detection at 490 nm using the CLARIOstar Plus plate reader (BMG LABTECH). The values were normalized by defining 0% as the arithmetic mean of the absorbance of the puromycin positive control and 100% as the arithmetic mean of the absorbance of the DMSO/non treated negative control. Individual dose-response curves were made for each biological replicate, and the IC_50_ was reported as the geometric mean. All data analysis was performed on GraphPad Prism (11.0.2).

### Degradation assays

6-well plates were seeded at 300,000 – 600,000 cells per well, dependent on cell doubling time. The cells were allowed to adhere for 24 h, then treated with compounds. The plates were incubated at 37 °C with 5% CO_2_ for 24 hours (SAS1001) or 72 h (DACs). Cells were harvested and used as in the western blot protocol below.

### Western blot analysis

Cells were washed three times with ice-cold DPBS. Ice-cold RIPA lysis buffer (R0278, Sigma-Aldrich) was freshly combined with cOmplete^TM^ EDTA-free protease inhibitor cocktail solution (11873580001, Roche) and added to the cells. The inhibitor stock was prepared by dissolving one tablet in 1 mL ddH₂O and added to RIPA buffer in a 1:50 dilution (20 µL stock per 980 µL RIPA lysis buffer). Following centrifugation (4 °C, 20 min, 17,000 x g), the supernatant was collected, and the protein concentration was determined by DC protein assay (5000112, Bio-Rad). The normalized lysate (1 mg/mL, 13 µg/well) was combined with 4x Laemmli buffer (1610747, Bio-Rad) containing 10% β-mercaptoethanol and denatured at 90 °C for 10 min. The samples were loaded onto a precast 4-20% SDS-PAGE gel (456102, Bio-Rad) and separated at 124 V for 1 h 15 min. The proteins were transferred to a nitrocellulose membrane using Trans-Blot Turbo Transfer System (Bio-Rad, Bio-Rad Mixed MW protocol, 10 min). The membranes were cut according to molecular weight and blocked for 2 h at RT in 5% solution of skim milk in tris-buffered saline with 0.1% Tween-20 (TBS-T). The membranes were incubated separately overnight at 4 °C with the primary antibody, diluted in 5% BSA in TBS-T (CDK12, 1:1000; CCNK, 1:2000; γH2AX, 1:1000; β-actin, 1:1000; GAPDH, 1:2500; CatB, 1:1000) or 5% skim milk in TBS-T (HER2, 1:1000). Following three washes for 5 min with TBS-T, the membranes were incubated with fluorescent IRDye 800CW Goat anti-rabbit IgG secondary antibodies (CDK12, CCNK, γH2AX) or IRDye 680RD anti-mouse (β-actin) as 1:10,000 dilutions in 5% BSA in TBS-T for 1 h at RT. Proteins were visualized on the LI-COR Biosciences Odyssey M system using the 700 and 800 nm channels.

| **Antibody** | **Supplier** | **Product code** |
|---|---|---|
| Anti-CCNK | Bethyl Laboratories (Thermofisher) | A301-939A |
| Anti-CDK12 | Cell Signaling Technologies | 11973 |
| Anti-GAPDH | Abcam | Ab9485 |
| Anti-β-actin | Abcam | Ab8226 |
| Anti-γH2AX | Cell Signaling Technologies | 9718 |
| Anti-cathepsin B | Cell Signaling Technologies | 31718S |
| Anti-HER2 | Cell Signaling Technologies | 2165S |
| IRDye 800CW Goat anti-rabbit IgG secondary | LI-CORbio | 926-32211 |
| IRDye 680RD Goat anti-mouse IgG secondary | LI-CORbio | 926-68070 |

### Reporter line generation

HEK293T-Cas9 cells expressing a CCNK (AA1–270)–eGFP degradation reporter (HEK293T-CCNK.Cilantro) were generated by lentiviral transduction. The reporter construct was expressed from the Cilantro 2 vector (Addgene, plasmid #74450), which additionally co-expresses mCherry as an internal reference. Lentiviral particles were produced in HEK293T cells by co-transfection of the reporter plasmid with the packaging plasmid psPAX2 and the envelope plasmid pVSV-G using TransIT-LT1 transfection reagent (Mirus Bio, Cat. No. MIR 2300). Virus-containing supernatants were collected 48 h after transfection and used to transduce HEK293T-Cas9 cells by spinfection, following previously described procedures.^16^

### Reporter-gene assay

Cells were seeded into 96-well plates at a density of 1 × 10⁵ cells per well in 150 µL of complete culture medium. Plates were prepared for samples to be measured in three technical replicates. Cells were treated with CR8 (3605, Tocris), SAS1001, or SR-4835 (39158, Cayman Chemical) over a concentration range of 9.6 pM to 1 µM for CR8 and SR-4835, and 1 pM to 1 µM for SAS1001. Where indicated, cells were pretreated with either MLN4924 (HY-70062, MedChemExpress) or TAK243 (T16974, TargetMol) at a final concentration of 1 µM for 1 h before addition of the respective CDK inhibitor. All compounds were dispensed using a TECAN Duo Digital Dispenser. Following compound addition, cells were incubated for 16 h before sample preparation for flow cytometric analysis. For conditions involving MLN4924 or TAK243 pretreatment, the total inhibitor exposure time was 17 h. Following compound treatment, culture medium was carefully aspirated, and cells were detached by adding 20 µL of 0.05% trypsin–EDTA solution (5-51F00-H, BioConcept) per well, followed by incubation at 37 °C for 3–5 min. Trypsinized cells were subsequently resuspended in 140 µL of pre-cooled FACS buffer consisting of phosphate-buffered saline (PBS) supplemented with 2% FBS. Cell suspensions were maintained on ice during transport to the flow cytometry facility. Flow cytometric measurements were performed using an Attune CytPix Flow Cytometer (Invitrogen, Thermo Fisher Scientific). FlowJo (BD Biosciences, 10.10.1) was used to determine the mean fluorescent intensity of the eGFP and mCherry signal for mCherry positive and round cells (Figure S3c). This was normalized to the triplicate DMSO controls.

### TR-FRET assay

Recombinant CDK12–CCNK and DDB1 proteins were expressed, purified, and labeled as previously described.^12^ Briefly, CCNK contained a C-terminal AviTag for BirA-mediated biotinylation, whereas DDB1 contained an N-terminal SpyTag for conjugation to Alexa Fluor 488-labeled SpyCatcher. For TR-FRET-based compound evaluation, biotinylated CDK12–CCNK (50 nM), Alexa Fluor 488-labeled DDB1 (100 nM), and terbium-labeled streptavidin (4 nM; LanthaScreen™ Tb-Streptavidin, Invitrogen, Thermo Fisher Scientific) were combined in TR-FRET assay buffer containing 50 mM HEPES (pH 7.4), 150 mM NaCl, 0.005% Tween 20, 0.5% DMSO, 1 mM tris(2-carboxyethyl) phosphine (TCEP), and 2 mM ethylenediaminetetraacetic acid (EDTA). The reaction mixture was dispensed into white 384-well microplates (Greiner Bio-One, Cat. No. 784075) at a volume of 6 µL per well. Compounds were subsequently dispensed from DMSO stock solutions using a D300 Digital Dispenser (Tecan). The compounds SAS1001 and CR8 (Tocris Bioscience, Cat. No. 3605) were tested using 11-point concentration series ranging from 0.001 to 2.013 µM. TR-FRET measurements were carried out using a PHERAstar FSX microplate reader (BMG Labtech) equipped with a 337-520-490 filter set. The TR-FRET signal was obtained through calculating the ratio of 520 nm to 490 nm fluorescence, and Prism Version 11.1.0 (GraphPad) was used for further data analysis. The curves resulting from plotting the TR-FRET against compound concentration were fitted as described previously to determine the half-maximal effective concentrations (EC_50_).^12^

### Ubiquitin-proteasome system inhibition assay

The method for degradation assay was followed, with an additional pre-treatment step prior to addition of compounds. Appropriate wells were pretreated with MLN4924 (10 µM), bortezomib (10 µM) or DMSO for 1 h, with incubation at 37 °C with 5% CO_2_.

### FabFluor internalization assay

SK-BR-3 (7000 cells per well) and MDA-MB-231 (4000 cells per well) were seeded in a 96-well plate and incubated overnight. A freshly prepared 2x solution of biologic compound (100 nM) and Sartorius FabFluor dye (300 nM, 4722, Sartorius) was incubated for 15 min, before being added to the plate in technical triplicate. The plate was incubated in an Incucyte S3 for 24 h, with scans every 1 h. Images were taken in the phase and red channels, with an acquisition time of 400 ms. A 10x objective on the S3/SX1 G/R optical module was used. Following data acquisition, analysis was performed using the Incucyte software. Phase was analyzed by AI Confluence segmentation with no cleanup. Filters were not set. Red was analyzed by Surface Fit segmentation with a Threshold (RCU) of 0.13 with Edge Split On with no cleanup. Filters were set for a maximum mean intensity of 0.9. Red area per image was normalized to phase (confluence) area per image following background subtraction, followed by plotting in GraphPad Prism (11.0.2).

### Whole proteome – sample preparation

SK-BR-3, BT-474 and MDA-MB-231 cells were seeded in a 12-well plate at 200,000 cells per well and incubated overnight. Cells were treated with SAS1001 (10 nM, 2 h, 6 h or 24 h), Tz-SS2009 (10 nM, 24 h), b12-SS2009 (10 nM, 24 h), Tz (10 nM, 24 h) or an equivalent volume of formulation buffer in quadruplicate. Cells were washed three times with ice-cold DPBS. Ice-cold RIPA lysis buffer (R0278, Sigma-Aldrich) was freshly combined with cOmpleteTM EDTA-free protease inhibitor cocktail solution (11873580001, Roche) and Benzonase endonuclease (1:1000, E1014-25KU, Millipore) and added to the cells. The inhibitor stock was prepared by dissolving one tablet in 1 mL ddH₂O and added to RIPA buffer in a 1:50 dilution (20 µL stock per 980 µL RIPA lysis buffer). Following centrifugation (4 °C, 20 min, 17,000 x g), the supernatant was collected, and the protein concentration of the lysate was determined by DC protein assay (5000112, Bio-Rad). The samples were diluted to 1 mg/mL using lysis buffer. 10 µg of protein lysate was taken forward for reduction and alkylation using 0.5 volumes each of tris(2-carboxyethyl)phosphine (40 mM in 50 mM HEPES, pH 8) and chloroacetamide (160 mM in 50 mM HEPES, pH 8). The samples were shaken for 10 min at RT. SP4 precipitation was carried out by combination of reduced/alkylated protein mixture with acetonitrile in a 1:4 ratio.^99^ Following centrifugation at 16,000 x g for 5 min, the pellet was washed 3 times with 80% ethanol in a 1:10 ratio, with 2-min centrifugation cycles at 16,000 x g in between washes. The supernatant was removed, and the pellet left to dry for 10 min prior to resuspension in trypsin (14 ng/µL in 50 mM HEPES, pH 8). The samples in digestion buffer were incubated at 37 °C overnight, with shaking at 600 rpm. Peptides were collected as the supernatant following 2 min centrifugation at 16,000 x g and concentrations quantified by Pierce Quantitative Fluorometric Peptide Assay (23290, Thermo Scientific). Peptides were normalized to 500 ng peptide in 50 µL in LC-MS grade H_2_O with 0.1% formic acid (FA).

### Whole proteome – Evosep – timsTOF – DIA

Peptides were analyzed by nanoLC-MS/MS using an Evosep One (Evosep) coupled with a timsTOF HT (Bruker) equipped with an 8 cm x 150 µm, 1.5 µm analytical column (Evosep). 200 ng peptides were separated by the Evosep 60SPD workflow (Analytical solvents A: 0.1% FA and B: acetonitrile plus 0.1% FA). Column was held at 40 °C. Data were acquired in data-independent acquisition PASEF mode with the following settings: m/z range from 100 m/z to 1700 m/z, ion mobility range from 1/K0 = 1.30 to 0.85 Vs/cm^2^ using equal ion accumulation and ramp times in the dual TIMS analyzer of 100 ms each. Each cycle consisted of 8 PASEF ramps covering 21 mass steps each with 25 Da windows each with 2/3 non-overlapping ion mobility windows covering the 475 to 1000 m/z range and 0.85 and 1.26 Vs/cm^2^ ion mobility range. The collision energy was lowered as a function of increasing ion mobility from 59 eV at 1/K0 = 1.6 Vs/cm^2^ to 20 eV at 1/K0 = 0.6 Vs/cm^2^.

### DIA-NN processing

diaPASEF Bruker.d files were processed in DIA-NN (2.6.0 Academia)^100^ against the UniProt human reference proteome UP000005640 (downloaded 17 August 2026; 20,749 entries), with the DIA-NN contaminants database included. A spectral library was first generated from the data by library-free analysis with deep learning-based prediction of spectra, retention times and ion mobilities, and the runs were then searched against this library. Search parameters were: Trypsin/P with up to one missed cleavage; N-terminal methionine excision, cysteine carbamidomethylation, methionine oxidation and N-terminal acetylation, with a maximum of two variable modifications per precursor; peptide length 7–30 residues; precursor charge 1–4; precursor m/z 300–1800; fragment m/z 200–1800. MS1 accuracy, MS2 accuracy, calibration accuracy and scan window were set to 0, so that DIA-NN inferred them from the data. Match-between-runs was enabled. Scoring was performed on peptidoforms using cross-validated neural networks, protein inference was enabled with proteotypicity assessed at gene level, quantification used QuantUMS (precision), and cross-run normalization was RT-dependent. Output matrices were filtered by DIA-NN at 1% FDR.

### Perseus data analysis

The results from the DIA-NN processing were analyzed in Perseus (version 1.6.2.3). The matrix was uploaded and the intensity data transformed by log2(x). Rows were annotated to assign group replicates. A text filter was applied to remove contaminants (cRAP or CON_). The data was filtered based on valid values, with a requirement of at least 3 valid values in each group of replicates. Missing values were imputed from normal distribution with width 0.3 and down shift 1.8. Volcano plots were created, plotting the difference in protein intensities between treatment and vehicle was plotted in RStudio (2025.05.1 Build 513) with a two-sided Student’s t-test; p-values were corrected by the Benjamini–Hochberg procedure. Proteins were considered significant where adj. *p* < 0.05 and |log₂FC| > 1.

### Cathepsin B cleavage assay

A solution of Cathepsin B from human liver (Sigma-Aldrich, C8571-25UG) at 0.25 µM was prepared in activation buffer (30 mM DTT, 15 mM EDTA, pH 5.5) and incubated at 37 °C for 15 min. A batch of linker-payload SS2009 was prepared at 312.5 µM in DMSO then further dissolved at 125 µM in sample buffer (25 mM NaOAc, 1 mM EDTA, pH 5.5). Cleavage was initiated by mixing equal volumes of activated cathepsin B and linker-payload solution, resulting in a final enzyme-to-payload molar ratio of 1:500. A negative control was prepared by replacing the activated cathepsin B solution with an equal volume of activation buffer. Cleavage was monitored over 16 h by RP-HPLC and LC-MS, with samples collected at predetermined time points.

### Synthesis of b12-SS2009 DAR 4

b12 (23 mg, 13.2 mg/mL) in PBS was reduced with tris(2-carboxyethyl)phosphine (TCEP) (10 mM in H_2_O, 2.8 eq.) in presence of 5% of pH adjust buffer (0.5 M Tris, 25 mM EDTA, pH 8.0). The mixture was incubated at 22 °C for 1.5 h. Following that, SS2009 (10 mM in dimethylacetamide (DMA), 10 eq.) was added and incubated at 22 °C for 1.0 h. The crude conjugate was then analyzed via HIC-HPLC, PLRP-HPLC and LC-MS for DAR determination, and SEC-HPLC for monomeric content. The final conjugate was buffer exchanged (30 kDa MWCO, Sartorius Vivaspin) into formulation buffer (30 mM Histidine, 175 mM Sucrose, 0.02% PS20, pH 6.0) and filtered (Millex-GV 0.22 µm). The final conjugates were then analyzed by HIC-HPLC, PLRP-HPLC, LC-MS and SEC-HPLC.

### Synthesis of Tz-SS2009 DAR 4

Tz (23 mg, 9.49 mg/mL) in PBS was reduced with tris(2-carboxyethyl)phosphine (TCEP) (10 mM in H_2_O, 2.8 eq.) in presence of 5% of pH adjust buffer (0.5 M Tris, 25 mM EDTA, pH 8.0). The mixture was incubated at 22 °C for 1.5 h. Following that, SS2009 (10 mM in DMA, 10 eq.) was added and incubated at 22 °C for 1 h. The crude conjugate was then analyzed via HIC-HPLC, PLRP-HPLC and LC-MS for DAR determination, and SEC-HPLC for monomeric content. The final conjugate was buffer exchanged (30 kDa MWCO, Sartorius Vivaspin) into formulation buffer and filtered (Millex-GV 0.22 µm). The final conjugates were then analyzed by HIC-HPLC, PLRP-HPLC, LC-MS and SEC-HPLC.

### Quantification and statistical analysis

One replicate was removed from the 6 h SK-BR-3 proteomic analysis following sample preparation. The same was performed for one replicate of Tz-SS2009 in BT-474. Both samples were analyzed and excluded on the basis of experimental error and principal component analysis. The 24 h MDA-MB-231 proteomic experiment was excluded in full because cell loss left insufficient material. No other data were excluded.

Cell proliferation data is representative of one biological replicate, with mean ± SD from technical triplicates. Normalization and IC_50_ were calculated and displayed in GraphPad Prism 11.0.2 (four-parameter variable slope). IC₅₀ and DC₅₀ are geometric means back-transformed from the mean pIC₅₀ or pDC₅₀. pIC₅₀ and pDC₅₀ are mean ± SD calculated on the log scale. Where a curve could not be fitted for an individual replicate, the n of the fitted parameters is reported separately from the n contributing to endpoint measurements. Viability was normalized to the puromycin control (0%) and the vehicle or untreated control (100%) on the same plate. For the TR-FRET assay the bottom parameter was constrained to the no-compound control.

Western blot data is representative of three biological replicates performed independently. Band intensities were quantified in LI-COR Empiria Studio 3.2.0.186, normalized to loading control and then to the vehicle or untreated control on the same membrane, and log₂-transformed before analysis. Transformed values were compared by ordinary two-way ANOVA with Šídák’s correction (factors: treatment and concentration for Figure 4b; cell line and concentration for Figure 4d). Effects are reported as differences in log₂ abundance with 95% confidence intervals, back-transformed to fold differences. Statistical details pertaining to experiments can be found in figure captions. Degradation at 10 nM is 1 − normalized abundance at that single concentration and is not a fitted D_max_.

pIC₅₀ values were compared by ordinary two-way ANOVA with cell line and treatment as factors, using Type III sums of squares for the unbalanced design; the interaction was significant (F(2, 11) = 12.58, p = 0.0014). Pairwise comparisons used Šídák’s correction within five pre-specified families. Fold differences are back-transformed from differences in pIC₅₀.

Whole proteome analysis was performed as n=4, with exclusions as noted above. Analyses used log₂-transformed LFQ intensities exported from Perseus, filtered to require at least three valid values in each group and with remaining missing values imputed from a downshifted normal distribution (width 0.3, downshift 1.8); the exported matrix was complete and no further filtering or imputation was applied. Log₂ fold change is the difference between group means, and significance was assessed by two-sided two-sample Student’s t-test. Immunoglobulin-derived proteins were excluded before correction and are presented separately. p values were corrected by the Benjamini–Hochberg procedure within each contrast. Proteins were considered differentially abundant at adj. p < 0.05 and |log₂ fold change| > 1, marked by dashed lines on all volcano plots.

Statistical tests were selected according to the structure of each comparison. Adjusted values are labelled “adj. p” and unadjusted values “p”. Exact p values are given to two significant figures, or as p < 0.0001 below that threshold. Analyses were performed in GraphPad Prism 11.0.2 and R 4.5.0, with proteomic processing in DIA-NN 2.6.0 and Perseus 1.6.2.3.

Potency values (DC₅₀, IC₅₀, EC₅₀) and fold differences are reported to two significant figures. Standard deviations are given to one significant figure (two where the leading digit is 1), and pXC₅₀ values to the corresponding number of decimal places.

## Notes

### Competing Interest Statement

The authors have declared no competing interest.

