## Supplementary Information for "A molecular glue degrader-antibody conjugate for HER2-targeted cyclin K degradation"

#### Supporting figures

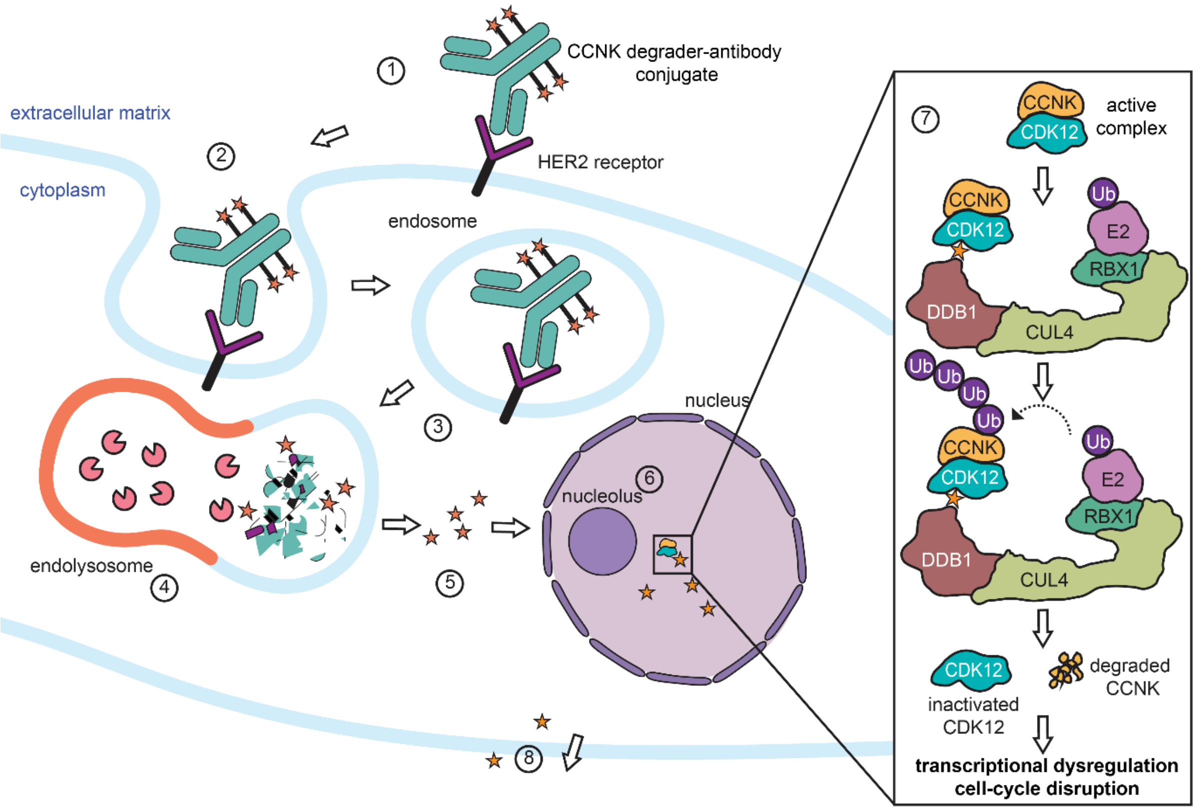


*Figure S1.* ***Proposed mechanism of antibody-directed CCNK degradation****. Following HER2-mediated binding (1) and internalization (2,3), lysosomal processing releases SAS1001 (4,5), which enters the nucleus (6) and promotes recruitment of the DDB1-CUL4 E3 ligase complex to CDK12/CCNK, resulting in CCNK ubiquitylation and degradation (7). Selective degradation of CCNK disrupts CDK12-dependent transcriptional programs and impairment of the cell-cycle. The released degrader also has the potential to diffuse out of the cell and into neighboring cells; the bystander effect (8), which was not assessed in this study.*

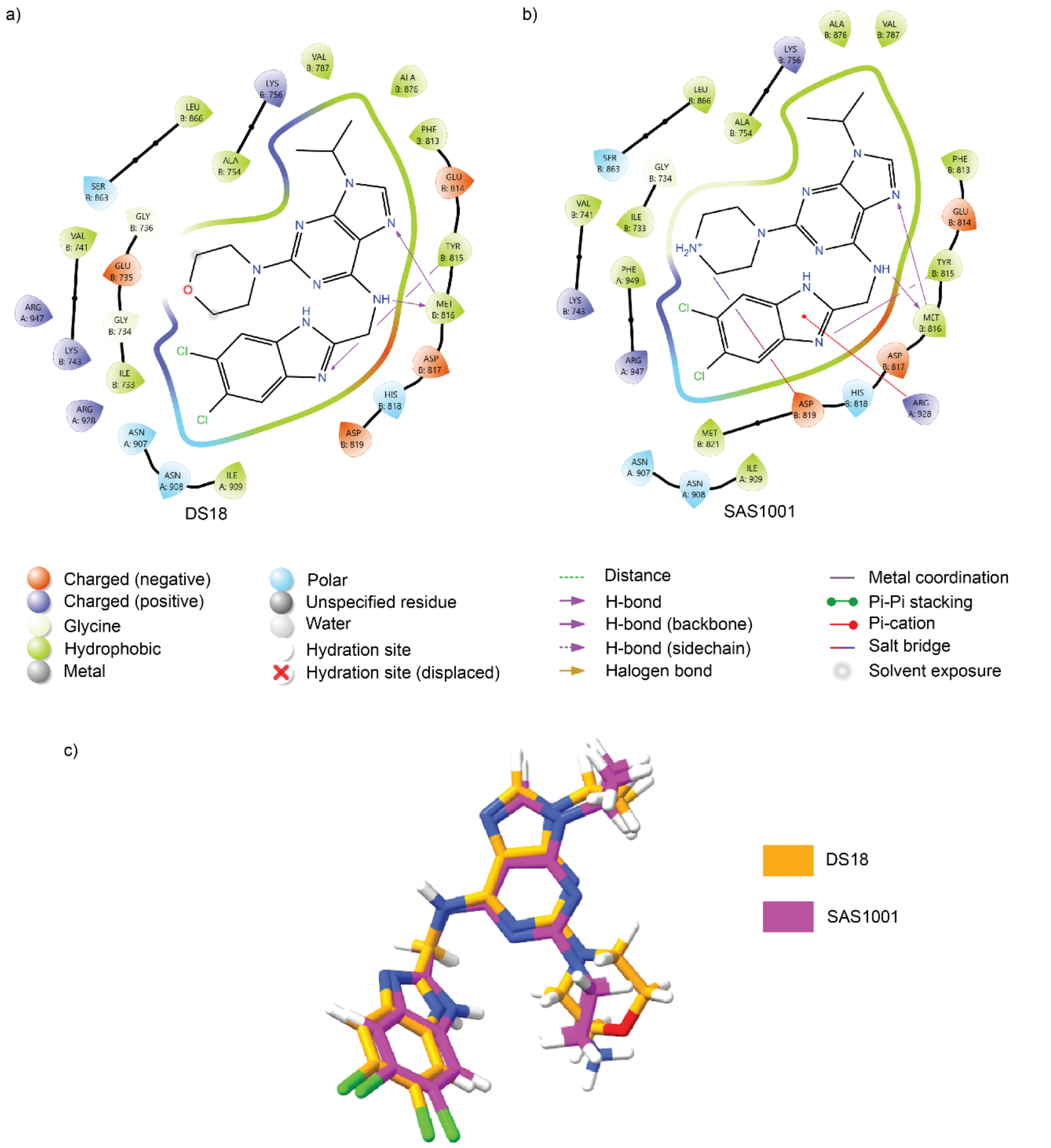


*Figure S2.* ***Structural comparison of DS18 and SAS1001 within the DDB1:CDK12:CCNK ternary complex. (a)*** *Two-dimensional interaction map of DS18 derived from the crystal structure of the DDB1:CDK12:CCNK:DS18 ternary complex (PDB: 8BU2).* ***(b)*** *Two-dimensional interaction map of SAS1001 docked into the 8BU2 ternary complex structure. SAS1001 retained the key hydrogen-bonding and hydrophobic interactions observed for DS18 and was additionally predicted to form a salt bridge with Asp819, providing a potential stabilizing electrostatic interaction.* ***(c)*** *Overlay of the crystallographic pose of DS18 (orange) and the docked pose of SAS1001 (magenta), demonstrating close alignment of the kinase-binding moieties and suggesting that linker modification is compatible with the binding mode adopted within the DDB1:CDK12:CCNK ternary complex.*

a)

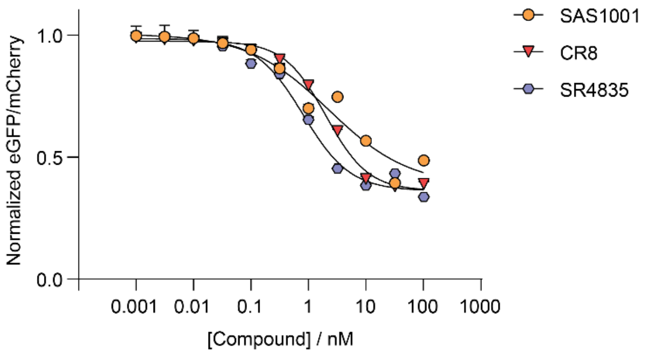


b)

| Compound | $DC_{50}$ (nM)[a] | $pDC_{50}$ ± SD[b] | n[c] | Literature $DC_{50}$ (nM)[d] | Difference to literature |
|---|---|---|---|---|---|
| SAS1001 | 0.917 | 9.0 ± 0.4 | 3 | - | - |
| CR8 | 12.6 | 7.9 ± 0.3 | 2 | 24 | 1.9 |
| SR-4835 | 6.03 | 8.22 ± 0.19 | 2 | 14 | 2.3 |
| DS18[e] | n.d. | - | - | 13 | - |

c)

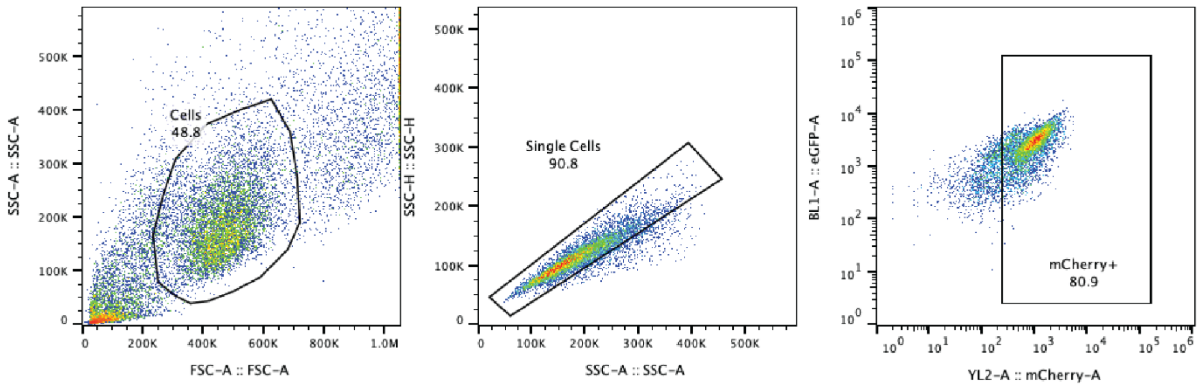


*Figure S3.* ***SAS1001 degrades a CCNK–eGFP reporter more potently than CR8 or SR-4835. (a)*** *Dose–response curves for SAS1001, CR8 and SR-4835 in HEK293T-CCNK.Cilantro reporter cells, expressing a CCNK(AA1–270)–eGFP fusion with mCherry as an internal reference. Cells were treated for 16 h over 9.6 pM to 1 µM (CR8, SR-4835) or 1 pM to 1 µM (SAS1001), and eGFP and mCherry mean fluorescence intensities were determined by flow cytometry in mCherry-positive single cells. Signal is expressed as eGFP/mCherry normalized to DMSO-treated controls on the same plate. Points are mean ± SD of three technical replicates; curves are four-parameter nonlinear regression fits.* ***(b)*** *Summary of reporter degradation parameters, with published values for the same reporter line shown for comparison.* ***(c)*** *Gating strategy: cells were gated on FSC-A/SSC-A, singlets on SSC-H/SSC-A, and analysis restricted to the mCherry-positive population.* [a] *Geometric mean, back-transformed from the mean $pDC_{50}$.* [b] *Arithmetic mean ± SD of $-\log_{10} DC_{50}$ [M] (mean and SD calculated on log-transformed values). SD is given only where $n > 1$.* [c] *Number of independent biological replicates.* [d] *Published $DC_{50}$ values in the same reporter line, from Kozicka et al.*[1] [e] *DS18 was not available for assay in parallel. n.d., not determined.*

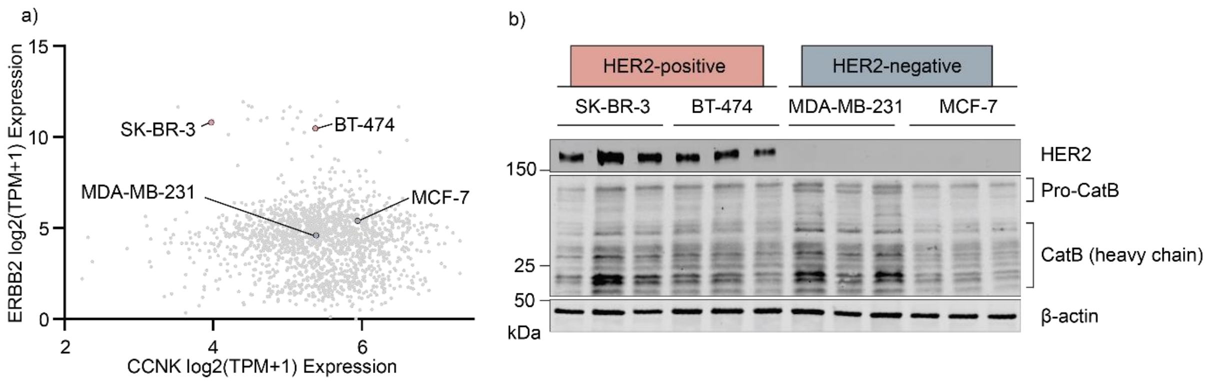


*Figure S4*. ***Selection and characterization of breast cancer cell line models. (a)*** *Relative HER2 and CCNK expression in BT-474, SK-BR-3, MDA-MB-231 and MCF-7 cells obtained from DepMap datasets.* ***(b)*** *Western blot confirming expression of HER2 and cathepsin B across the cell line panel. β-Actin was used as a loading control. Each individual lane represents a biological replicate.*

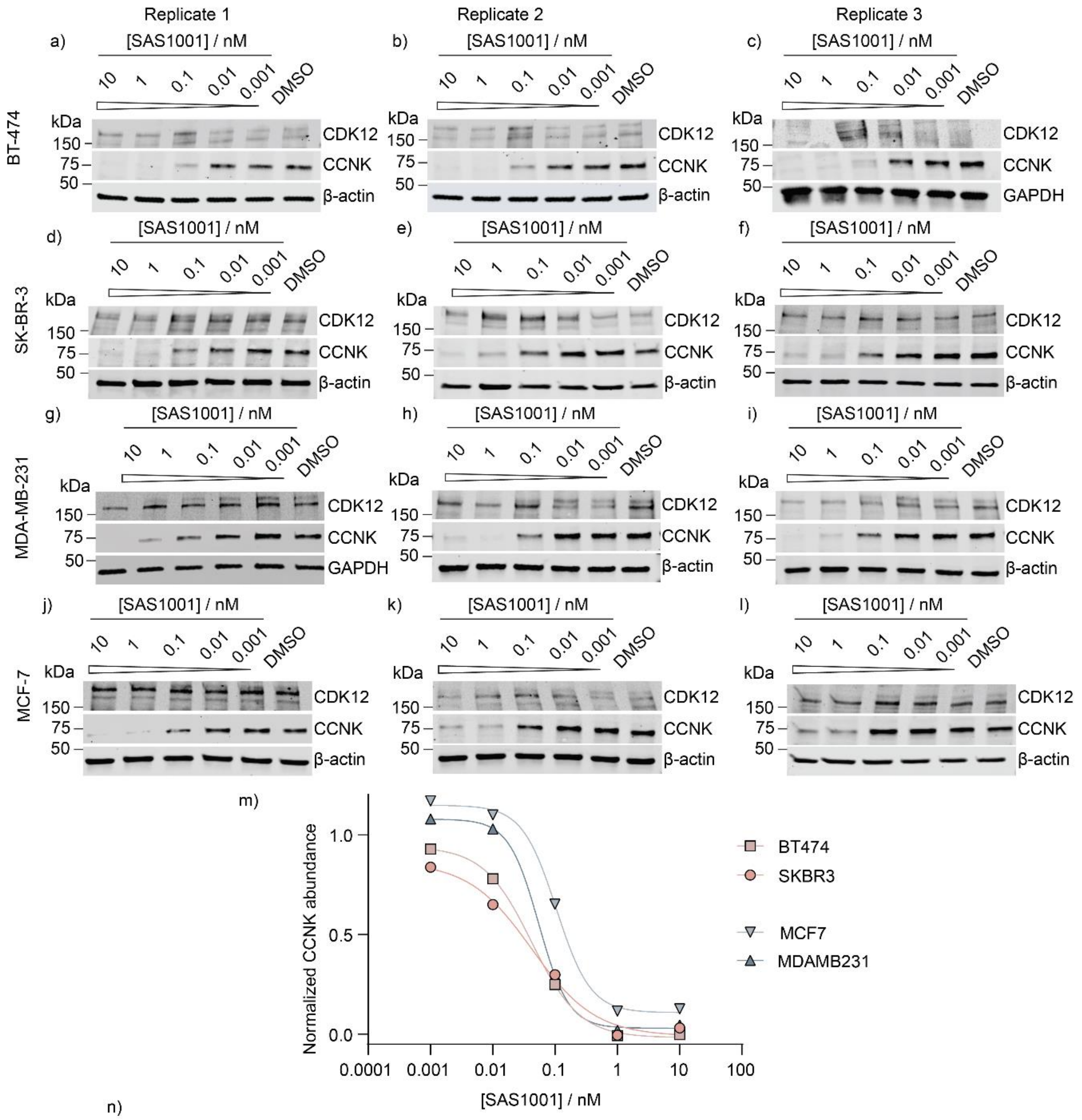


n)

| Cell line | $DC_{50}$ / pM [(a)] | $pDC_{50}$ ± SD [(b)] | $D_{10\ nM}$ / % [(c)] | n |
|---|---|---|---|---|
| BT-474 | 37 | 10.4 ± 0.2 | ≥ 99 | 3 |
| SK-BR-3 | 32 | 10.5 ± 0.6 | ≥ 99 | 3 |
| MCF-7 | 97 | 10.01 ± 0.08 | 83 | 3 |
| MDA-MB-231 | 69 | 10.2 ± 0.3 | ≥ 99 | 3 |

*Figure S5* ***Western blots of treatment by SAS1001.*** *Cells were seeded overnight and treated with SAS1001 or DMSO for 24 h prior to lysis. Membranes were probed for CDK12, CCNK and the loading control (β-actin or GAPDH, as indicated).* ***(a)-(c)****, BT-474 (HER2 positive);* ***(d)-(f)****, SK-BR-3 (HER2 positive);* ***(g)-(i)****, MDA-MB-231 (HER2 negative);* ***(j)-(l)****, MCF-7 (HER2 negative).* ***(m)*** *Densitometric quantification of CCNK abundance, normalized to loading control (β-actin or GAPDH, as indicated) and DMSO vehicle. Data are from a single representative biological replicate with a four-parameter fit nonlinear regression curve.* ***(n)*** *Summary of CCNK degradation parameters. Apparent $DC_{50}$, $pDC_{50}$ ± SD[b] and $D_{max}$[c] were derived from four-parameter fits of each of the three independent biological replicates.* [a] *Geometric mean of n = 3 (back-transformed from the mean $pDC_{50}$).* [b] *Arithmetic mean ± SD of $-\log_{10} DC_{50}$ [M]; mean and SD calculated on log-transformed values.* [c] *$D_{max}$ (1 – fitted plateau).*

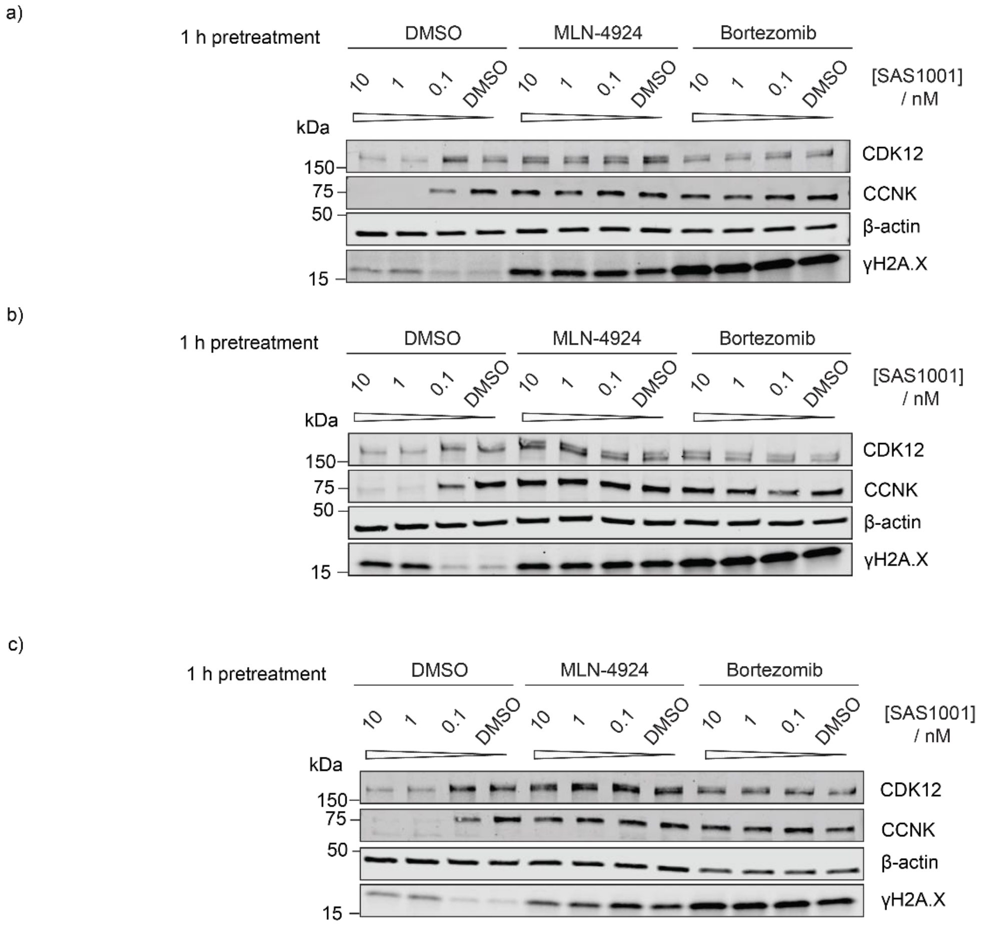


*Figure S6.* ***Rescue of CCNK degradation upon pretreatment with ubiquitin proteasome inhibitors*****. *(a-c)*** *SK-BR-3 cells were pretreated with bortezomib (10 µM), MLN-4924 (10 µM) or vehicle for 1 h before addition of SAS1001. Western blot analysis after 24 h demonstrates rescue of CCNK degradation following inhibition of the proteasome or Cullin neddylation. DNA-damage induction can be seen through the appearance of γH2AX in the top two concentrations in the DMSO pretreated lysates. Strong signal for γH2AX can also be seen in response to bortezomib and MLN-4924 pretreatment.*

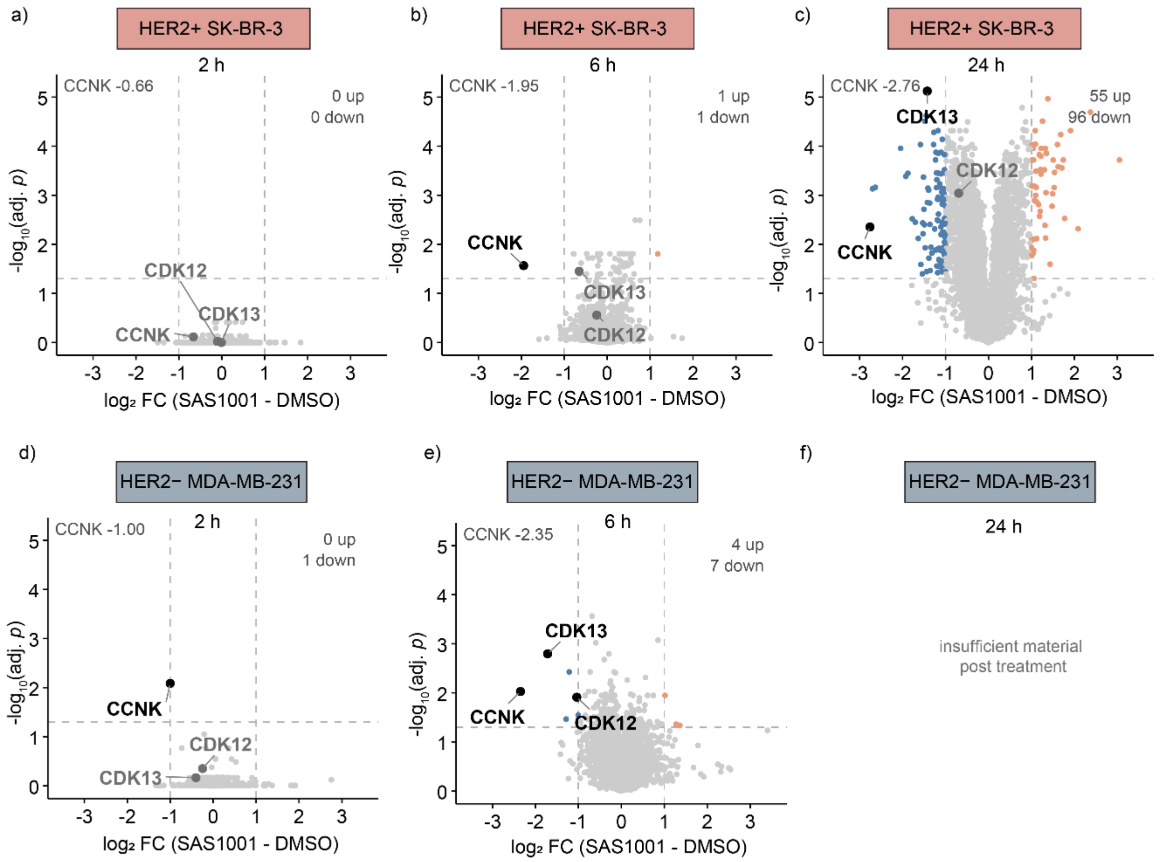


*Figure S7* ***CCNK is depleted by free SAS1001 with comparable efficiency in HER2-negative and HER2-positive cells****. SK-BR-3 (HER2+, top;* ***a-c****) and MDA-MB-231 (HER2−, bottom;* ***d-f****) were treated with SAS1001 (10 nM) or DMSO for 2, 6 or 24 h and analysed by label-free DIA proteomics (n = 4 per condition). Each duration was a separate experiment, processed and searched independently, and each is compared against its own matched DMSO control; the panels are independent comparisons rather than a single time course. The 24 h MDA-MB-231 experiment was performed but excluded* ***(f)****: cell loss at this timepoint yielded insufficient material for reliable quantification. Proteins passing* $|log_2FC| > 1$ *and adj.* $p < 0.05$ *are coloured by direction (orange, increased; blue, decreased) with dashed lines marking the thresholds, and CCNK, CDK12 and CDK13 are labelled throughout, in black where they pass both thresholds and grey where they do not. Significance was assessed by two-sided Student's t-test and corrected by the Benjamini–Hochberg procedure.*

| Cell line | $IC_{50}$ (nM)[a] | $pIC_{50}$ ± SD[b] | n |
|---|---|---|---|
| BT-474 | 1.0 | 8.99 ± 0.10 | 3 |
| SK-BR-3 | 2.2 | 8.66 ± 0.13 | 3 |
| MCF-7 | 14 | 7.86 ± 0.03 | 3 |
| MDA-MB-231 | 3.8 | 8.42 ± 0.09 | 3 |

*Figure S8.* ***Antiproliferative potency of SAS1001 across the breast cancer cell line panel.*** *IC50[a] and pIC50 ± SD[b] determined by MTS assay after 5 days treatment. [a] Geometric mean of n = 3 independent biological replicates (back-transformed from the mean pIC50). [b] Arithmetic mean ± SD of $-\log_{10}$ IC50 [M]; mean and SD calculated on log-transformed values.*

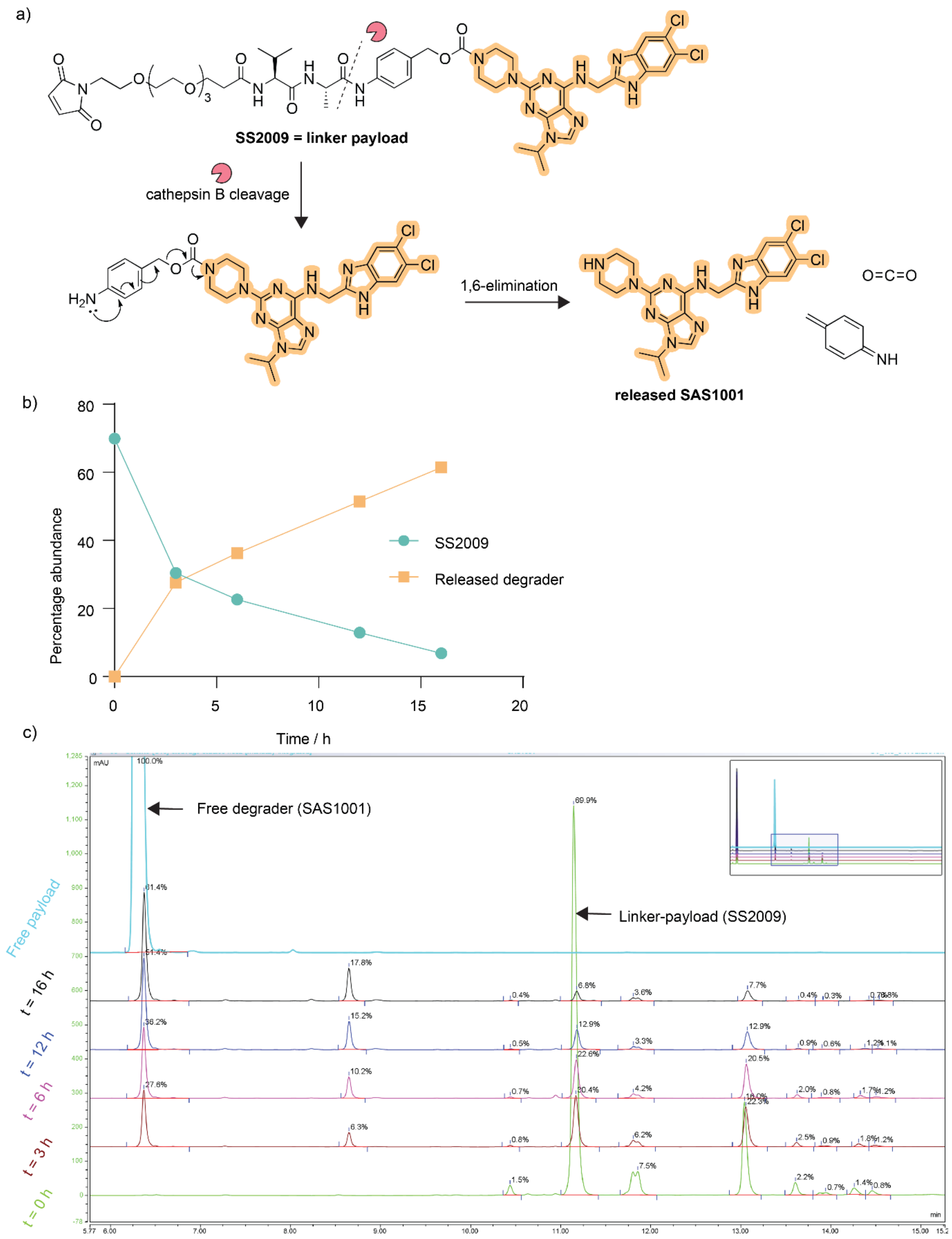


*Figure S9.* ***Cathepsin B cleavage assay. (a)*** *Val-Ala linkers release their payloads after the amide bond proximal to the p-aminobenzyl moiety is cleaved by cathepsin B. This leads to the formation of an unstable p-aminobenzyl fragment which releases the free degrader upon 1,6-elimination. Aza-quinone methide is also released, which is typically quenched by water in the cell, as well as carbon dioxide.* ***(b)*** *SAS1001 release over time versus disappearance of the linker-payload SS2009.* ***(c)*** *Time-course cathepsin B cleavage kinetics assay. SS2009) was treated with cathepsin B (from human liver) at 37 °C and followed by reverse-phase high-performance liquid chromatography (RP-HPLC) for 16 h.*

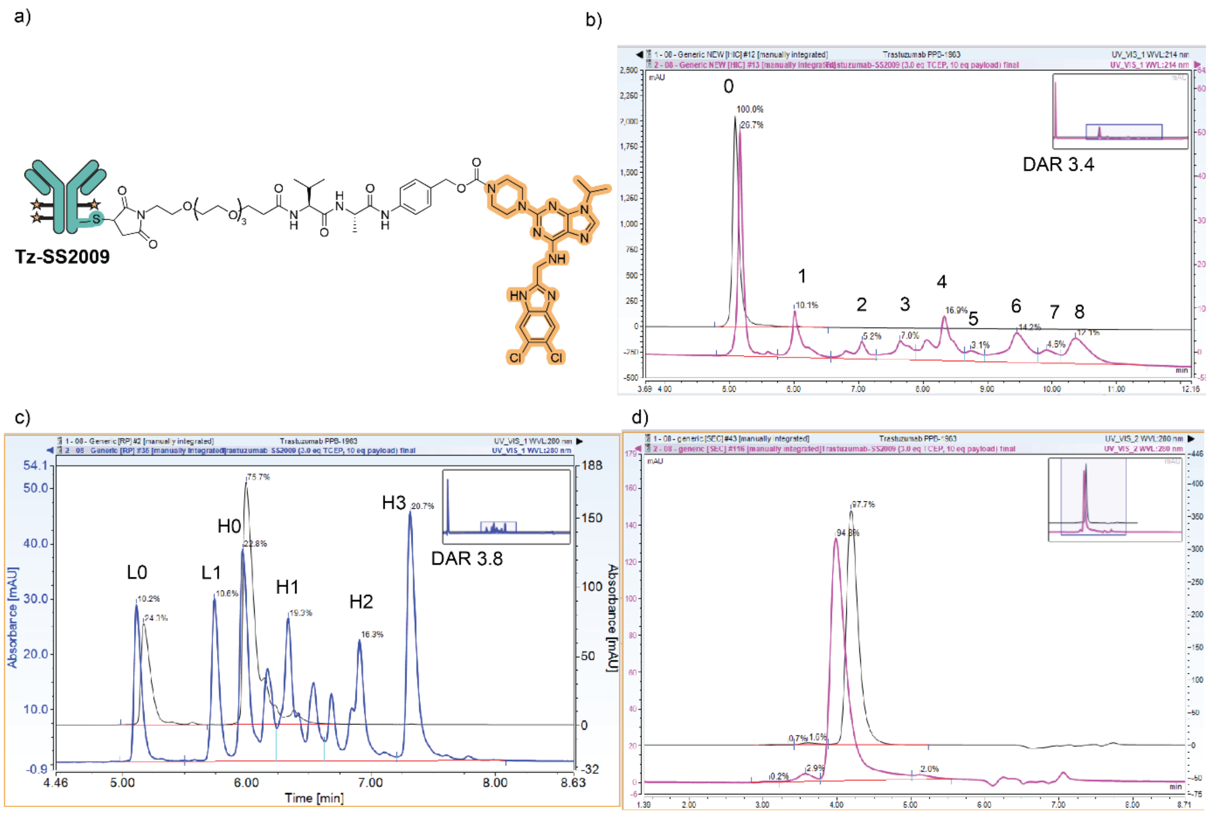


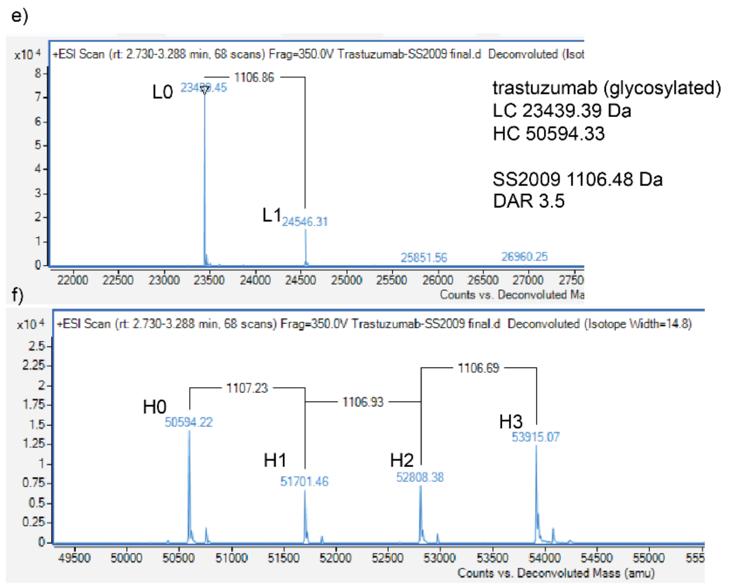


g)

| | Tz-SS2009 | |
|---|---|---|
| TCEP | 3 | eq |
| Input | 23 | mg |
| [P] | 6.63 | mg/mL |
| Volume | 2.65 | mL |
| Amount | 17.6 | mg |
| Yield | 74.2 | % |
| DAR (PLRP) | 3.8 | |
| DAR (HIC) | 3.4 | |
| DAR (LC-MS) | 3.5 | |
| Monomer (SEC) | 94.8 | % |
| Endotoxin | 0.046 | EU/mg |
| Batch | PPB-1985 | |
| MW | 149881 | Da |
| $EC_{280}$ | 251108 | |

*Figure S10. **Tz-SS2009 characterization.** **(a)** Representation of a ~DAR 4 Tz-SS2009 DAC. **(b)** Hydrophobic interaction chromatography (HIC) spectrum for trastuzumab (black) and Tz-SS2009 (pink). **(c)** PLRP spectrum for trastuzumab (black) and Tz-SS2009 (blue). **(d)** Size-exclusion chromatography (SEC) spectrum for trastuzumab (black) and Tz-SS2009 (pink). **(e)** LCMS spectrum for Tz-SS2009, depicting light chain conjugation. **(f)** LCMS spectrum for Tz-SS2009, depicting heavy chain conjugation. **(g)** Conjugation summary table for Tz-SS2009.*

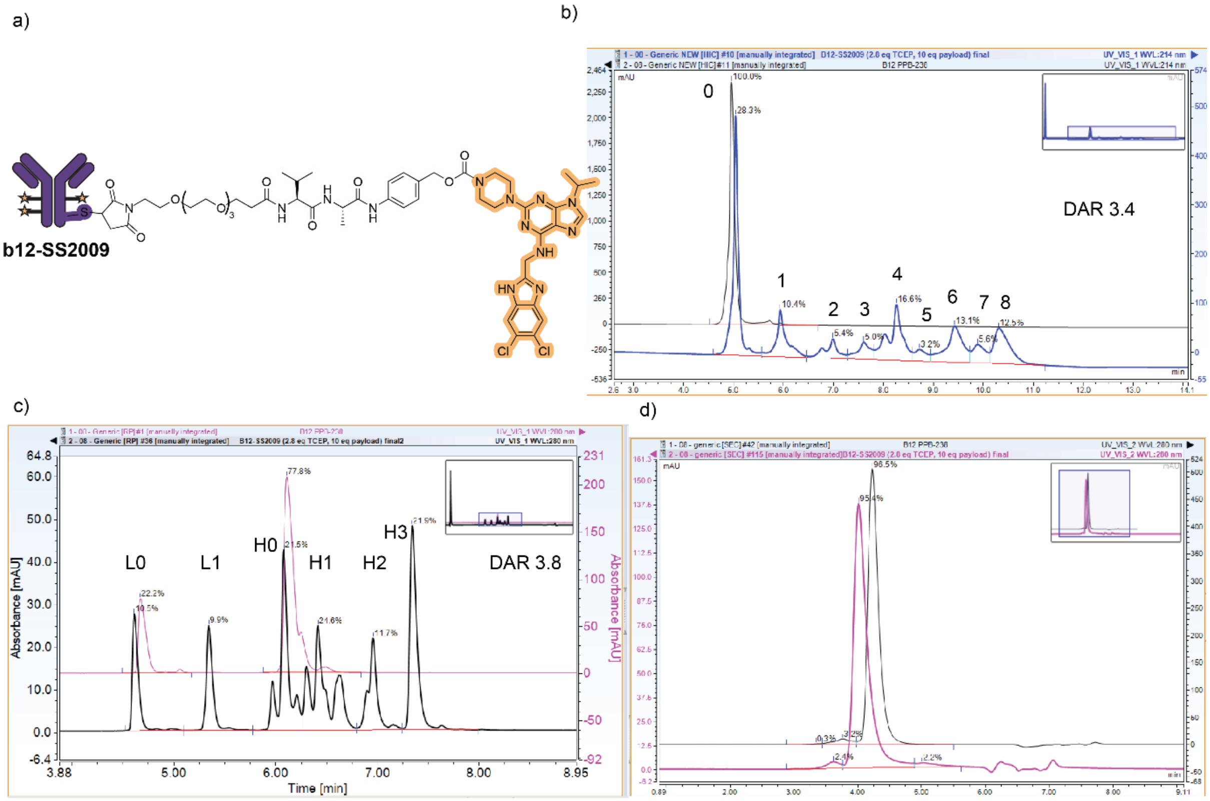


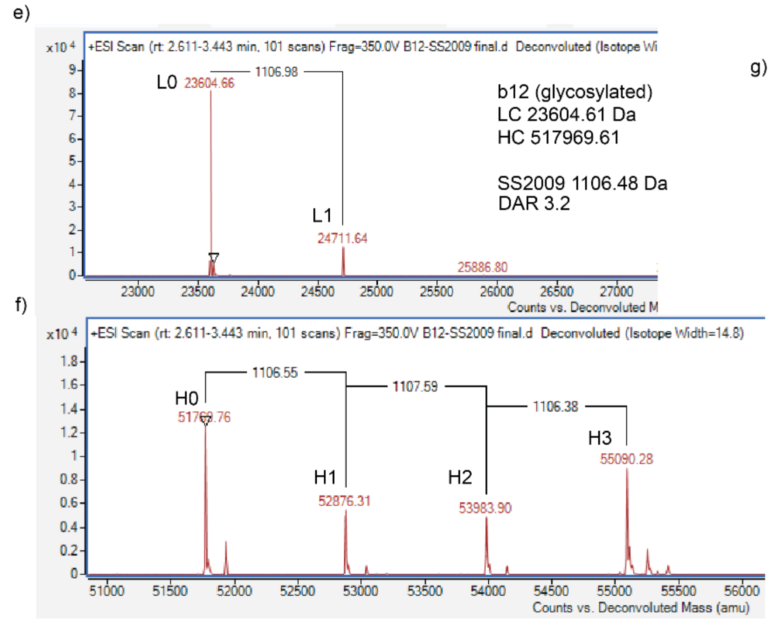


g)

| | B12-SS2009 | |
|---|---|---|
| TCEP | 2.8 | eq |
| Input | 23 | mg |
| [P] | 7.65 | mg/mL |
| Volume | 2.5 | mL |
| Amount | 19.1 | mg |
| Yield | 80.8 | % |
| DAR (PLRP) | 3.8 | |
| DAR (HIC) | 3.4 | |
| DAR (LC-MS) | 3.2 | |
| Monomer (SEC) | 95.4 | % |
| Endotoxin | 0.036 | EU/mg |
| Batch | PPB-1984 | |
| MW | 152596 | Da |
| $EC_{280}$ | 257368 | |

*Figure S11.* ***b12-SS2009 characterization. (a)*** *Representation of a ~DAR 4 b12-SS2009 DAC.* ***(b)*** *Hydrophobic interaction chromatography (HIC) spectrum for b12 (black) and b12-SS2009 (blue).* ***(c)*** *PLRP chromatogram for b12 (black) and b12-SS2009 (pink).* ***(d)*** *Size-exclusion chromatogram for b12 (black) and b12-SS2009 (pink).* ***(e)*** *LCMS spectrum for b12-SS2009, depicting light chain conjugation.* ***(f)*** *LCMS spectrum for b12-SS2009, depicting heavy chain conjugation.* ***(g)*** *Conjugation summary table for b12-SS2009.*

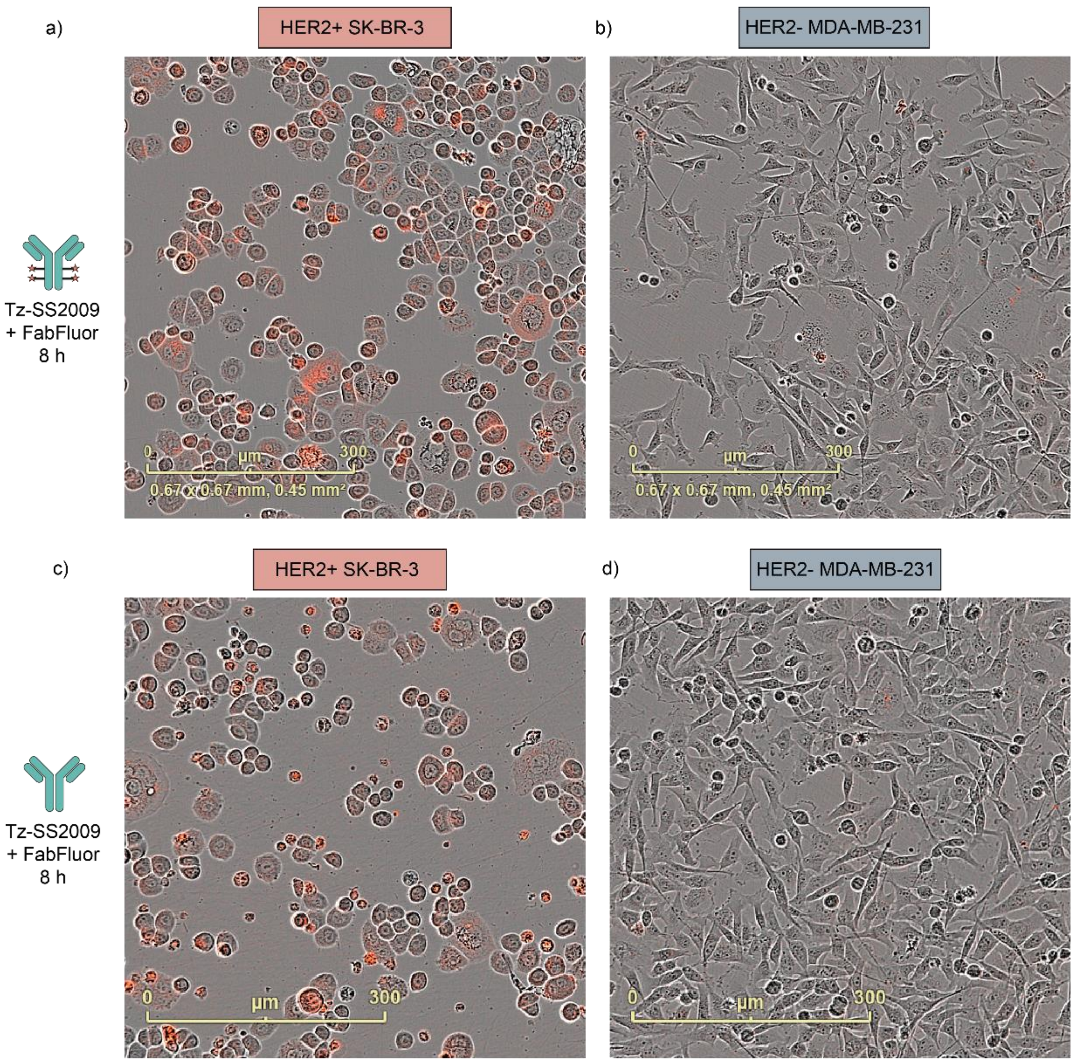


*Figure S12.* ***Representative images from the Tz-SS2009 internalization time-course assay at the 8 h time point.*** *HER2-positive SK-BR-3 cells (left) and HER2-negative MDA-MB-231 cells (right) were incubated with Tz-SS2009* ***(a), (b)*** *or unconjugated trastuzumab* ***(c), (d)*** *which had been incubated with FabFluor for 15 min prior to addition. A strong fluorescence signal was observed in SK-BR-3 cells, consistent with HER2-mediated antibody uptake and intracellular accumulation, whereas minimal fluorescence was detected in HER2-negative MDA-MB-231 cells. Similar levels of fluorescence were observed for Tz-SS2009 and unconjugated trastuzumab, indicating that payload conjugation does not substantially alter cellular internalization. Scale bar = 300 μm.*

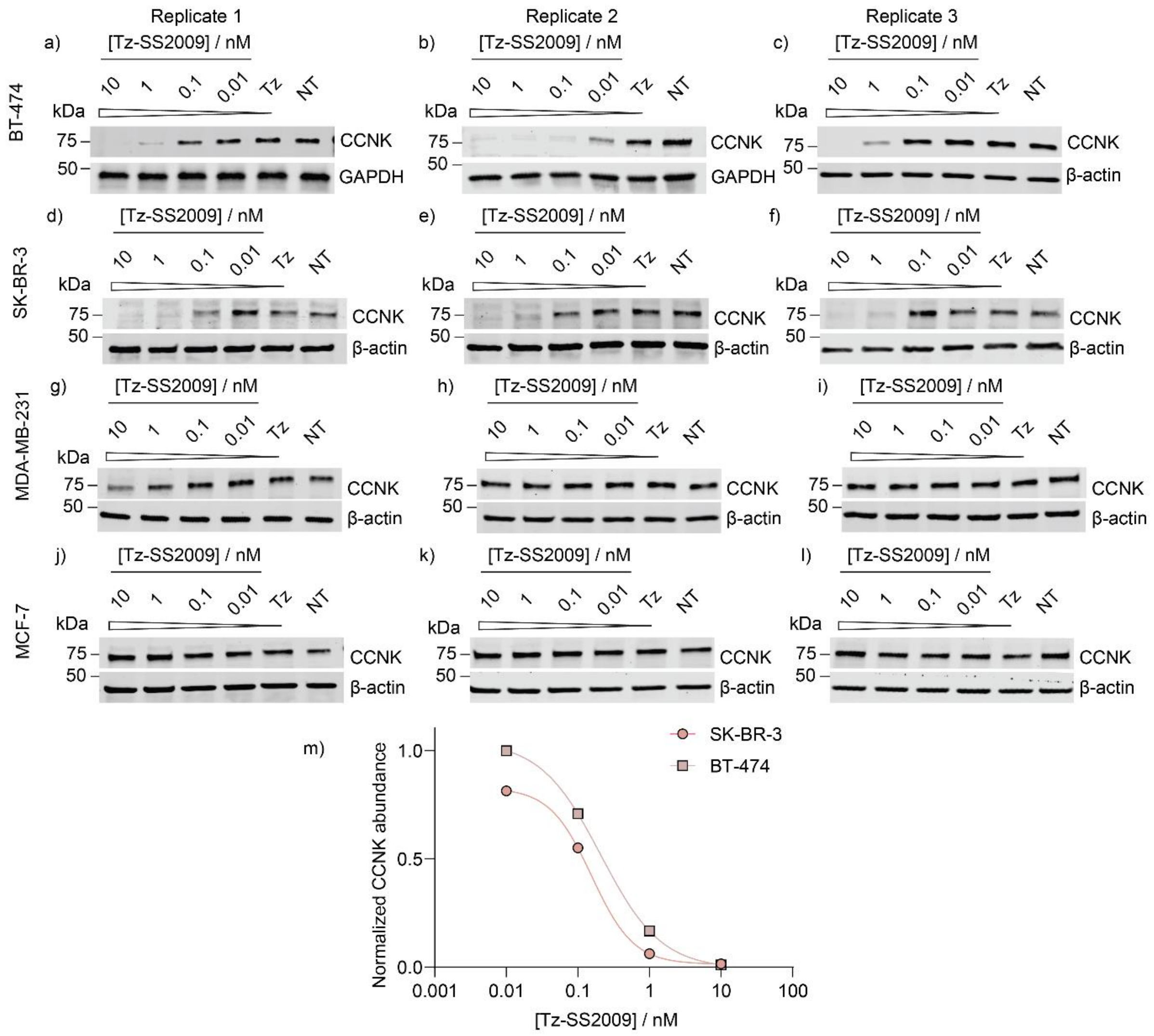


n)

| Cell line | $DC_{50}$ / pM [(a)] | $pDC_{50}$ ± SD [(b)] | Degradation at 10 nM / % [(c)] | n ($DC_{50}$ / $D_{10\,nM}$) |
|---|---|---|---|---|
| BT-474 | 270 | 9.57 ± 0.10 | 98 | 3 / 3 |
| SK-BR-3 | 210 | 9.7 ± 0.6 | 96 | 2 / 3 |

*Figure S13.* ***Western blots of treatment by Tz-SS2009.*** *Cells were seeded overnight, and treated with Tz-SS2009, unconjugated trastuzumab Tz (10 nM) or not treated for 72 h prior to lysis.* ***(a)-(c)****, BT-474 (HER2-positive);* ***(d)-(f)****, SK-BR-3 (HER2-positive);* ***(g)-(i)****, MDA-MB-231 (HER2-negative);* ***(j)-(l)****, MCF-7 (HER2-negative). Western blots shown are biological replicates, n=3.* ***(m)*** *Densitometric quantification of CCNK abundance in BT-474 and SK-BR-3 cells, normalized to loading control (β-actin or GAPDH, as indicated) and the untreated (NT) control. Data are from a single representative biological replicate with a four-parameter fit nonlinear regression curve.* ***(n)*** *Summary of CCNK degradation parameters. Apparent $DC_{50}$ and $pDC_{50}$ ± SD[b] were derived from four-parameter fits of each of the three independent biological replicates. [a] Geometric mean of n = 3 (back-transformed from the mean $pDC_{50}$). [b] Arithmetic mean ± SD of $-\log_{10} DC_{50}$ [M]; mean and SD calculated on log-transformed values. [c] Degradation at 10 nM = $D_{10\,nM}$ (1 − normalized CCNK abundance).*

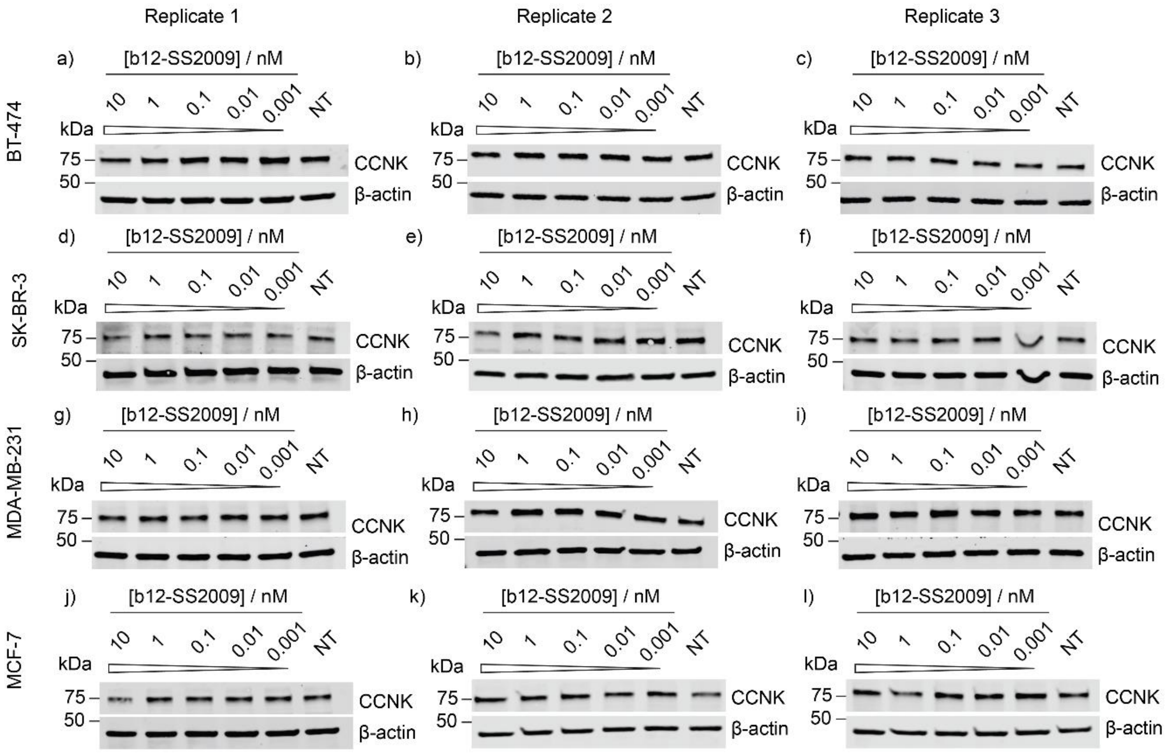


*Figure S14.* ***Western blots of treatment by b12-SS2009.*** *Cells were seeded overnight and treated with b12-SS2009 or not treated for 72 h prior to lysis.* ***(a)-(c)****, BT-474 (HER2-positive);* ***(d)-(f)****, SK-BR-3 (HER2-positive);* ***(g)-(i)****, MDA-MB-231 (HER2-negative);* ***(j)-(l)****, MCF-7 (HER2-negative). Western blots shown are biological replicates, n=3.*

a)

| DAC | [DAC] / nM | Rep 1 | Rep 2 | Rep 3 | Geometric mean | n |
|---|---|---|---|---|---|---|
| Tz-SS2009 | 10 | 0.014 | 0.011 | 0.025 | 0.016 | 3 |
| | 1 | 0.12 | 0.17 | 0.037 | 0.11 | 3 |
| | 0.1 | 0.81 | 0.71 | 0.071 | 0.34 | 3 |
| | 0.01 | 1.0 | 1.0 | 0.31 | 0.69 | 3 |
| b12-SS2009 | 10 | 0.83 | 0.72 | 0.61 | 0.71 | 3 |
| | 1 | 0.87 | 1.0 | 0.82 | 0.90 | 3 |
| | 0.1 | 0.74 | 1.2 | 1.2 | 1.0 | 3 |
| | 0.01 | 0.64 | 1.1 | 1.1 | 0.93 | 3 |

b)

| [DAC] / nM | Difference, $log_2$ | 95% CI ($log_2$) | Fold change | 95% CI (fold change) | Adjusted p-value |
|---|---|---|---|---|---|
| 10 | -5.5 | - 7.7 to -3.3 | 45 | 9.9 to 208 | <0.0001 |
| 1 | -3.1 | - 5.3 to -0.9 | 8.6 | 1.9 to 39 | 0.0046 |
| 0.1 | -1.5 | - 3.7 to 0.7 | 2.9 | 0.64 to 13 | 0.24 |
| 0.01 | - 0.4 | -2.6 to 1.8 | 1.4 | 0.30 to 6.2 | 0.97 |

c)

| Cell line | HER2 status | [Tz-SS2009] / nM | Rep 1 | Rep 2 | Rep 3 | Geometric mean | n |
|---|---|---|---|---|---|---|---|
| BT-474 | Positive | 10 | 0.014 | 0.011 | 0.025 | 0.016 | 3 |
| | | 1 | 0.19 | 0.17 | 0.037 | 0.11 | 3 |
| | | 0.1 | 0.81 | 0.71 | 0.071 | 0.34 | 3 |
| | | 0.01 | 1.0 | 1.0 | 0.31 | 0.69 | 3 |
| MDA-MB-231 | Negative | 10 | 0.59 | 0.88 | 0.99 | 0.80 | 3 |
| | | 1 | 0.86 | 0.88 | 0.91 | 0.88 | 3 |
| | | 0.1 | 1.2 | 1.2 | 0.93 | 1.1 | 3 |
| | | 0.01 | 1.2 | 1.1 | 0.89 | 1.1 | 3 |

d)

| [Tz-SS2009] / nM | Difference, $log_2$ | 95% CI ($log_2$) | Fold change | 95% CI (fold change) | Adjusted p-value |
|---|---|---|---|---|---|
| 10 | - 5.7 | - 7.8 to -3.5 | 51 | 11 to 230 | <0.0001 |
| 1 | - 3.1 | - 5.2 to -0.9 | 8.4 | 1.9 to 38 | 0.0044 |
| 0.1 | - 1.7 | - 3.8 to 0.5 | 3.2 | 0.71 to 14 | 0.17 |
| 0.01 | - 0.64 | - 2.8 to 1.5 | 1.6 | 0.14 to 7.0 | 0.89 |

*Figure S15.* ***Source data and full statistical comparisons for DAC-mediated CCNK degradation.*** *Densitometric quantification of CCNK abundance from the western blots in Figures 4a, 4c, S13 and S14, normalized to loading control (β-actin or GAPDH, as indicated on each blot) and to the untreated (NT) control.* ***(a)*** *Individual biological replicate values and geometric means for Tz-SS2009 and b12-SS2009 in HER2-positive BT-474 cells after 72 h treatment.* ***(b)*** *Pairwise comparisons of Tz-SS2009 against b12-SS2009 in BT-474 at each matched concentration.* ***(c)*** *Individual replicate values and geometric means for Tz-SS2009 in HER2-positive BT-474 and HER2-negative MDA-MB-231 cells; the BT-474 values are the same data shown in* ***(a)****.* ***(d)*** *Pairwise comparisons of Tz-SS2009 between BT-474 and MDA-MB-231 at each matched concentration. In* ***(b)*** *and* ***(d)****, normalized abundances were $log_2$-transformed and analysed by ordinary two-way ANOVA with Šídák's correction for multiple comparisons. Differences are reported on the $log_2$ scale with 95% confidence intervals, and back-*

*transformed to fold changes (2^Δ). Negative differences indicate lower CCNK abundance in the first-named condition. The comparisons at 10 and 1 nM in* ***(b)*** *and* ***(d)*** *are those quoted in the main text and in Figure 4b and 4d. All concentrations refer to antibody (conjugate) equivalents.*

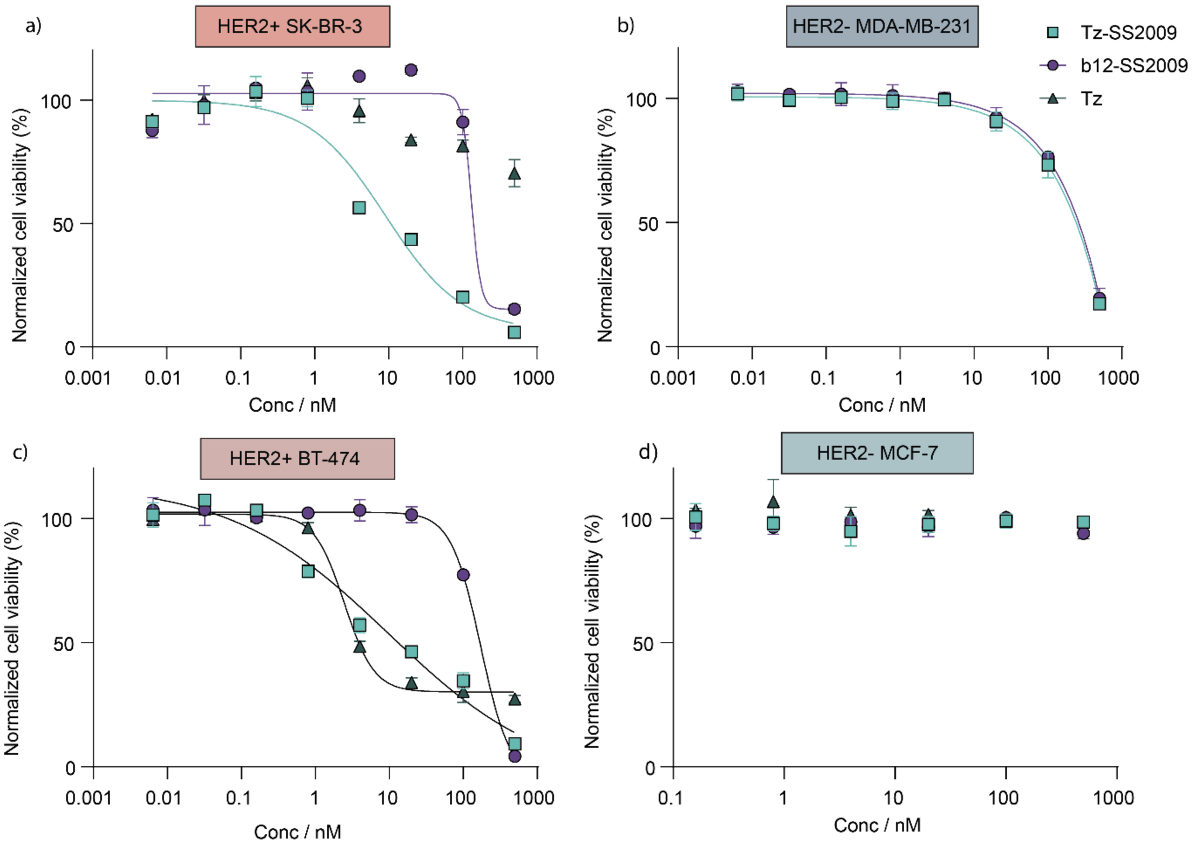


e)

| Cell line | Treatment | $IC_{50}$ / nM [(a)] | $pIC_{50}$ ±SD [(b)] | Viability at 500 nM / % [(c)] | n |
|---|---|---|---|---|---|
| BT-474 | Tz -SS2009 | 5.0 | 8.3 ± 0. 3 | 6 ± 3 | 3 |
| | b12 -SS2009 | 30 | 6.89 ± 0.12 | 2.9 ± 1.3 | 3 |
| | Tz | 5.9 | 8.2 ± 0.2 | 19 ± 7 | 3 |
| SK-BR-3 | Tz -SS2009 | 14 | 7. 9 ± 0. 5 | 7 ± 4 | 3 |
| | b12 -SS2009 | 200 | 6.70 ± 0.12 | 9 ± 7 | 2 [(d)] |
| | Tz | > 500 | n.d. | 61 ± 9 | 3 |
| MCF-7 | Tz -SS2009 | > 500 | n.d. | 106 ± 22 | 3 |
| | b12 -SS2009 | > 500 | n.d. | 104 ± 19 | 3 |
| | Tz | > 500 | n.d. | 100 ± 5 | 3 |
| MDA-MB-231 | Tz -SS2009 | 180 | 6.75 ± 0.06 | 17 ± 7 | 3 |
| | b12 -SS2009 | 200 | 6.69 ± 0.08 | 23 ± 5 | 3 |
| | Tz | n.t. [(g)] | n.d. [(f)] | n.d. [(f)] | n.a. [(g)] |

*Figure S16.* ***Effects of DAC treatment on cell proliferation.*** *Representative dose–response curves for the HER2-targeted conjugate (Tz-SS2009), the non-targeting isotype-control conjugate (b12-SS2009) and naked Tz in* ***(a)*** *HER2-positive SK-BR-3,* ***(b)*** *HER2-negative MDA-MB-231,* ***(c)*** *HER2-positive BT-474 and* ***(d)*** *HER2-negative MCF-7 cells. Cell viability was determined by MTS assay after 9 days treatment (7 days for MCF-7) and normalized to vehicle-treated and puromycin controls. 3 independent biological experiments were performed in technical triplicate (technical duplicate for BT-474), with* ***(a)-(d)*** *depicting a representative dose-response curve per cell line. Points are mean ± SD of the* *technical replicates, curves are four-parameter nonlinear regression. Concentrations refer to antibody (conjugate) equivalents.* ***(e)*** *$IC_{50}$ (geometric mean), $pIC_{50}$ ± SD and normalized viability at the highest concentration tested (500 nM).* [a] *$IC_{50}$ values are geometric means of 3 independent experiments.* [b] *$pIC_{50}$ ($-\log_{10}$ $IC_{50}$ [M]), mean ± SD; means and SD calculated on log-transformed values.* [c] *Mean ± SD normalized viability at 500 nM.* [d] *Three biological replicates were performed; curve fitting was possible for two, so $IC_{50}$ and $pIC_{50}$ are reported for n = 2, whereas viability at 500 nM is reported for n = 3.* [e] *50% inhibition was not reached within the range tested; $IC_{50}$ is reported as greater than the highest concentration (500 nM).* [f] *n.d., not determined.* [g] *n.t., not tested; n.a., not applicable.*

a)

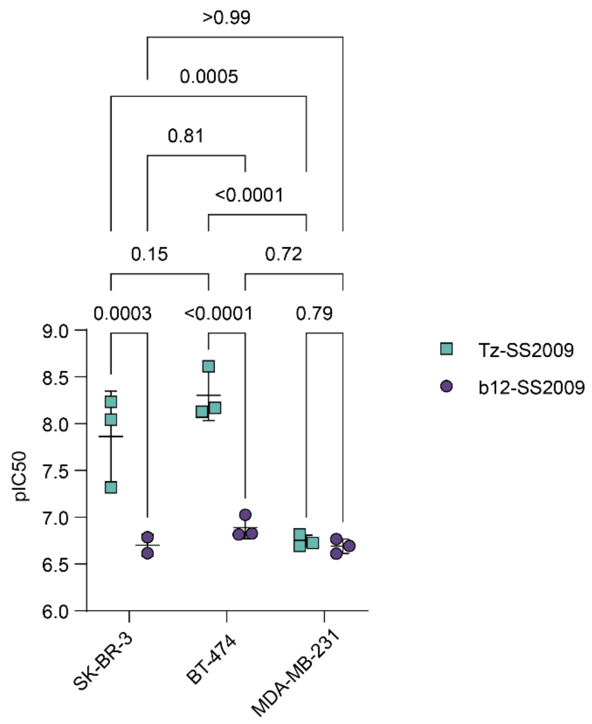


b)

| Comparison | $\Delta pIC_{50}$ (95% CI) | Fold difference (95% CI) | Adjusted p | Comparisons in family |
|---|---|---|---|---|
| Within cell line (Tz-SS2009 vs b12-SS2009) | | | | |
| SK-BR-3 | 1.16 (0.67 to 1.66) | 15 (4.6 to 46) | 0.0003 | 1 |
| BT-474 | 1.41 (0.97 to 1.86) | 26 (9.3 to 72) | <0.0001 | 1 |
| MDA-MB-231 | 0.06 (−0.39 to 0.50) | 1.1 (0.4 to 3.2) | 0.79 | 1 |
| Between cell lines, Tz-SS2009 | | | | |
| SK-BR-3 vs BT-474 | −0.44 (−1.01 to 0.13) | 0.36 (0.10 to 1.3) | 0.15 | 3 |
| SK-BR-3 vs MDA-MB-231 | 1.12 (0.55 to 1.69) | 13 (3.6 to 49) | 0.0005 | 3 |
| BT-474 vs MDA-MB-231 | 1.56 (0.989 to 2.13) | 36 (9.8 to 134) | <0.0001 | 3 |
| Between cell lines, b12-SS2009 | | | | |
| SK-BR-3 vs BT-474 | −0.19 (−0.82 to 0.45) | 0.65 (0.15 to 2.8) | 0.1 | 3 |
| SK-BR-3 vs MDA-MB-231 | 0.01 (−0.63 to 0.65) | 1.0 (0.24 to 4.4) | >0.99 | 3 |
| BT-474 vs MDA-MB-231 | 0.20 (−0.37 to 0.78) | 1.6 (0.43 to 5.9) | 0.72 | 3 |

*Figure S17* ***Full statistical comparison of $pIC_{50}$ values for Tz-SS2009 and b12-SS2009 across HER2-positive and HER2-negative cell lines. (a)*** *$pIC_{50}$ values derived from four-parameter fits of individual dose–response curves in HER2-positive SK-BR-3 and BT-474 and HER2-negative MDA-MB-231 cells following 9 days treatment. Points are individual biological replicates ($n = 3$, except b12-SS2009 in SK-BR-3 where curve fitting was possible for only two replicates, $n = 2$); bars show mean ± SD. Numbers above brackets are adjusted p values from Šídák's multiple-comparisons test. Data are as shown in Figure 4g, with all nine pairwise comparisons displayed.* ***(b)*** *Complete multiple-comparisons output. Data were analyzed by ordinary two-way ANOVA (factors: cell line and treatment) with Type III sums of squares; the interaction was significant ($F(2, 11) = 12.58$, $p = 0.0014$). Comparisons were grouped into five families with Šídák's correction applied within each family. Fold differences are back-transformed from $\Delta pIC_{50}$ ($10^{\Delta}$); values below 1 indicate the first-named condition is less potent. Tz-*

*SS2009 was significantly more potent than b12-SS2009 in both HER2-positive lines but not in MDA-MB-231, and b12-SS2009 potency did not differ between any pair of cell lines. Concentrations refer to antibody (conjugate) equivalents.*

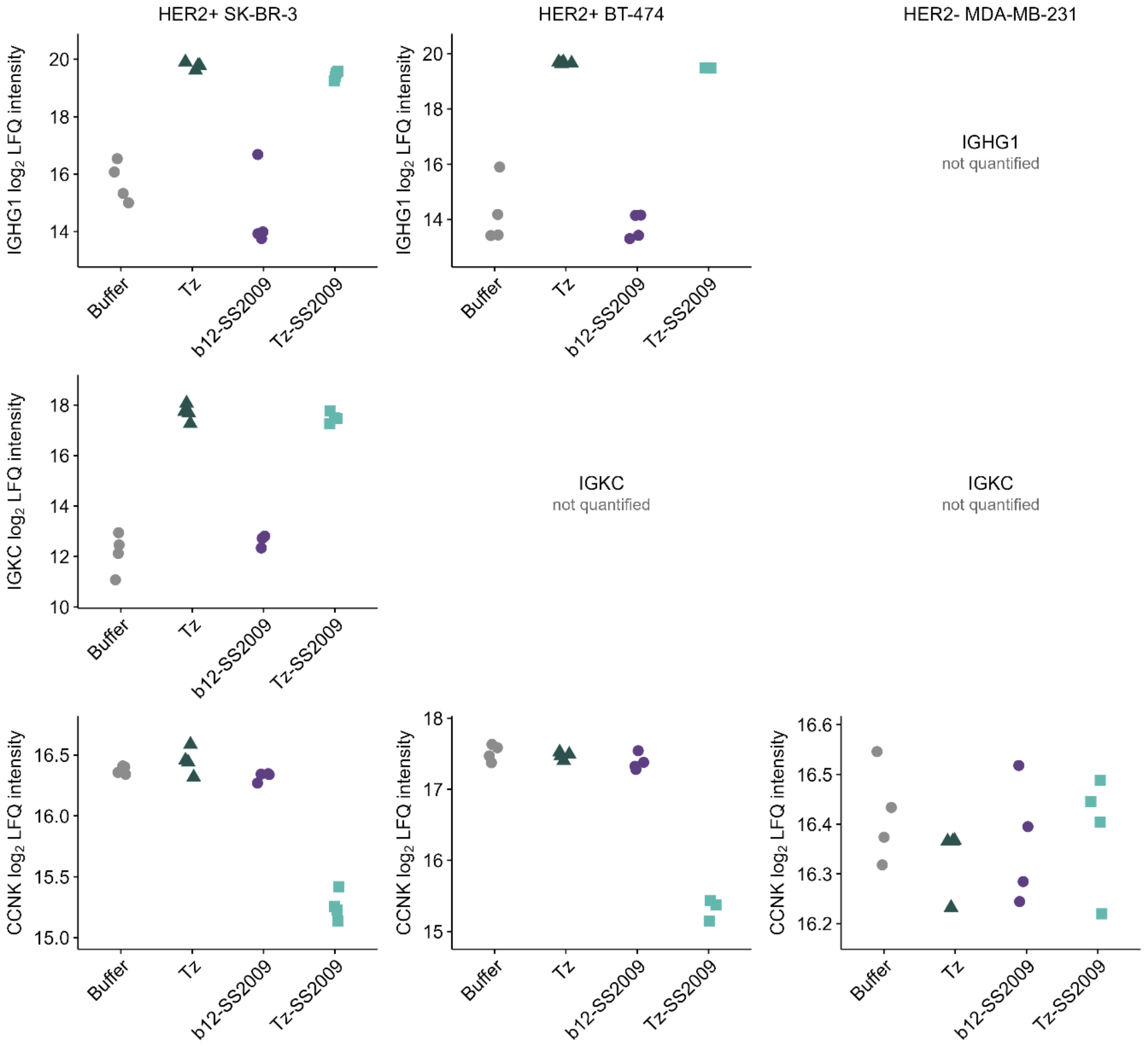


*Figure S18.* ***Antibody uptake and CCNK depletion across all three cell lines.*** *$log_2$ LFQ intensities of IGHG1, IGKC and CCNK (top-to-bottom rows) in HER2-positive SK-BR-3 and BT-474, and MDA-MB-231 (HER2-negative) after 24 h of treatment with 10 nM Tz-SS2009, trastuzumab (Tz), b12-SS2009, or formulation buffer. Each point is an individual replicate ($n$ = 4 per condition, except Tz-SS2009 in BT-474, $n$ = 3) and horizontal bars denote group means. Shape and colour both encode treatment: square, Tz-SS2009; triangle, Tz; circles, buffer and b12-SS2009. Axes are scaled independently in each panel. The immunoglobulin derived proteins IGKC and IGHG1 were not found in all datasets; IGKC was only found in SK-BR-3 while IGHG1 was present in BT-474 and SK-BR-3 but not in MDA-MB-231.*

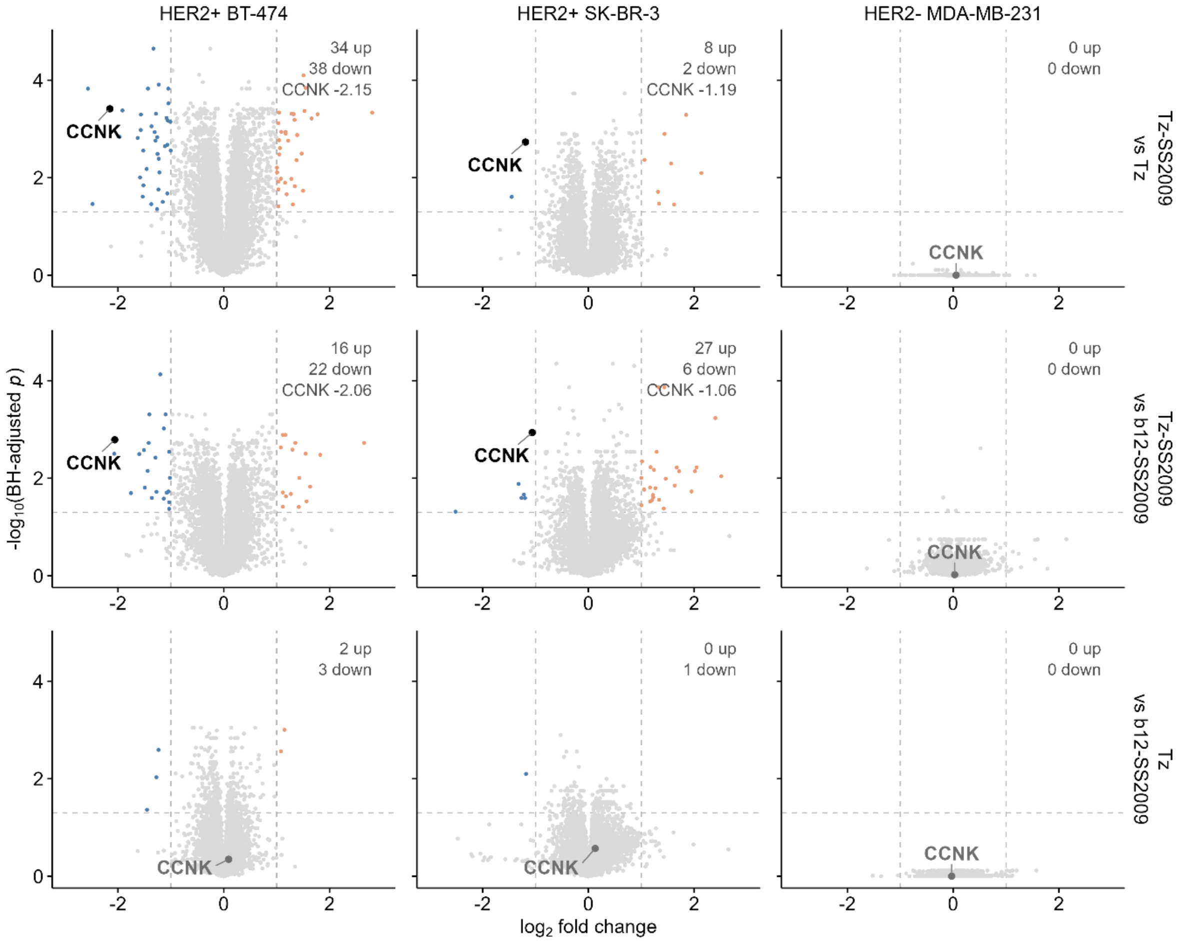


*Figure S19.* ***Direct contrasts between treatments across all three cell lines.*** *Volcano plots of all pairwise treatment contrasts from the 24 h DIA proteomics experiment, isolating the contribution of the delivered degrader from that of antibody binding alone. Rows: Tz-SS2009 versus Tz (top), Tz-SS2009 versus b12-SS2009 (middle), and Tz versus b12-SS2009 (bottom). Columns: HER2-positive BT-474 and SK-BR-3, and HER2-negative MDA-MB-231. All treatments were at 10 nM for 24 h (n = 4, except Tz-SS2009 in BT-474, n = 3). Significance was assessed by two-sided Student's t-test with Benjamini–Hochberg correction applied within each contrast, with immunoglobulin proteins excluded before correction. Proteins passing adj. $p < 0.05$ and $|log_2FC| > 1$ are coloured by direction (orange, increased; blue, decreased), with dashed lines marking the thresholds and the number passing in each direction given at the top right of each panel. CCNK is labelled throughout, in black where it passes both thresholds and grey where it does not. CCNK $log_2$ fold change given where significant. Axes are identical across all nine panels.*

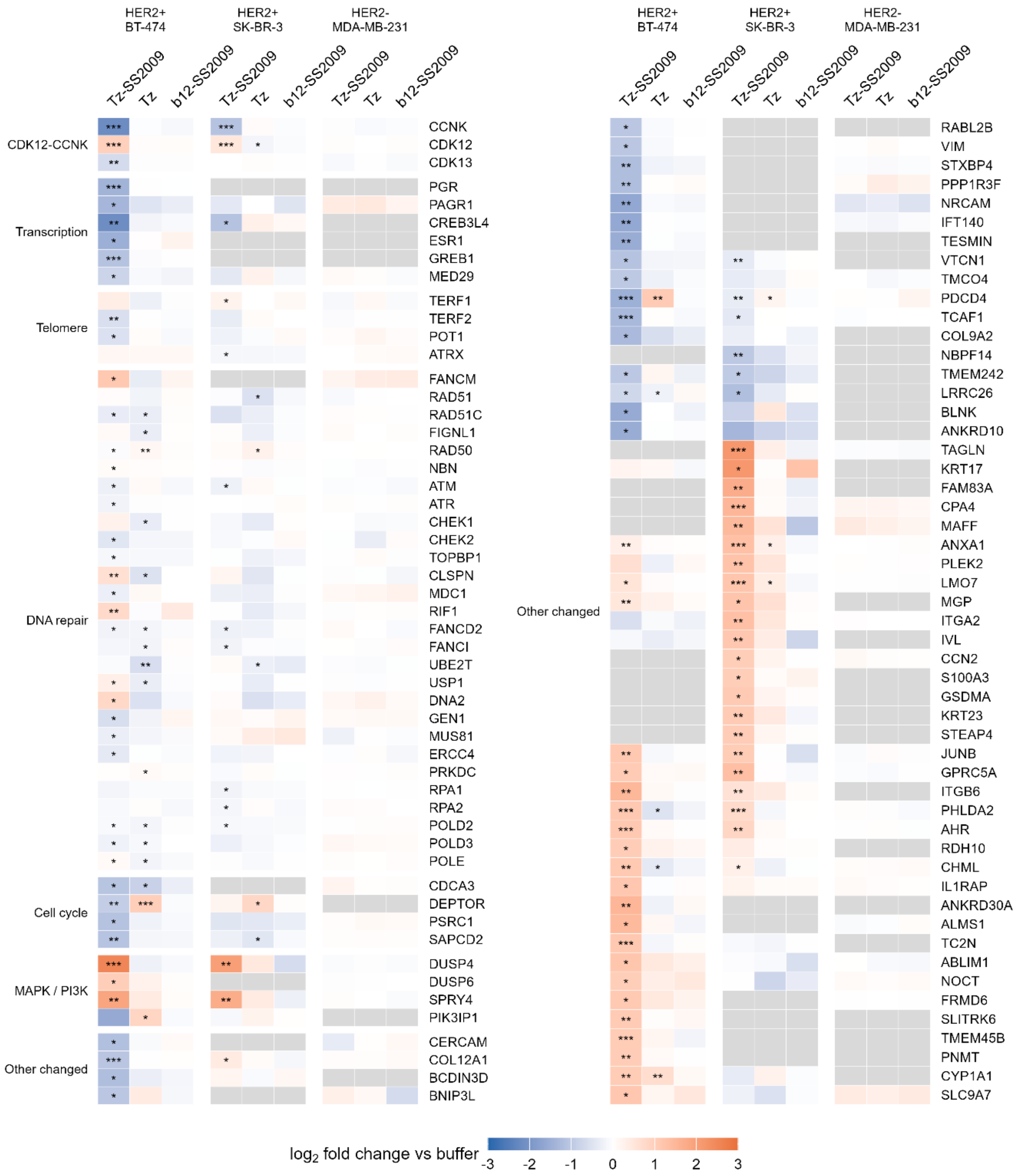


*Figure S20.* ***Full proteomic response to Tz-SS2009 across HER2-positive and HER2-negative cells.*** *Expanded version of Figure 5d, showing all curated candidate proteins and all further proteins reaching significance, across three cell lines and three treatments. Samples, contrasts and statistics are as in Figure 5 (24 h, 10 nM, each treatment versus formulation buffer; n = 4, except Tz-SS2009 in BT-474, n = 3). Tiles show $\log_2$ fold change versus buffer, truncated at ±3. Asterisks denote adjusted p values (* adj. $p < 0.05$, ** adj. $p < 0.01$, *** adj. $p < 0.001$) and are independent of the fold-change criterion used for the counts in Figure 5a. Grey indicates that a protein was not quantified in that cell line. Rows come from two sources: curated candidates, retained where they reach adj. $p < 0.05$ in at least one displayed contrast, with the CDK12–CCNK block shown irrespective of significance; and further proteins recruited on adj. $p < 0.05$ with $|\log_2 FC| > 1$ for Tz-SS2009 versus buffer in any line ("Other changed"), and ordered by hierarchical clustering of their $\log_2$ fold changes across lines.*

#### Synthesis

Chemicals and solvents were purchased from Sigma-Aldrich UK, Fluorochem UK, Tocris, Apollo Scientific UK and Acros Organics, and used without further purification.

Reactions were monitored by TLC, NMR or LC-MS. Analytical TLC was performed on aluminum support silica gel plates (60 F254, Merck) and visualized by UV light (254 nm), followed by staining with aqueous potassium permanganate solution or ninhydrin and heating with a heat gun.

LC-MS Method A was performed on an Agilent 1260 II machine with an Agilent Poroshell HPH C18 3.0 x 50 mm 2.7 µm basic column (95% $H_2O$ (0.1% $NH_3$) to 95% MeCN (0.1% $NH_3$) in 8 min, then 95% $H_2O$ (0.1% $NH_3$) for 2 min). LC-MS Method B was performed on an Agilent Infinity II 1260 machine with an InfinityLab Poroshell 120 EC-C18 2.1 x 50 mm 1.9 µm column. A flow rate of 0.5 mL/min with an injection volume of 2.5 µL was used. LC traces were recorded on an LC single quad InfinityLab LC/MSD MS in positive or negative mode. The following parameters were used: 12.0 L/min $N_2$ at 350 °C, 35 psig nebulizer pressure, 3000 V capillary volume. Mobile phase A: $H_2O$ (0.1% $CH_2O_2$) mobile phase B: MeCN; 95% A for 3.5 min then 95% A to 95% B over 1.5 min, hold at 95% B for 0.5 min 95% B to 95% A over 0.1 min.

NMR experiments were performed on Bruker AV instruments at RT and referenced to the respective deuterated solvent peaks. NMR analysis was performed by MestReNova 14.3.3 (2023). Data are given as chemical shift, multiplicity, coupling constants (J), integration and assignment. Multiplicity is reported as follows: s = singlet, d = doublet, t = triplet, q = quartet, sept = septet, m = multiplet, br = broad.

Column purifications were carried out by either manual flash column chromatography using Geduran silica 60 (0.040-0.063 mm) silica gel or by automated flash column chromatography using a Biotage Selekt Flash Chromatography Instruments with the stated solvent systems. The final product was purified by high-performance liquid chromatography (HPLC) on a Shimadzu preparative HPLC with an Aeris Peptide 5 µm XB-C18 column.

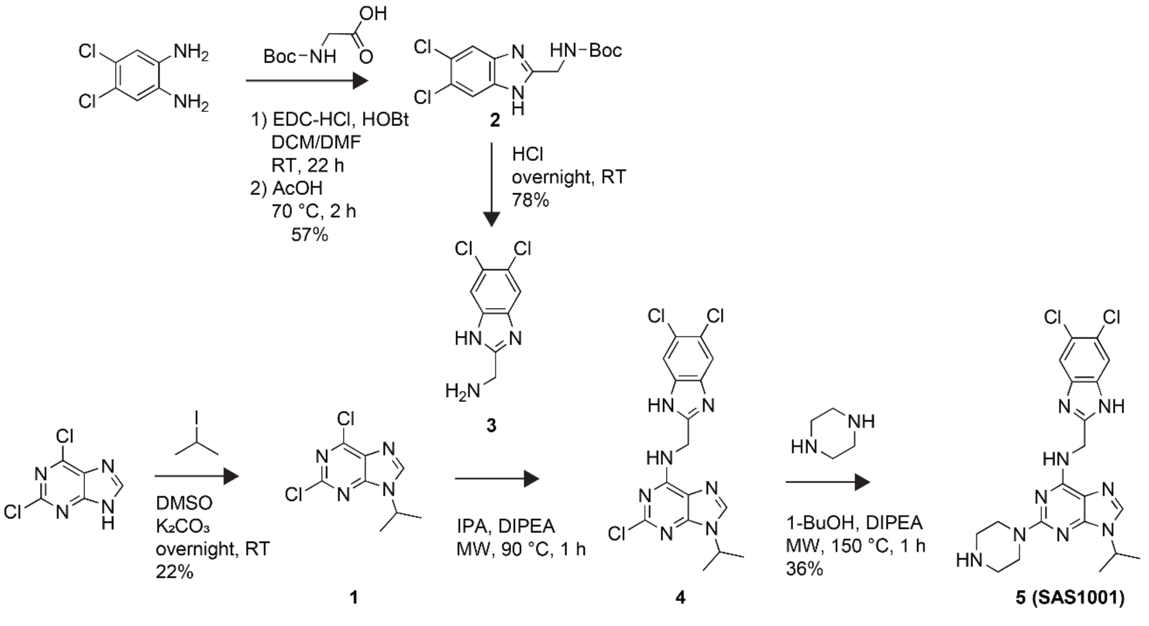


*Scheme S1* ***Synthetic scheme for SAS1001 (degrader).***

**2,6-dichloro-9-isopropyl-9*H*-purine (1)**

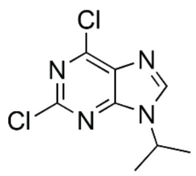

The following protocol was adapted from Kozicka *et al.* and Hu *et al.*[1,2]

2,6-Dichloropurine (249 mg, 1.30 mmol) and potassium carbonate (557 mg, 4.00 mmol) were dissolved in anhydrous DMSO (5 mL). 2-Iodopropane (0.67 mL, 6.6 mmol) was added dropwise, and the reaction mixture stirred overnight. The product was precipitated into ice-cold water (10 mL) and purified by automated flash column chromatography (0-50% EtOAc in hexane) to yield **1** (66.1 mg, 22%) as a white solid. This reaction was scaled for material.

$^1$H NMR ($CDCl_3$, 400 MHz) δ [ppm] 8.14 (s, 1H), 4.89 (sept, J = 6.8 Hz, 1H), 1.63 (d, J = 6.8 Hz, 6H).

LC-MS (5-95% MeCN in $H_2O$, positive mode) Rt = 4.23 min, $[M+H]^+$ = 231.0

**tert-butyl ((5,6-dichloro-1*H*-benzo[*d*]imidazol-2-yl)methyl)carbamate (2)**

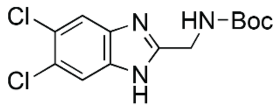

The following protocol was adapted from WO2019217421A1.[3]

Boc-Gly-OH (2.09 g, 11.9 mmol), 1-ethyl-3-(3-dimethylaminopropyl)carbodiimide hydrochloride (2.84 g, 14.9 mmol) and 1-hydroxybenzotriazole monohydrate (2.50 g, 16.3 mmol) were dissolved in anhydrous DCM (115 mL). The mixture was stirred at RT for 15 min until dissolved. 4,5-Dichloro-1,2-benzenediamine (2.10 g, 11.9 mmol) and anhydrous DMF (1 mL) were added. The reaction mixture was stirred for 22 h at RT. The volatiles were removed, and ethyl acetate added (200 mL). The organic phase was washed with brine (3x), saturated ammonium chloride solution (2x), saturated sodium bicarbonate solution (2x) and brine (1x) before drying over anhydrous sodium sulfate. Following concentration, glacial acetic acid (13 mL) was added, and the mixture was heated at 70 °C for 2 h. Precipitation into DCM:$Et_2O$:MeOH (90:8:2) was performed to yield **2** (2.15 g, 57%) as an off-white solid.

$^1$H NMR (DMSO-$d_6$, 400 MHz) δ [ppm] 12.51 (s, 1H), 7.76 (s, 2H), 7.49 (t, J = 5.9 Hz, 1H), 4.35 (d, J = 5.9 Hz, 2H), 1.41 (s, 9H).

LC-MS (5-95% MeCN in $H_2O$, positive mode) Rt = 4.54 min, $[M+H]^+$ = 316.1

**(5,6-dichloro-1*H*-benzo[*d*]imidazol-2-yl)methanamine (3)**

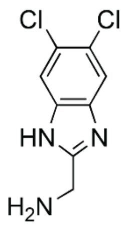

To **2** (754 mg, 2.39 mmol) dissolved in ethyl acetate (50 mL) was added HCl (3 mL, 12 M, 36 mmol). After 30 min, ethyl acetate (10 mL) was added, and the mixture was stirred overnight. The white precipitate was filtered and washed in ice-cold $Et_2O$ to yield **3** as a white solid (533 mg, 78%).

$^1$H NMR (400 MHz, DMSO-$d_6$) δ 8.86 (s, 3H), 7.94 (s, 2H), 6.25 (br, s, 4H), 4.35 (q, $J$ = 5.4 Hz, 2H).

LC-MS (5-95% MeCN in $H_2O$, positive mode) Rt = 2.99 min, $[M+H]^+$ = 216.0

**2-chloro-*N*-((5,6-dichloro-1*H*-benzo[*d*]imidazol-2-yl)methyl)-9-isopropyl-9*H*-purin-6-amine (4)**

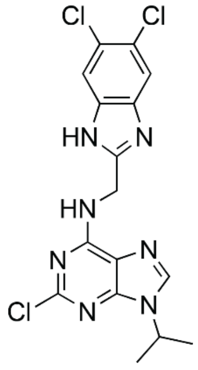

The following protocol was adapted from Monastyrskyi et al.[4]

**1** (528 mg, 2.28 mmol), **3** (635 mg, 2.52 mmol), anhydrous DIPEA (2.0 mL, 11.4 mmol) and isopropanol (9.15 mL) were combined and heated at 90 °C for 1 h in a microwave reactor. Upon completion of the reaction, as assessed by LC-MS, the reaction mixture was concentrated *in vacuo* and used in the next step without further purification.

LC-MS (5-95% MeCN in $H_2O$, positive mode) Rt = 4.29 min, $[M+H]^+$ = 410.1

***N*-((5,6-dichloro-1*H*-benzo[*d*]imidazol-2-yl)methyl)-9-isopropyl-2-(piperazin-1-yl)-9*H*-purin-6-amine (5, SAS1001)**

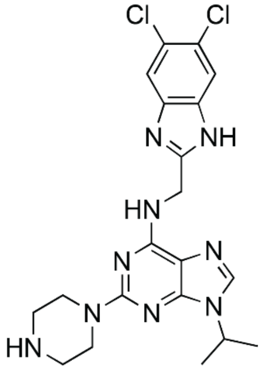

To the crude reaction mixture from the previous step (**4**) was added piperazine (1.13 g, 13.1 mmol), anhydrous DIPEA (0.68 mL, 3.9 mmol) and 1-butanol (2.0 mL). The reaction mixture was heated to 150 °C for 1 h in a microwave reactor. The reaction mixture was concentrated *in vacuo* and extracted into toluene/methanol (9:1) prior to purification in four portions by automated reversed phase flash column chromatography (5-95% MeCN (0.1% FA) in $H_2O$). The combined fractions yielded 5 (**SAS1001)** a white solid as the mono-formate salt (418 mg, 36%) across two steps from **1**.

$^1H$ NMR (DMSO-d$_6$, 400 MHz) δ [ppm] 8.11 (s, 1H), 7.94 (s, 1H), 7.72 (s, 2H), 4.78 (s, 2H), 4.59 (sept, $J$ = 6.8 Hz, 1H), 3.60 (s, 4H), 2.77–2.68 (m, 4H), 1.48 (d, $J$ = 6.7 Hz, 6H).

$^{13}C$ NMR (101 MHz, MeOD) δ 159.23, 158.37, 155.46, 137.96, 126.95, 44.79, 43.61, 40.32, 22.48.

LC-MS (5-95% MeCN in $H_2O$, positive mode) Rt = 3.786 min $[M+H]^+$ = 460.2

HRMS (TOF MS ES$^+$) Calc. for $C_{20}H_{24}N_9Cl_2^+$ = 460.1532, found $[M+H]^+$ = 460.1541

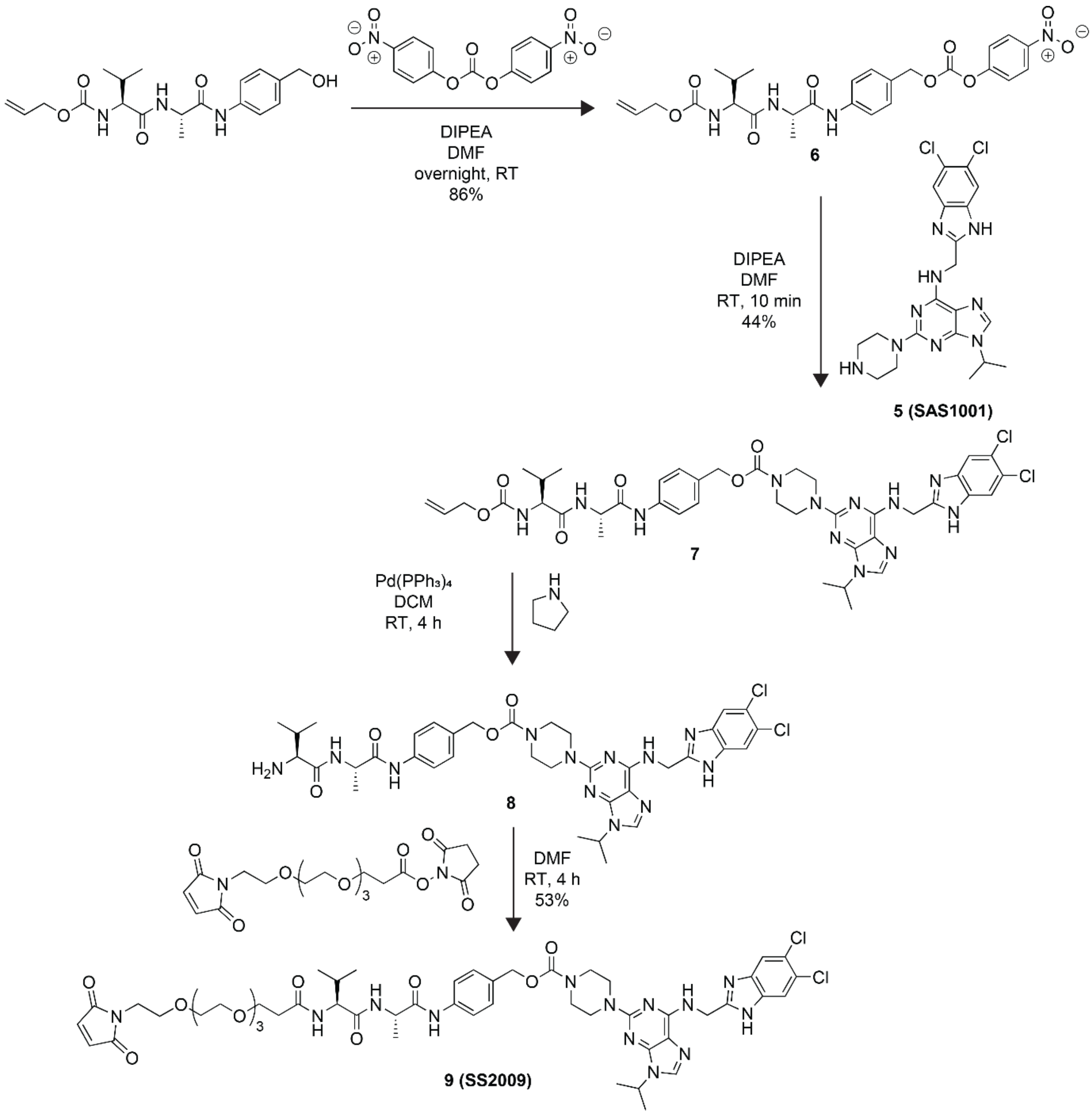


*Scheme S2* ***Synthetic scheme for SS2009 (linker-degrader).***

**Allyl-((*S*)-3-methyl-1-(((*S*)-1-((4-((((4-nitrophenoxy)carbonyl)oxy)methyl)phenyl)amino)-1-oxopropan-2-yl)amino)-1-oxobutan-2-yl)carbamate (6)**

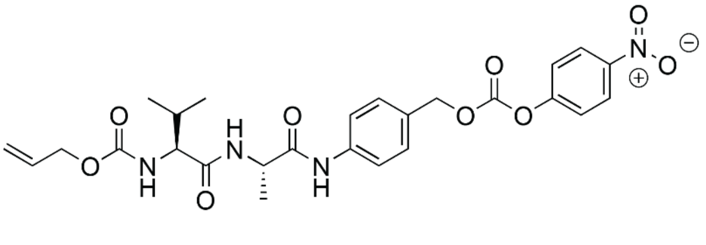

The following protocol was adapted from Cazzamalli et al.[5]

Alloc-Val-Ala-PAB-OH (249 mg, 0.66 mmol) and bis(4-nitrophenyl)carbonate (414 mg, 1.37 mmol) were dissolved in anhydrous DMF (1 mL) under a nitrogen atmosphere. DIPEA (1.50 mL, 8.6 mmol) was added, and the reaction mixture was stirred overnight at RT. Following concentration under a flow of $N_2$, precipitation in ice-cold $Et_2O$ (3 x 15 mL) yielded **6** (310 mg, 86%) as an off-white solid.

LC-MS (5-95% MeCN in $H_2O$, positive mode) Rt = 4.71 min, $[M+Na]^+$ = 565.3

$^1$H NMR (DMSO-$d_6$, 400 MHz) δ [ppm] 10.07 (s, 1H), 8.30 (s, 1H), 8.19 (d, *J* = 6.9 Hz, 1H), 7.63 (d, *J* = 8.3 Hz, 2H), 7.57 (d, *J* = 9.2 Hz, 2H), 7.42 (d, *J* = 8.3 Hz, 2H), 7.26 (d, *J* = 8.8 Hz, 1H), 5.91 (ddt, *J* = 16.1, 10.5, 5.3 Hz, 1H), 5.30 (d, *J* = 17.3 Hz, 1H), 5.24 (s, 2H), 5.17 (d, *J* = 10.5 Hz, 1H), 4.48 (d, 2H, *J* = 4.11 Hz), 4.41 (t, *J* = 7.0 Hz, 1H), 3.89 (t, *J* = 7.9 Hz, 1H), 1.97 (q, *J* = 6.9 Hz, 1H), 1.31 (d, *J* = 7.1 Hz, 3H), 0.89 (d, *J* = 6.8 Hz, 3H), 0.84 (d, *J* = 6.8 Hz, 3H).

**4-((*S*)-2-((*S*)-2-(((allyloxy)carbonyl)amino)-3-methylbutanamido)propanamido)benzyl 4-(6-(((5,6-dichloro-1*H*-benzo[d]imidazol-2-yl)methyl)amino)-9-isopropyl-9*H*-purin-2-yl)piperazine-1-carboxylate (7)**

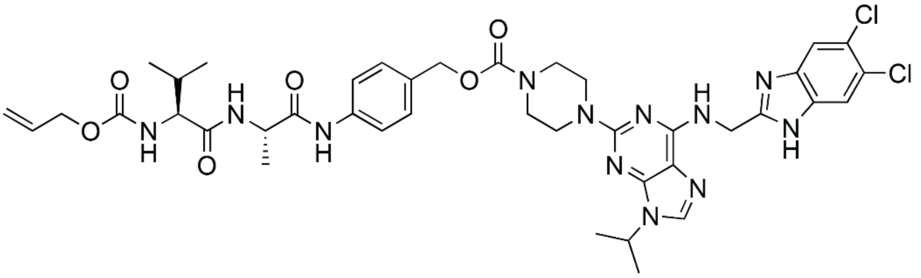

**SAS1001 mono-formate** (69.8 mg, 0.14 mmol) was suspended in anhydrous DMF (1 mL) under a nitrogen atmosphere at RT, and DIPEA (100 µL, 0.56 mmol) was added. **6** (56.4 mg, 0.10 mmol) was dissolved in anhydrous DMF (0.2 mL) and added dropwise. The reaction mixture was sonicated at 30 °C for 1 min until full dissolution, at which point LC-MS confirmed reaction completion. MeOH (1 mL) was added, and the reaction mixture was poured into toluene (150 mL) before concentration *in vacuo*. The residue was purified by automated reversed phase flash column chromatography (5-95% MeCN (0.1% FA) in $H_2O$), yielding **7** (39.3 mg, 44%) as a white solid. This reaction was subsequently repeated to provide material for the following step.

LC-MS (5-95% MeCN in $H_2O$, positive mode) Rt = 5.84 min, $[M+H]^+$ = 863.3

$^1$H NMR (400 MHz, DMSO-$d_6$) δ 12.42 (s, 1H), 10.02 (s, 1H), 8.19 (d, *J* = 6.9 Hz, 1H), 8.07 (s, 1H), 7.93 (s, 1H), 7.79 (s, 1H), 7.67 – 7.50 (m, 3H), 7.28 (dd, *J* = 16.8, 8.5 Hz, 3H), 5.91 (td, *J* = 10.9, 5.2 Hz, 1H), 5.30 (d, *J* = 17.2 Hz, 1H), 5.17 (d, *J* = 10.6 Hz, 1H), 5.01 (s, 2H), 4.77 (s, 2H), 4.69 – 4.53 (m, 1H), 4.44 (dd, *J* = 26.0, 6.1 Hz, 3H), 3.89 (t, *J* = 7.7 Hz, 1H), 3.24 (s, 7H), 1.98 (d, *J* = 6.8 Hz, 1H), 1.47 (d, *J* = 6.8 Hz, 6H), 1.31 (d, *J* = 7.0 Hz, 3H), 1.23 (s, 1H), 0.86 (dd, *J* = 19.1, 6.8 Hz, 6H).

**4-((*S*)-2-((*S*)-2-amino-3-methylbutanamido)propanamido)benzyl 4-(6-(((5,6-dichloro-1*H*-benzo[*d*]imidazol-2-yl)methyl)amino)-9-isopropyl-9*H*-purin-2-yl)piperazine-1-carboxylate (8)**

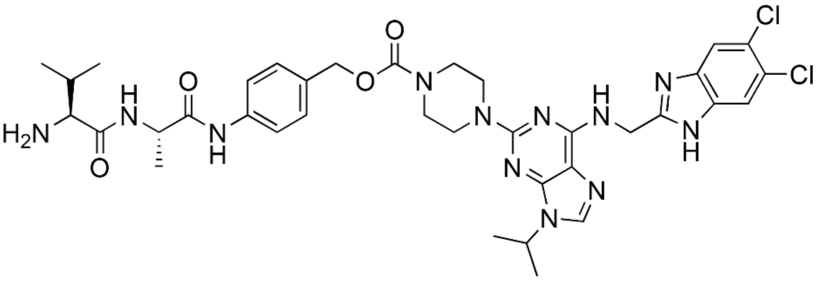

**7** (110 mg, 0.13 mmol) and $Pd(PPh_3)_4$ (9.40 mg, 8.00 µmol) were dissolved in anhydrous DCM (5 mL). Pyrrolidine (24.0 µL, 0.29 mmol) was dissolved in anhydrous DCM (2 mL) and added dropwise to the reaction mixture. The reaction was stirred at RT for 4 h before dilution with DCM (10 mL), washing with saturated ammonium chloride (10 mL), brine (10 mL) and drying over sodium sulfate. This compound was carried forward without further purification.

LC-MS (5-95% MeCN in $H_2O$, positive mode) Rt = 3.56 min, $[M+2H]^{2+}$ = 390.2

**4-((17*S*,20*S*)-1-(2,5-dioxo-2,5-dihydro-1*H*-pyrrol-1-yl)-17-isopropyl-20-methyl-15,18-dioxo-3,6,9,12-tetraoxa-16,19-diazahenicosan-21-amido)benzyl 4-(6-(((5,6-dichloro-1*H*-benzo[*d*]imidazol-2-yl)methyl)amino)-9-isopropyl-9*H*-purin-2-yl)piperazine-1-carboxylate (9, SS2009)**

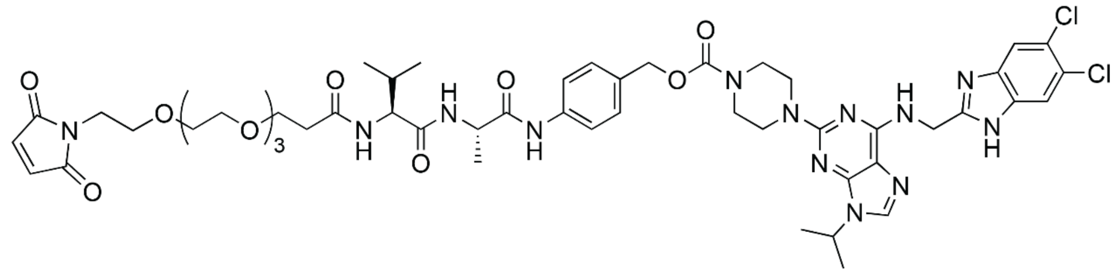

**8** (25.0 mg, 0.03 mmol) was dissolved in anhydrous DMF (2 mL). Mal-PEG4-NHS (17.7 mg, 0.04 mmol) was dissolved in anhydrous DMF (800 µL) and added to the solution dropwise. DIPEA (8.38 µL) was dissolved in anhydrous DMF (200 µL) and added dropwise to the reaction mixture. The reaction mixture was stirred at RT for 4 h, before concentration *in vacuo* and purification by high-performance liquid chromatography (30-95% MeCN (0.1% FA) in $H_2O$) to yield **SS2009** (18.8 mg, 53%) as a white solid.

LC-MS (5-95% MeCN in $H_2O$, negative mode) Rt = 4.00 min, $[M-H]^-$ = 1104.5

HRMS (TOF MS $ES^+$) Calc. for $C_{51}H_{66}N_{13}O_{11}Cl_2^+$ = 1106.4382, found $[M+H]^+$ = 1106. 4393

HPLC (30-95% % MeCN (0.1% FA) in $H_2O$, 220 nm) Rt = 4.36 min

#### Bioconjugations

##### Materials and methods

Conjugations were performed in Eppendorf Protein LoBind tubes at atmospheric pressure and temperatures stated, without shaking.

Antibodies (b12 and trastuzumab) were produced in house by ADC Therapeutics.

**Concentration determination**

Concentrations of antibodies and ADCs were determined by UV/Vis absorbance at 280 nm with a ThermoScientific NanoDrop One Microvolume UV/Vis spectrophotometer, with $\varepsilon_{280}$= 13.7 L $g^{-1}$ $cm^{-1}$ (preset IgG reference setting). MW and ε280 of each conjugate were calculated from those of the parent antibody and the DAR, as below.

MW of ADC = MW of mAb + (DAR x MW of payload)
EC280 of ADC = EC280 of mAb + (DAR x EC280 of payload)

B12
MW = 148170.43 Da
EC280 = 210800 $M^{-1}$ $cm^{-1}$

Trastuzumab
MW = 145455.49 Da
EC280 = 204540 $M^{-1}$ $cm^{-1}$

SS2009
MW (average) = 1107.1 Da
EC280 = 11642 $M^{-1}$ $cm^{-1}$

**Hydrophobic interaction chromatography (HIC)**

Antibody samples were diluted to 24 ug/mL using HIC loading buffer (3 M ammonium sulfate (AS) in 100 mM sodium phosphate (NaPi), pH 7.0) and PBS (pH 7.4) in a 1:2 ratio. The ThermoScientific Vanquish HPLC system was equipped with a TSKgel Butyl-NPR 2.5 µm, 4.6 mm x 35 mm column. 20 µL sample (24 ng/mL) was injected onto the column using analytical HIC buffer A (1.5 M AS in 50 mM NaPi buffer, pH 7) and analytical HIC buffer B (50 mM NaPi, 20% isopropyl alcohol (IPA), pH 7) over a 10 min 0-100% solvent B elution gradient with a flow rate of 1.2 mL/min. The autosampler temperature was set to 10 °C with the column oven maintained at 30 °C. UV (DAD) detection was obtained at 214, 280, 362 and 375 nm.

**Size exclusion chromatography (SEC)**

Samples were diluted to 1 mg/mL (with respect to antibody) using PBS buffer (pH 7.4). The ThermoScientific Vanquish HPLC system was equipped with an Acquity UPLC Protein BEH SEC 200 Å, 1.7 µm, 4.6 mm x 150 mm column with a MaxPeak Premier Protein SEC 250 Å, 2.5 µm, 4.6 mm x 30 mm guard column. 10 µL sample (1 mg/mL) was injected onto the column using 0.2 M potassium phosphate (15% IPA, 0.2 M KCl, pH 6.8) and an isocratic hold performed over 10 min, with a flow rate of 0.35 mL/min. The autosampler temperature was set to 10 °C with the column oven maintained at 30 °C. UV (DAD) detection was obtained at 214, 280, 362 and 375 nm.

**Polymeric reverse-phase chromatography (PLRP)**

Samples were diluted to 0.5 mg/mL (with respect to antibody) using PBS buffer (pH 7.4) and 10% (v/v) of a 50 mM dithiothreitol (DTT) stock solution, to a final volume of 100 µL. The mixture was reduced for 30 min at RT. The ThermoScientific Vanquish HPLC system was equipped with a ThermoScientific MAbPac Reversed Phase HPLC column (2.1mm x 50mm, 4 µm). 20 µL sample (0.5 mg/mL) was injected using PLRP mobile phase A (0.1% TFA in HPLC grade $H_2O$) and PLRP mobile phase B (0.1% TFA in HPLC grade MeCN) a flow rate of 0.3 mL/min. The autosampler temperature was set to 10 °C with the column oven maintained at 80 °C. UV (DAD) detection was obtained at 280, 330, 362 and 375 nm.

**Liquid chromatography mass spectroscopy (LC-MS)**

Samples (17 µL, 0.5 mg/mL in PBS pH 7.4) were incubated with PNGase F (1 µL, New England Biolabs, P0704S) for 3 h at 37 °C. DTT (2 µL, 50 mM) was added, and the mixture was incubated for 30 min at RT. Molecular masses were measured with an Agilent 6545XT AdvanceBio LC/Q-TOF system equipped with an Agilent 300SB-C3 guard. Samples (5 µL, 0.5 mg/mL) were injected onto the column using LC-MS solvent A (0.1 formic acid in HPLC grade $H_2O$) and LC-MS solvent B (0.1% formic acid in HPLC grade acetonitrile) with a flow rate of 0.5 mL/min and an oven temperature of 60 °C.

Q-TOF MS was acquired in a positive polarity mode coupled with a dual AJS ESI ion source. The source parameters were VCap of 4000 V, nozzle voltage of 2000 V, gas temperature of 350 °C, gas flow of 12 L/min, a skimmer of 65 V, a fragmentor of 180 V, and a nebulizer of 35 psig. The acquisition range was from 90 to 5000 m/z with a scan rate of 1.00 spectra/sec.

**Endotoxin testing**

Buffers, antibodies and ADCs were tested for endotoxin contamination using Charles River Endosafe LAL cartridges with a Charles River Endosafe Nexgen-PTS spectrophotometer according to the manufacturer's instructions. Samples were considered negative if endotoxin units (EU) per mg were below 1. Samples that failed the endotoxin tests were subject to further purification by protein A affinity purification (ThermoFisher Scientific, NAb™ Protein A Plus Spin Columns 1 mL, #89956) prior to conjugation.

### Compound characterization data

#### Compound 1

$^{1}H$ NMR

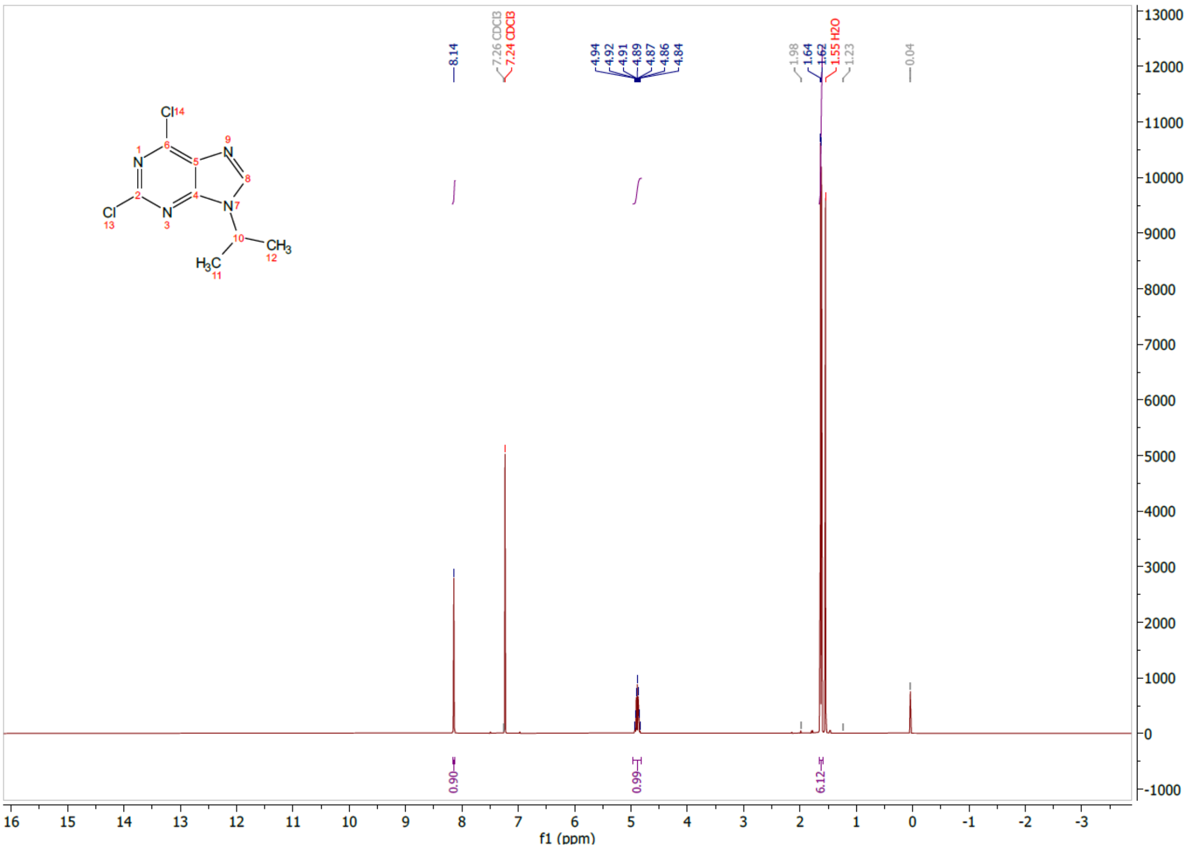

**Compound 2**

$^{1}H$ NMR

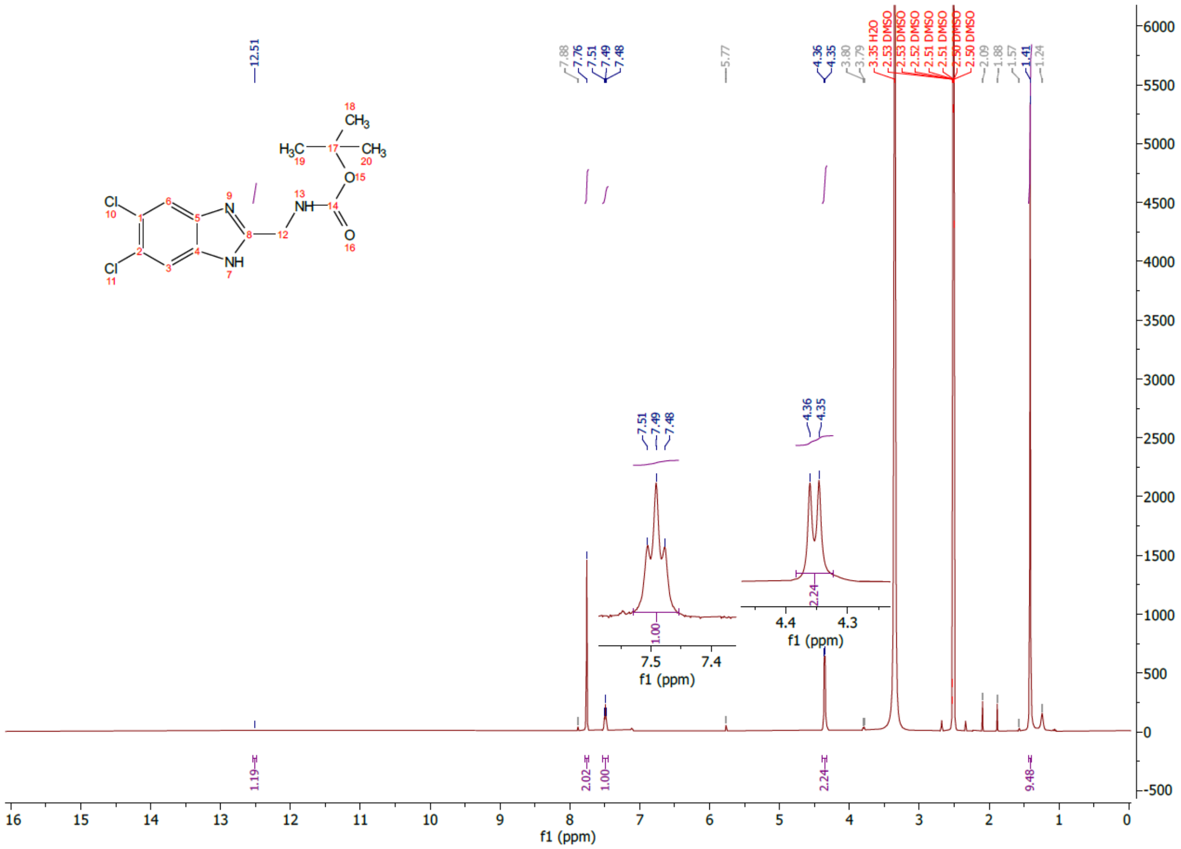

**Compound 3**

$^{1}$H NMR

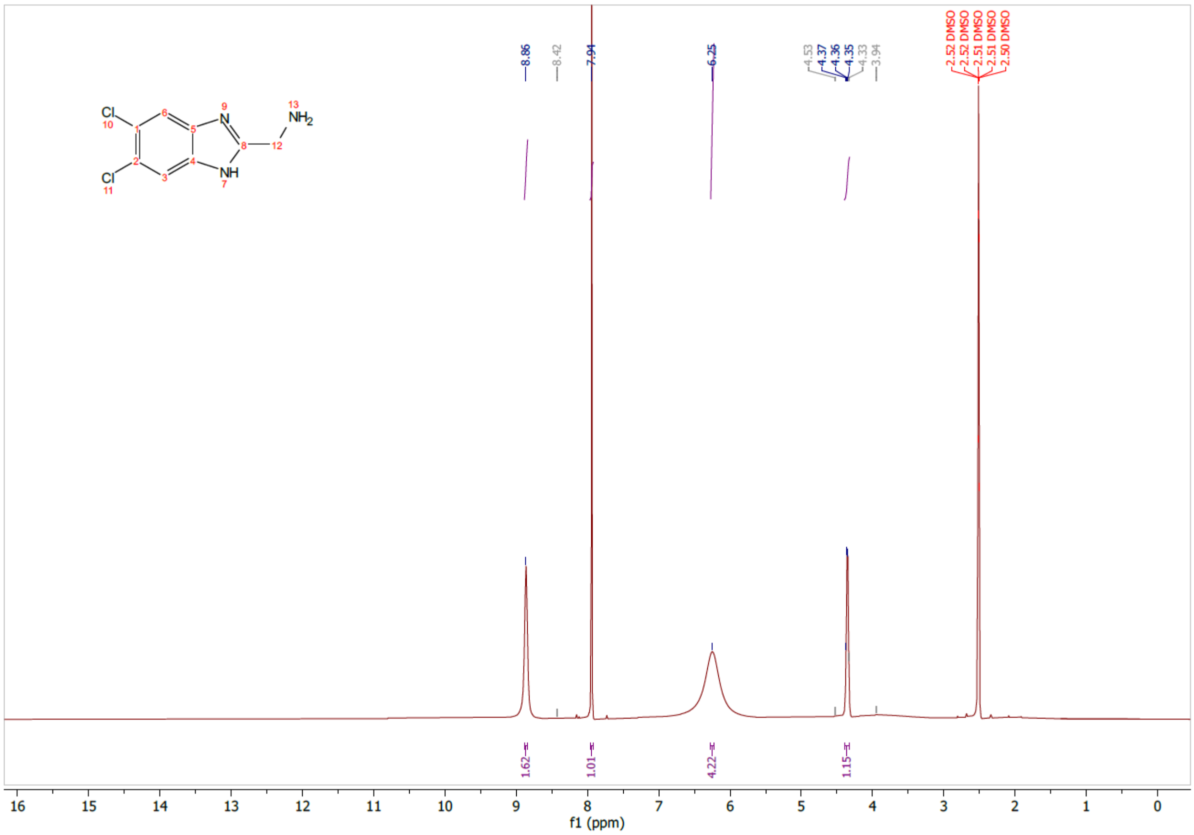

**Compound 5 (SAS1001)**

[1]H NMR

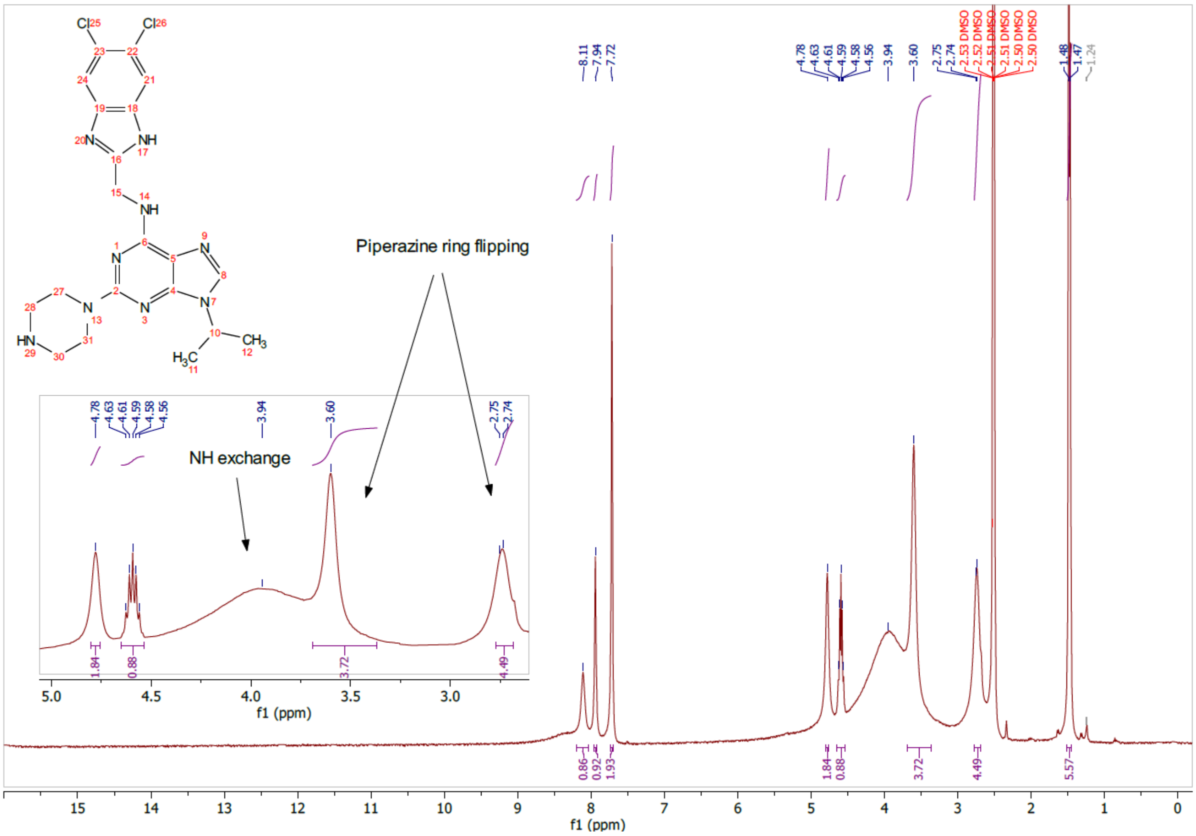


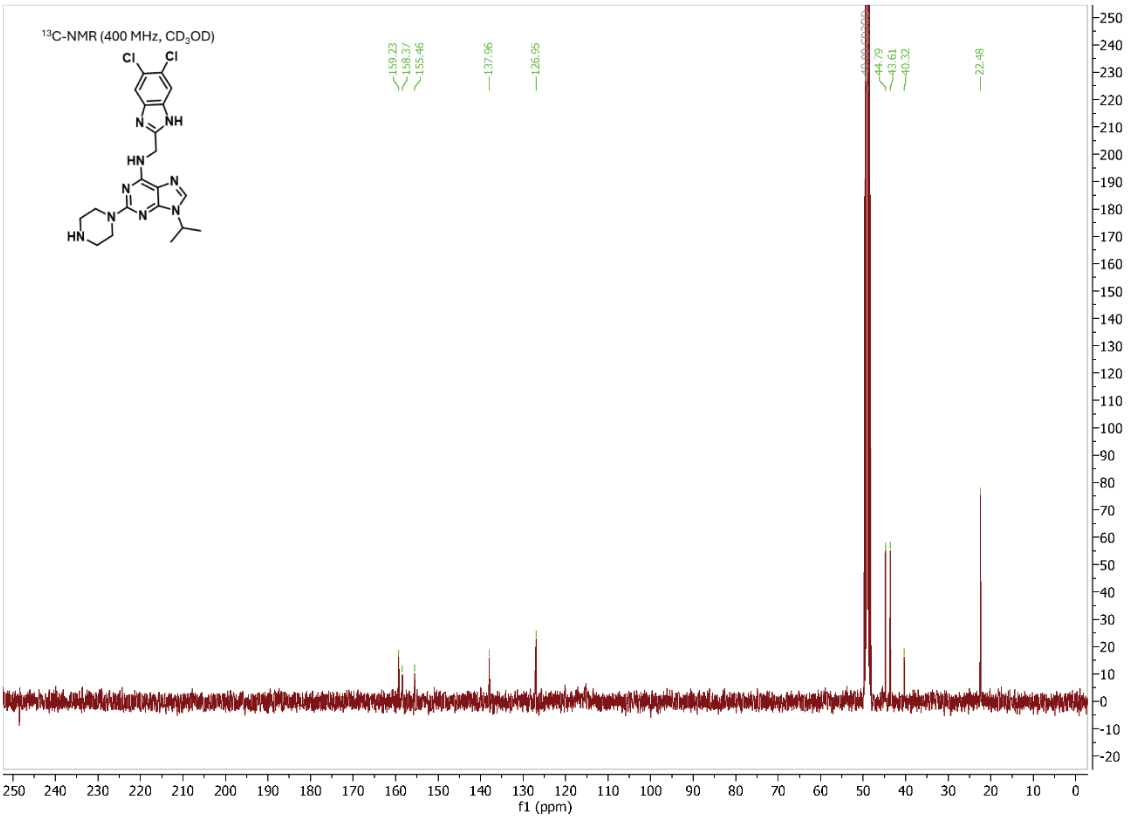

### HRMS

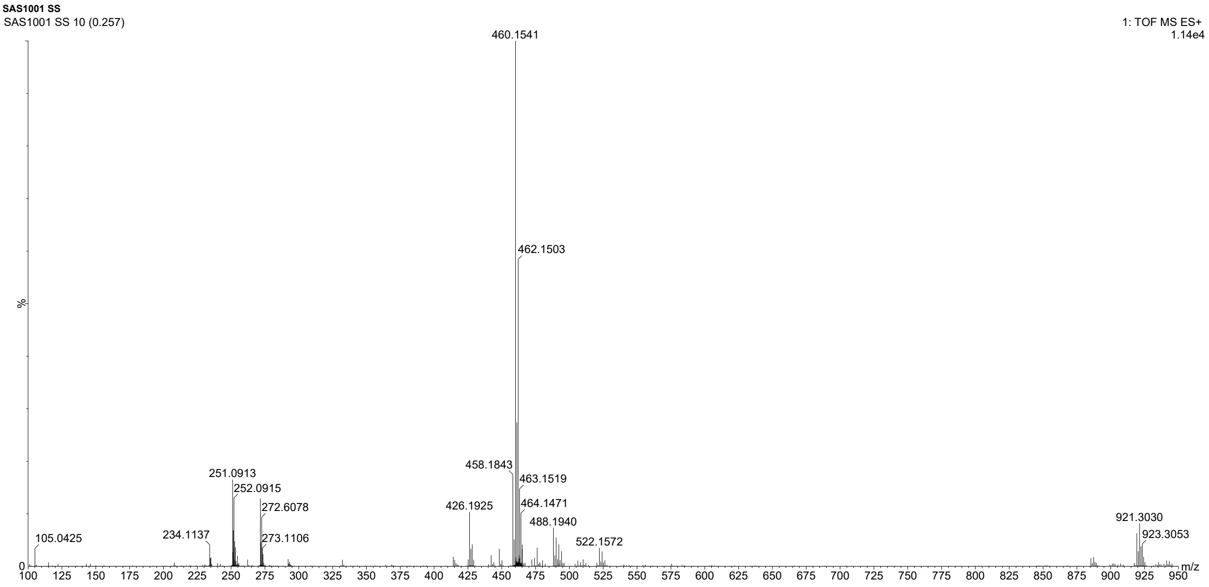

SAS1001 SS
SAS1001 SS 10 (0.257)
1: TOF MS ES+
1.14e4
460.1541
462.1503
458.1843
463.1519
464.1471
488.1940
522.1572
426.1925
251.0913
252.0915
272.6078
273.1106
234.1137
105.0425
921.3030
923.3053
%
m/z

**Compound 6**

$^1$H NMR

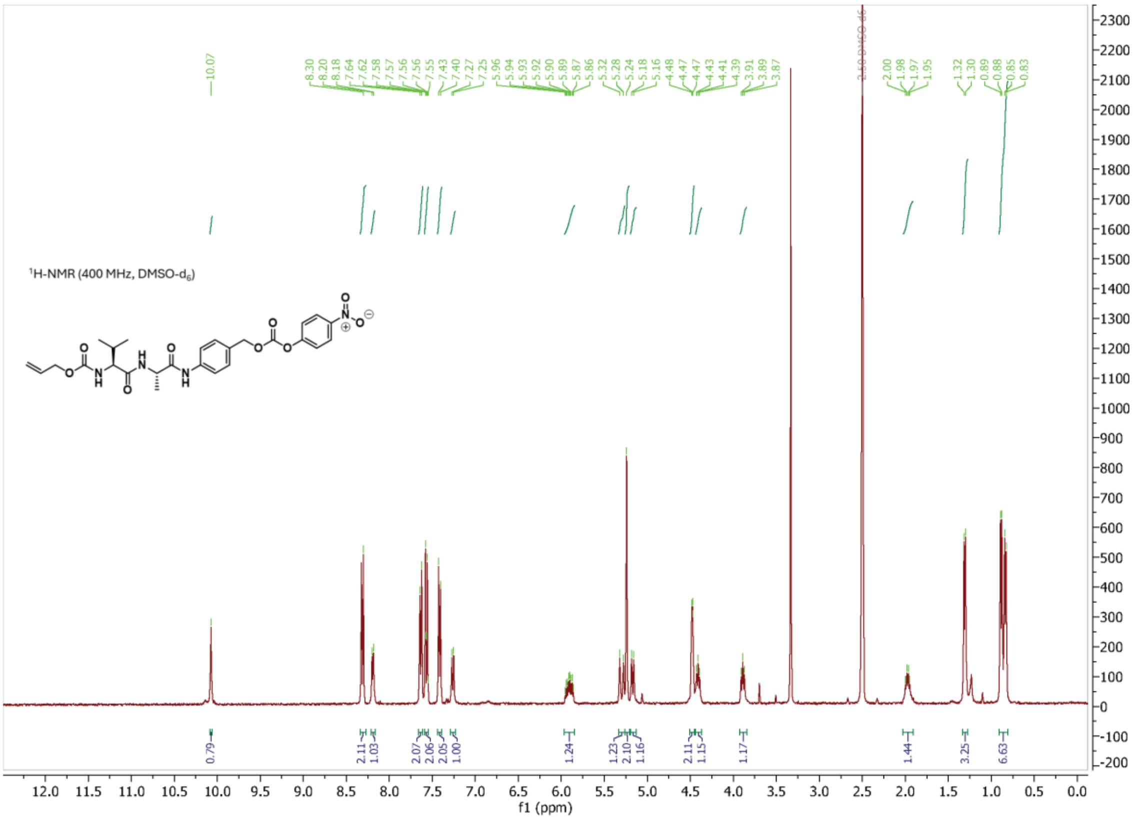

#### Compound 7

$^{1}H$ NMR

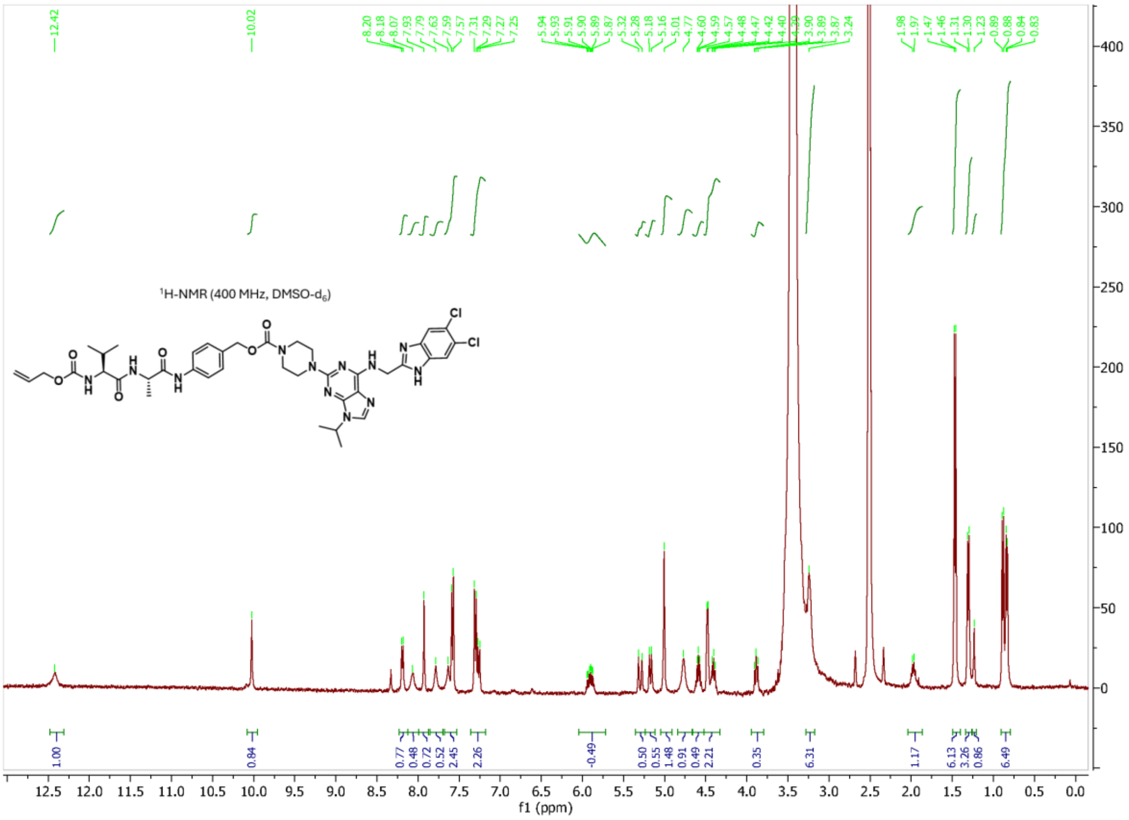

### Compound 9 (SS2009)

HRMS

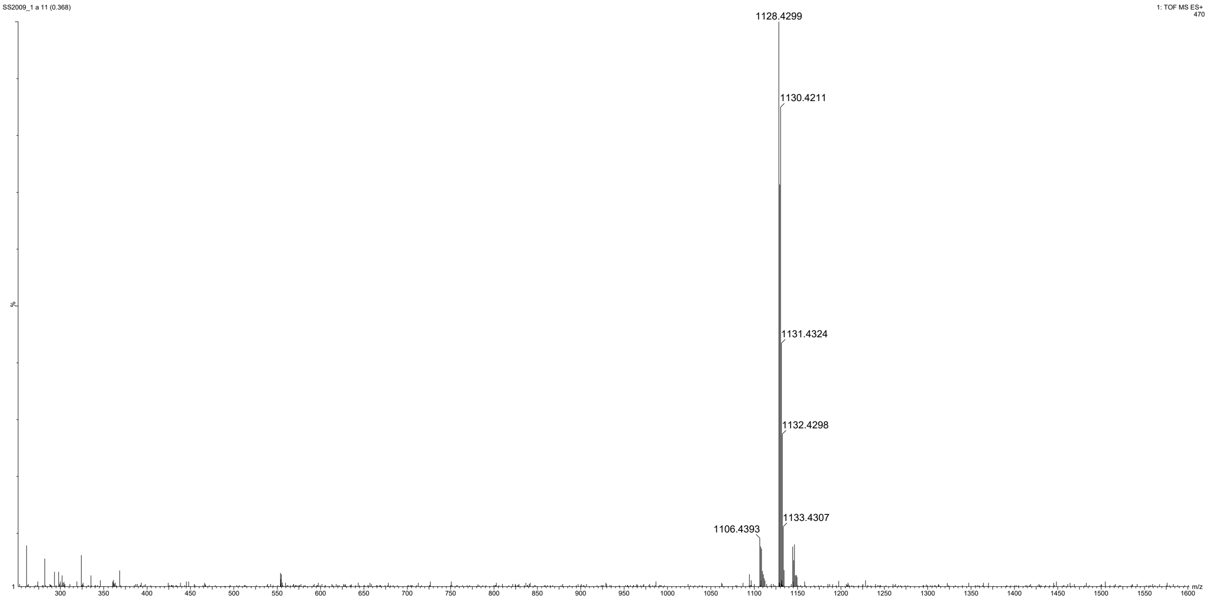


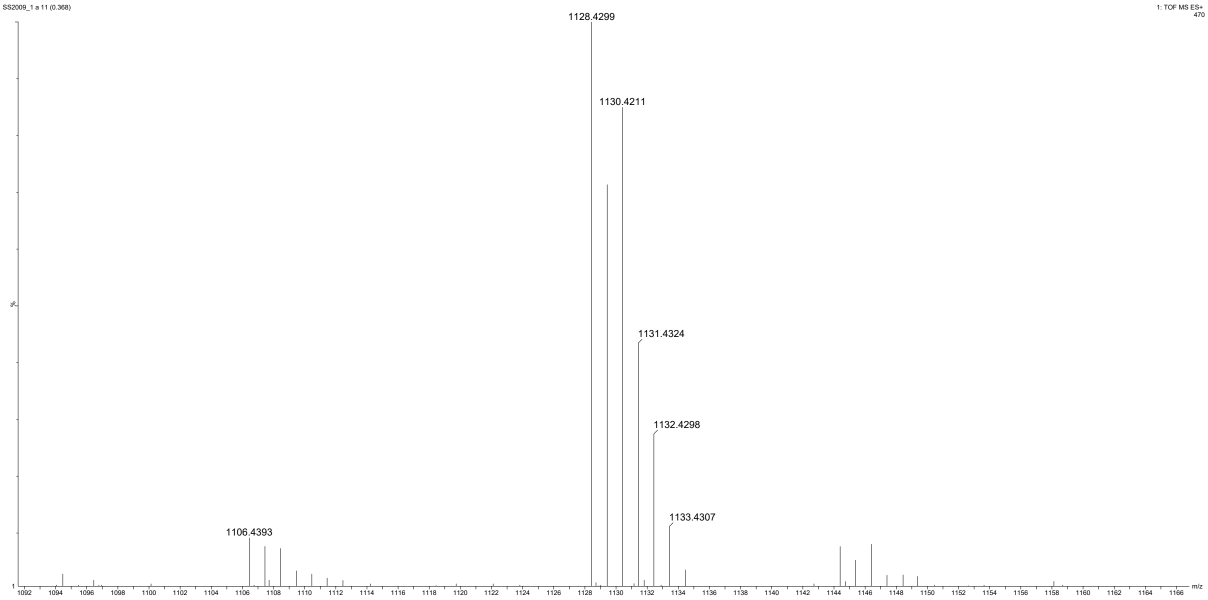

Analytical HPLC

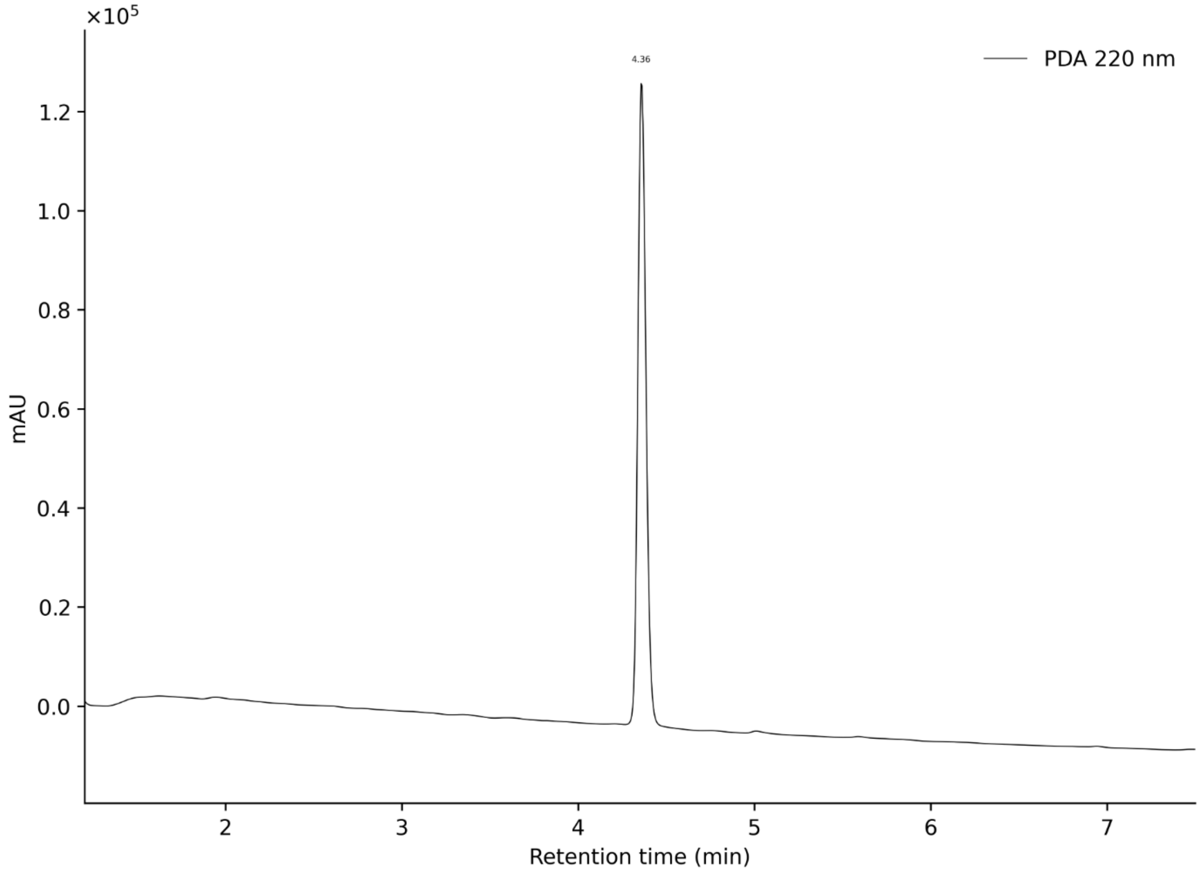

#### Original gels and blots

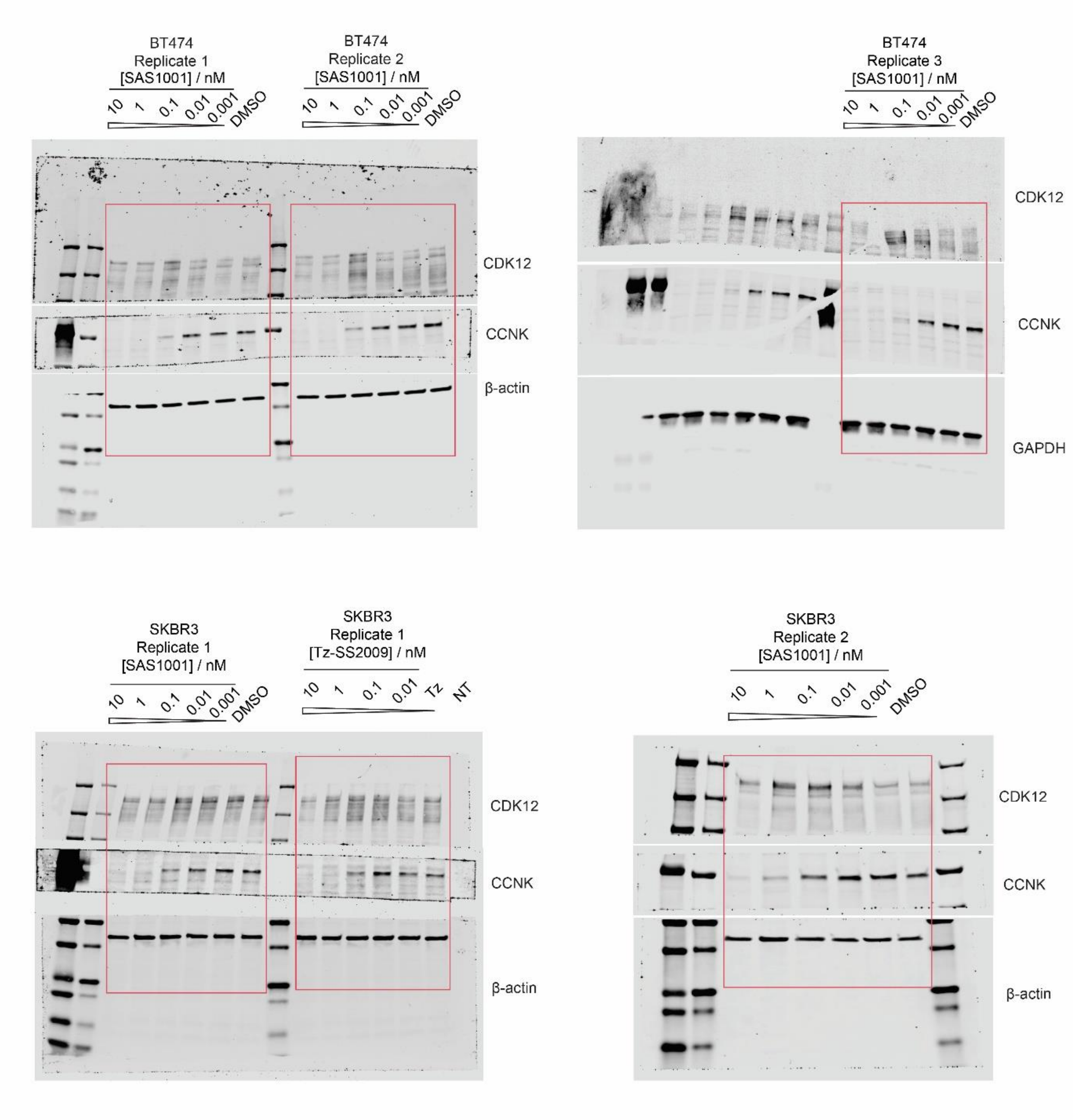

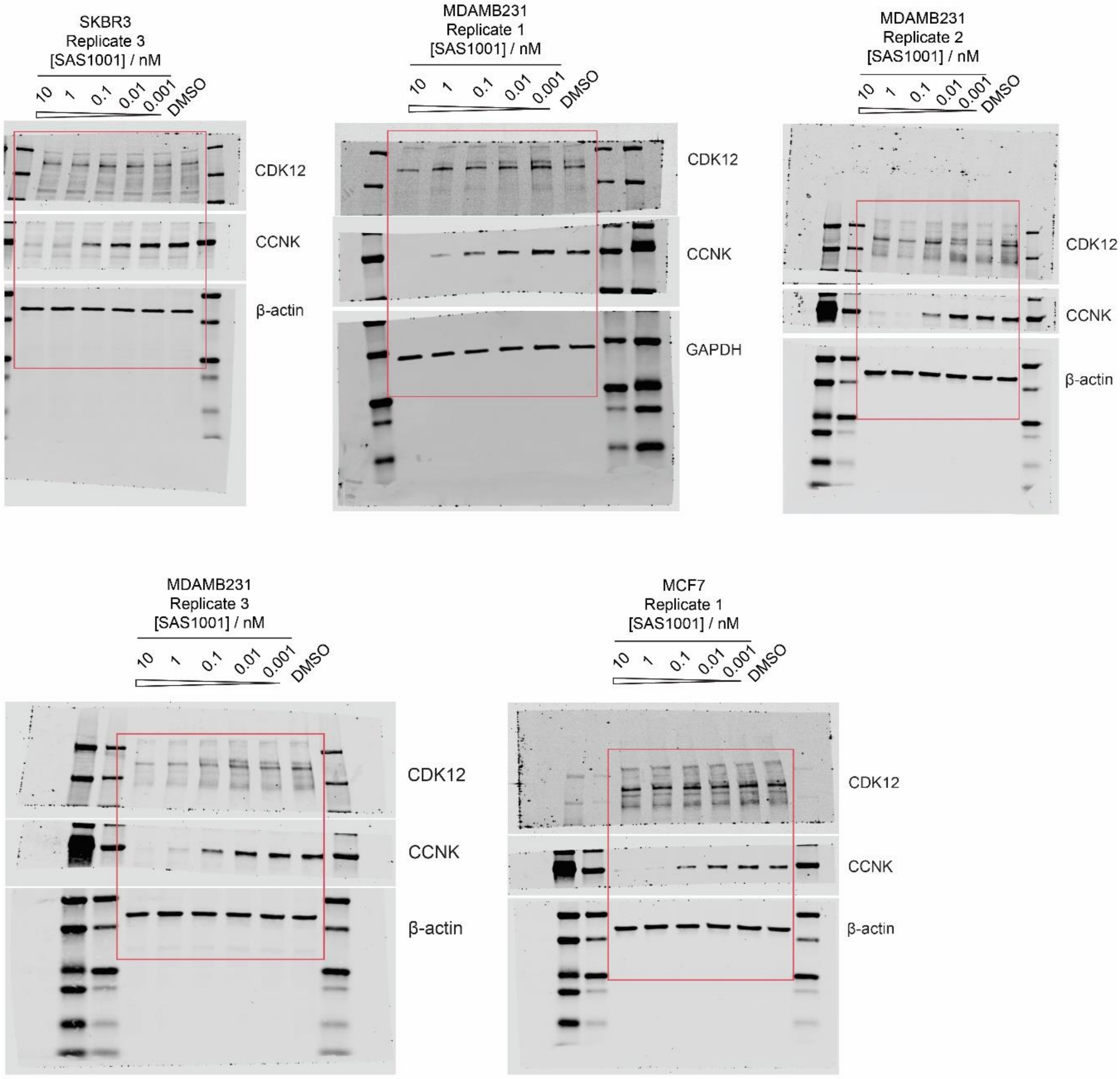
SKBR3
Replicate 3
[SAS1001] / nM
10
1
0.1
0.01
0.001
DMSO
CDK12
CCNK
β-actin
MDAMB231
Replicate 1
[SAS1001] / nM
10
1
0.1
0.01
0.001
DMSO
CDK12
CCNK
GAPDH
MDAMB231
Replicate 2
[SAS1001] / nM
10
1
0.1
0.01
0.001
DMSO
CDK12
CCNK
β-actin
MDAMB231
Replicate 3
[SAS1001] / nM
10
1
0.1
0.01
0.001
DMSO
CDK12
CCNK
β-actin
MCF7
Replicate 1
[SAS1001] / nM
10
1
0.1
0.01
0.001
DMSO
CDK12
CCNK
β-actin

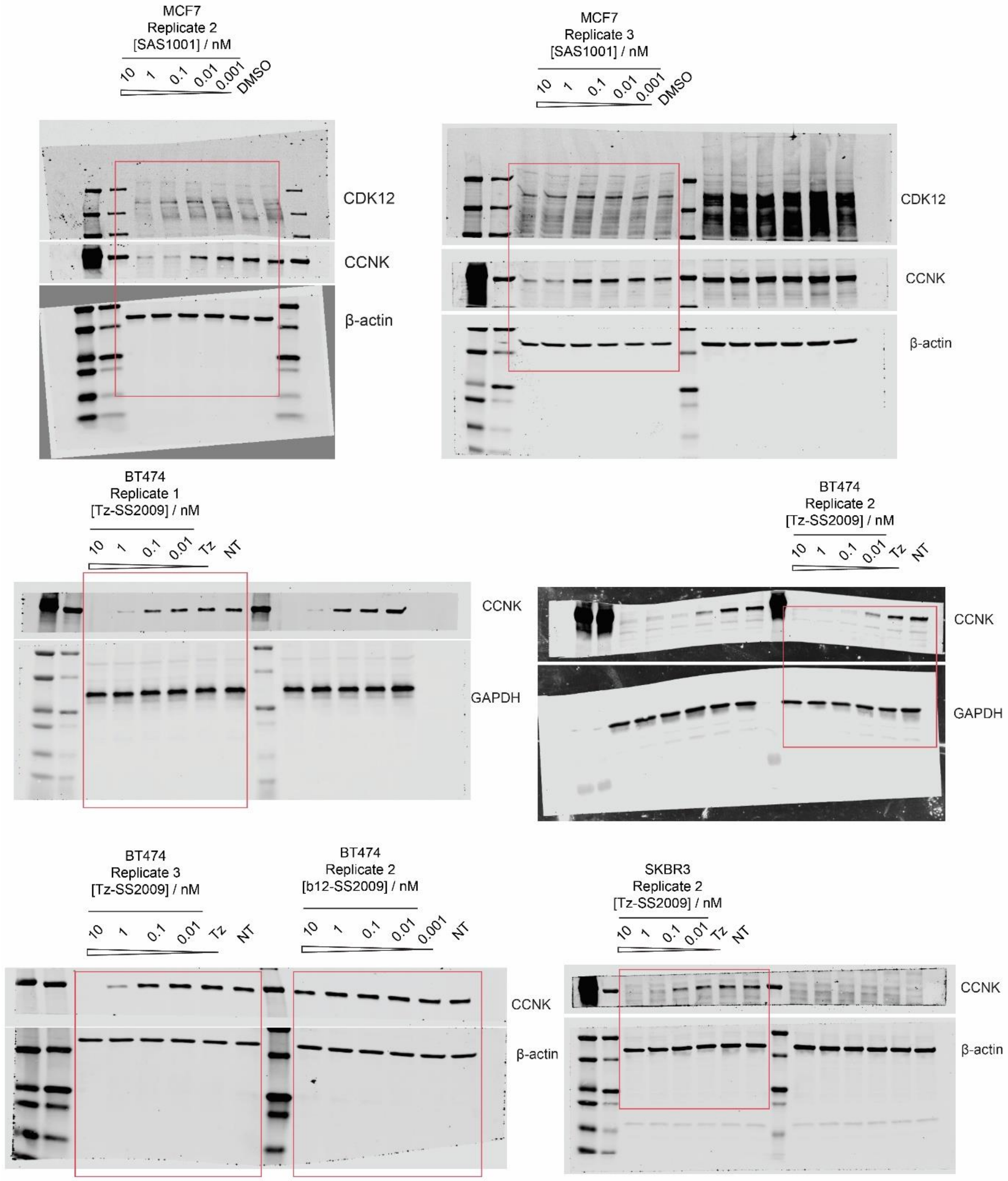
MCF7
Replicate 2
[SAS1001] / nM
10 1 0.1 0.01 0.001 DMSO
CDK12
CCNK
β-actin
MCF7
Replicate 3
[SAS1001] / nM
10 1 0.1 0.01 0.001 DMSO
CDK12
CCNK
β-actin
BT474
Replicate 1
[Tz-SS2009] / nM
10 1 0.1 0.01 Tz NT
CCNK
GAPDH
BT474
Replicate 2
[Tz-SS2009] / nM
10 1 0.1 0.01 Tz NT
CCNK
GAPDH
BT474
Replicate 3
[Tz-SS2009] / nM
10 1 0.1 0.01 Tz NT
BT474
Replicate 2
[b12-SS2009] / nM
10 1 0.1 0.01 0.001 NT
CCNK
β-actin
SKBR3
Replicate 2
[Tz-SS2009] / nM
10 1 0.1 0.01 Tz NT
CCNK
β-actin

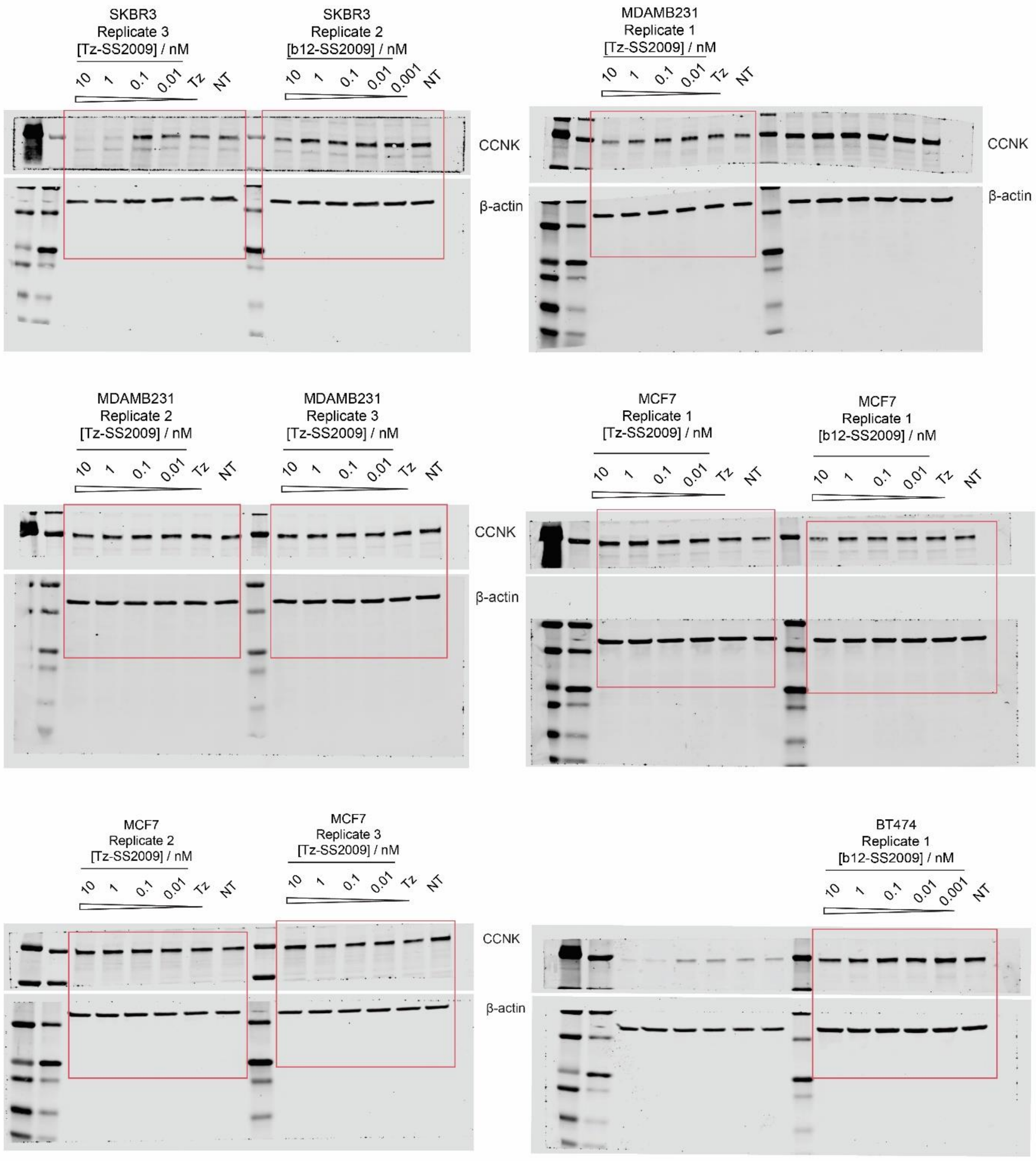
SKBR3
Replicate 3
[Tz-SS2009] / nM
10 1 0.1 0.01 Tz NT
SKBR3
Replicate 2
[b12-SS2009] / nM
10 1 0.1 0.01 0.001 NT
CCNK
β-actin
MDAMB231
Replicate 1
[Tz-SS2009] / nM
10 1 0.1 0.01 Tz NT
CCNK
β-actin
MDAMB231
Replicate 2
[Tz-SS2009] / nM
10 1 0.1 0.01 Tz NT
MDAMB231
Replicate 3
[Tz-SS2009] / nM
10 1 0.1 0.01 Tz NT
CCNK
β-actin
MCF7
Replicate 1
[Tz-SS2009] / nM
10 1 0.1 0.01 Tz NT
MCF7
Replicate 1
[b12-SS2009] / nM
10 1 0.1 0.01 Tz NT
MCF7
Replicate 2
[Tz-SS2009] / nM
10 1 0.1 0.01 Tz NT
MCF7
Replicate 3
[Tz-SS2009] / nM
10 1 0.1 0.01 Tz NT
CCNK
β-actin
BT474
Replicate 1
[b12-SS2009] / nM
10 1 0.1 0.01 0.001 NT

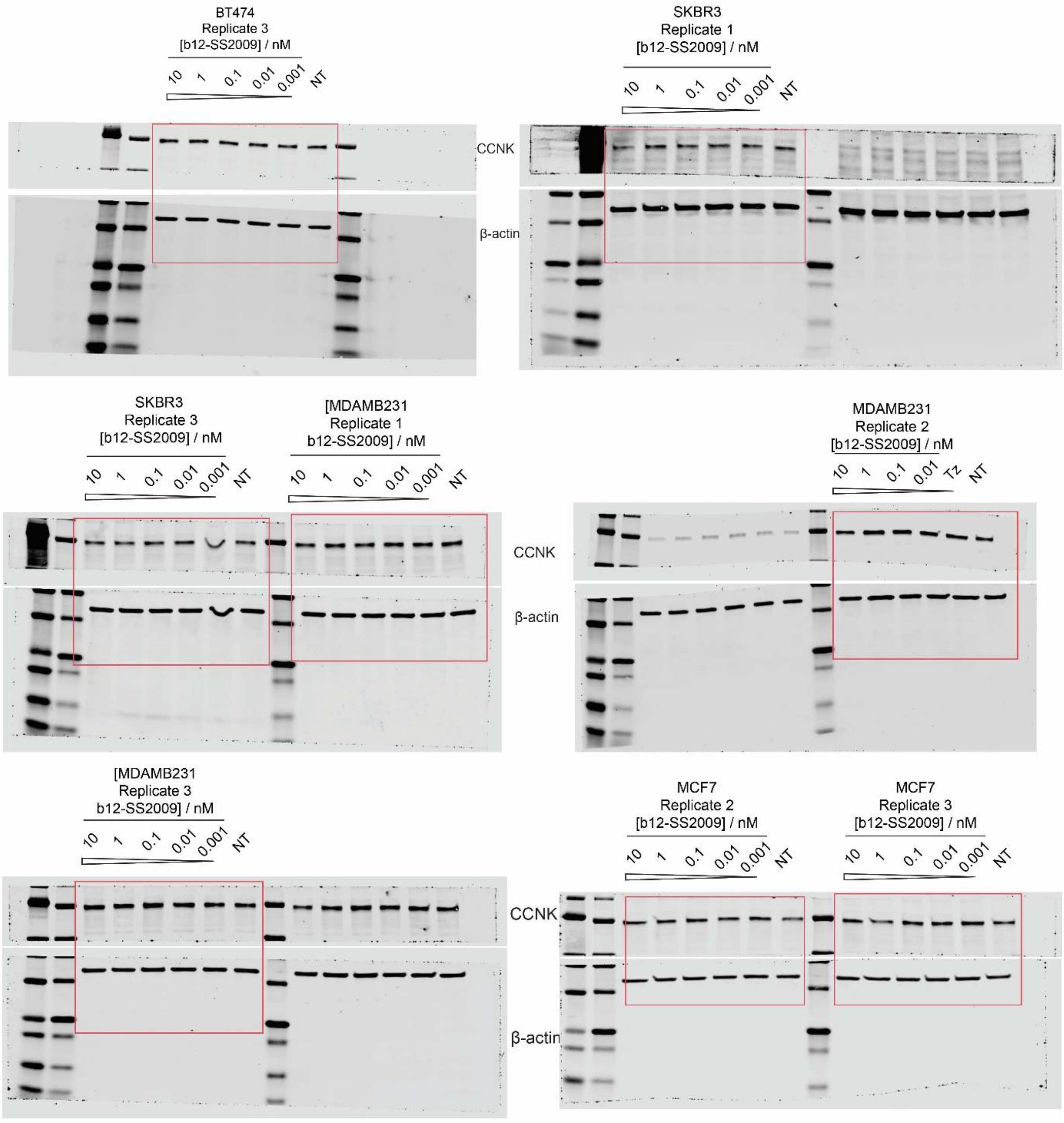

BT474
Replicate 3
[b12-SS2009] / nM
10
1
0.1
0.01
0.001
NT
CCNK
β-actin
SKBR3
Replicate 1
[b12-SS2009] / nM
10
1
0.1
0.01
0.001
NT
SKBR3
Replicate 3
[b12-SS2009] / nM
10
1
0.1
0.01
0.001
NT
[MDAMB231
Replicate 1
b12-SS2009] / nM
10
1
0.1
0.01
0.001
NT
CCNK
β-actin
MDAMB231
Replicate 2
[b12-SS2009] / nM
10
1
0.1
0.01
Tz
NT
[MDAMB231
Replicate 3
b12-SS2009] / nM
10
1
0.1
0.01
0.001
NT
CCNK
β-actin
MCF7
Replicate 2
[b12-SS2009] / nM
10
1
0.1
0.01
0.001
NT
MCF7
Replicate 3
[b12-SS2009] / nM
10
1
0.1
0.01
0.001
NT

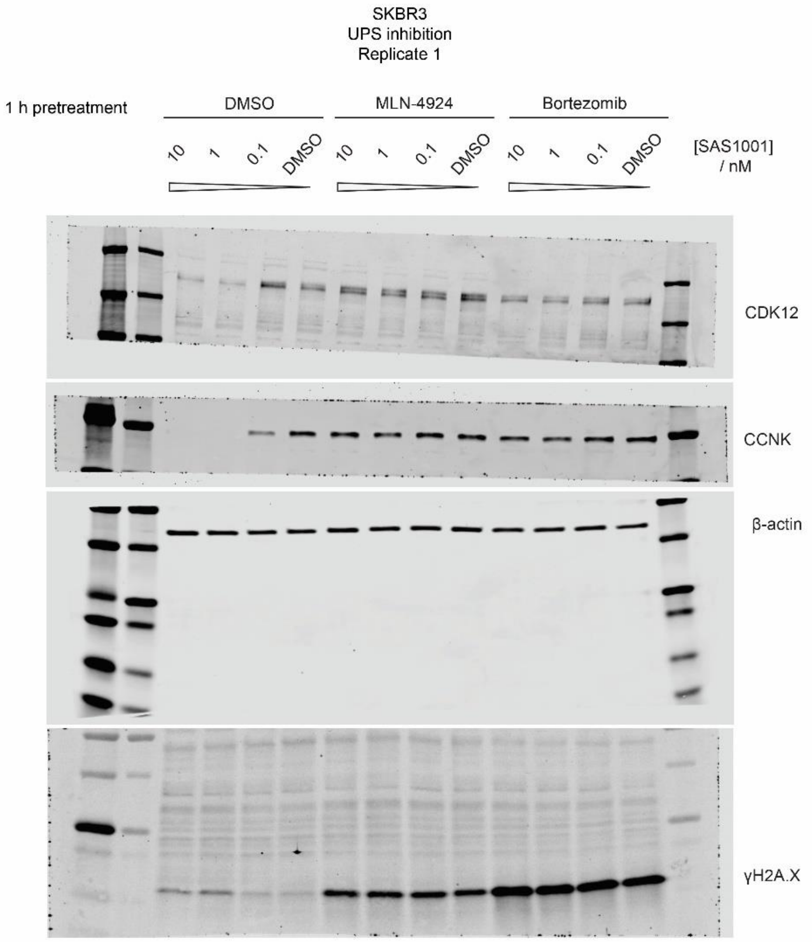
SKBR3
UPS inhibition
Replicate 1
1 h pretreatment
DMSO
MLN-4924
Bortezomib
10
1
0.1
DMSO
10
1
0.1
DMSO
10
1
0.1
DMSO
[SAS1001] / nM
CDK12
CCNK
β-actin
γH2A.X

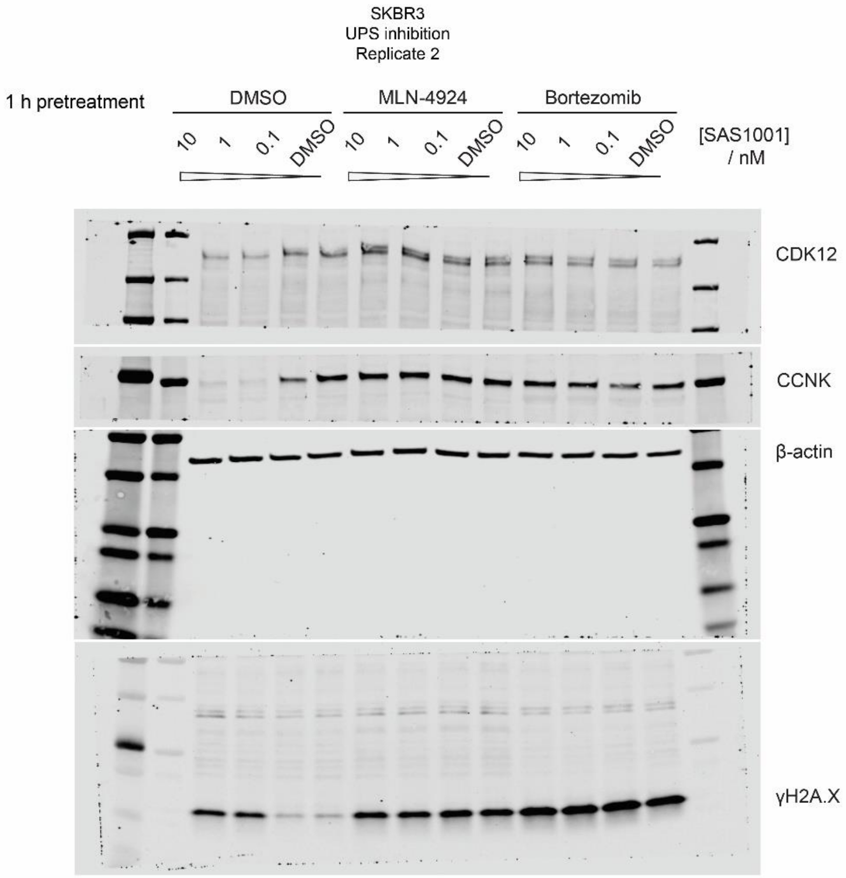
SKBR3
UPS inhibition
Replicate 2
1 h pretreatment
DMSO
MLN-4924
Bortezomib
10
1
0.1
DMSO
[SAS1001]
/ nM
CDK12
CCNK
β-actin
γH2A.X

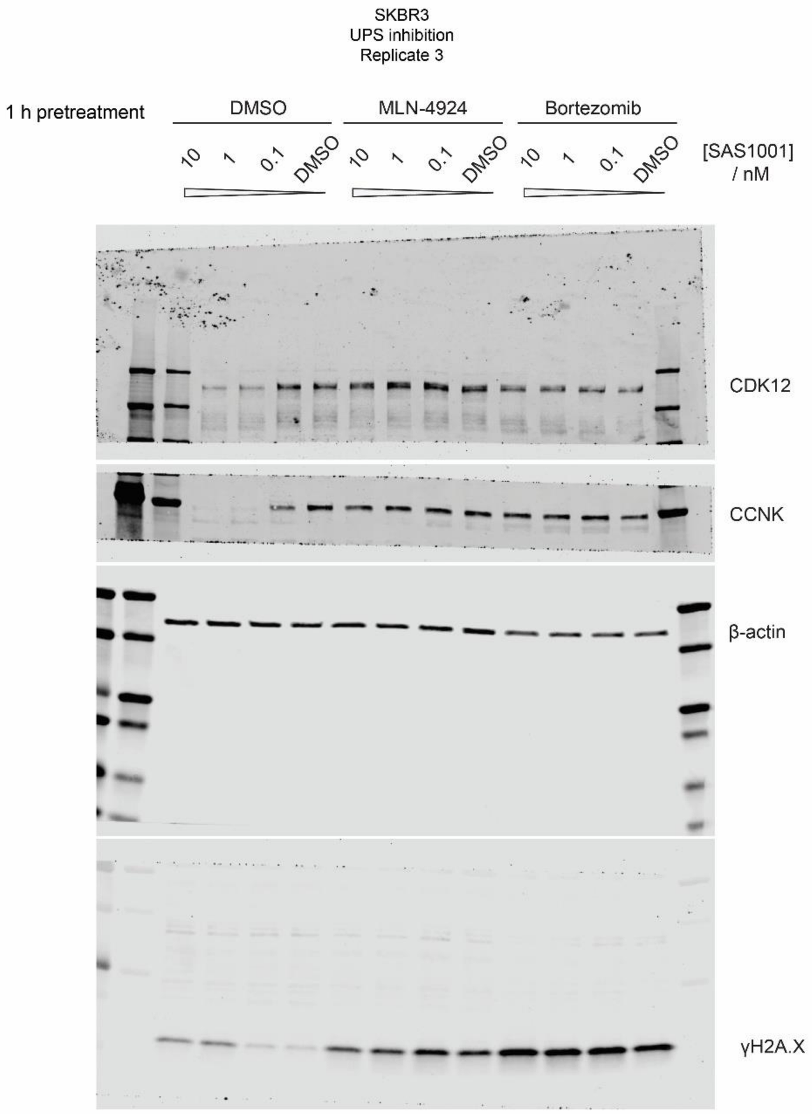
SKBR3
UPS inhibition
Replicate 3
1 h pretreatment
DMSO
MLN-4924
Bortezomib
10
1
0.1
DMSO
10
1
0.1
DMSO
10
1
0.1
DMSO
[SAS1001] / nM
CDK12
CCNK
β-actin
γH2A.X

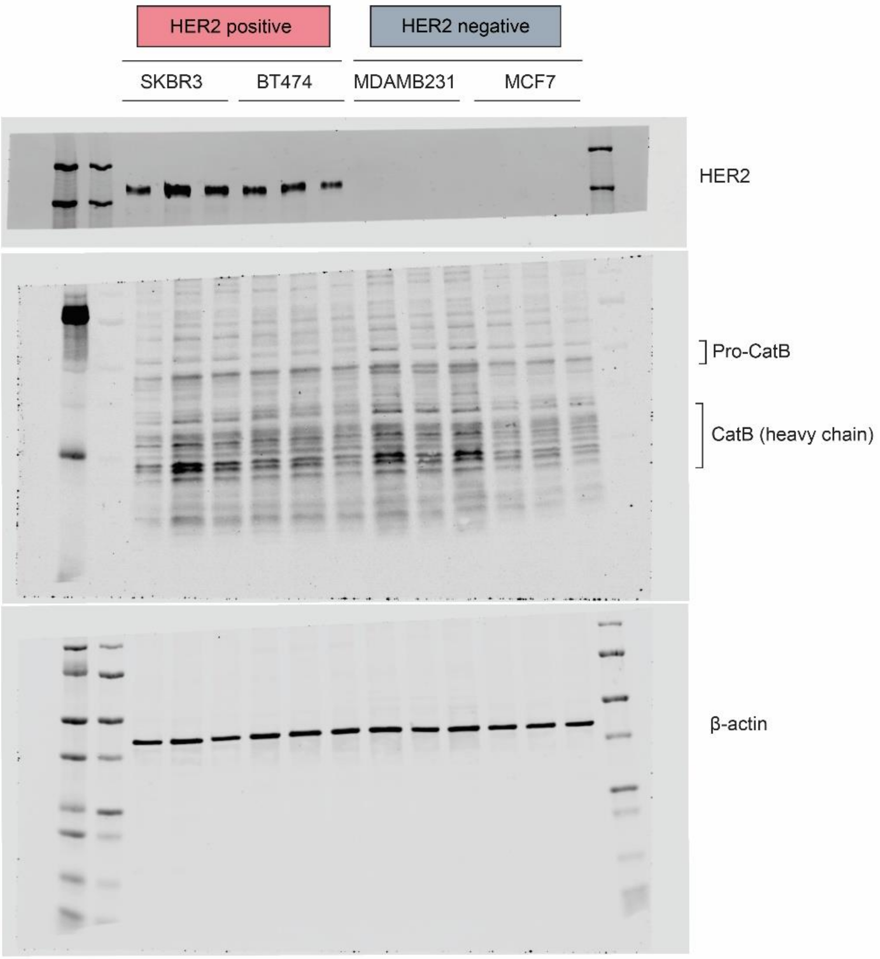
HER2 positive
HER2 negative
SKBR3
BT474
MDAMB231
MCF7
HER2
Pro-CatB
CatB (heavy chain)
β-actin
